# Integrating pharmacogenomics and cheminformatics with diverse disease phenotypes for cell type-guided drug discovery

**DOI:** 10.1101/2022.12.21.521456

**Authors:** Arda Halu, Julius L. Decano, Joan Matamalas, Mary Whelan, Takaharu Asano, Namitra Kalicharran, Sasha A. Singh, Joseph Loscalzo, Masanori Aikawa

## Abstract

Large-scale pharmacogenomic resources, such as the Connectivity Map (CMap), have greatly assisted computational drug discovery. However, despite their widespread use, CMap-based methods have thus far been agnostic to the biological activity of drugs, as well as to the genomic effects of drugs in multiple disease contexts. Here, we present a network-based statistical approach, Pathopticon, that uses CMap to build cell type-specific gene-drug perturbation networks and integrates these networks with cheminformatic data and diverse disease phenotypes for a nested prioritization of cell lines and drugs. Pathopticon demonstrates a better prediction performance than solely cheminformatic measures and state-of-the-art methods that rely exclusively on pharmacogenomic data. Top predictions made by Pathopticon have high chemical structural diversity, suggesting their potential for building compound libraries. In a proof-of-concept application on vein graft disease, we validate the predicted drugs *in vitro* and demonstrate that Pathopticon helps pinpoint the shared intermediate phenotypes targeted by each prediction. Overall, our analytical framework integrating pharmacogenomics and cheminformatics provides a feasible blueprint for a cell typespecific drug discovery and repositioning platform with broad implications for the efficiency and success of drug development.

## Introduction

Systems pharmacology has helped transcend the prevailing one drug-one target paradigm in drug discovery by analyzing complex networks of interactions between diseases, drugs, and targets to achieve an organism-wide view of drug action (*1–3*). Network pharmacological methods often use global drug-disease (*4, 5*) and drug-target (*6, 7*) networks as the basis upon which to predict new drug-target interactions and explore drug combinations. Large reference databases of molecular responses to systematic chemical, genetic, and disease perturbations, popularized by the Connectivity Map (CMap) (*8*), have proven to be an effective resource when used in conjunction with network-based drug discovery methods. In brief, CMap works by matching the transcriptional “signature” of an input set of genes (e.g., differentially expressed genes in a disease) with that of a perturbation (e.g., small molecule drugs), identifying, after a series of statistical steps, the perturbational signatures that best mimic or reverse the input signature (*9*). By allowing the rapid generation of testable hypotheses in a cell type- and input context-dependent manner, CMap-based network pharmacological approaches have helped expedite all aspects of drug discovery from drug repositioning (*10–13*) to classifying mechanisms of action of uncharacterized small molecules (*14, 15*) to identifying drug combinations (*16*) and side effects (*17*). Costeffective transcriptome profiling assays have further accelerated these efforts, scaling up the size of these resources to hundreds of thousands of gene expression signatures spanning tens of thousands of small molecules and nearly a hundred cell lines, as in the case of the Library of Integrated Network-Based Cellular Signatures (LINCS)-CMap (*18*). More than a decade after its inception, CMap is now established as a computational drug discovery framework, with many applications and methods (*9, 19, 20*) built upon its foundation.

Despite their widespread use and clear utility in drug discovery, CMap-based network pharmacology studies have had numerous limitations that can be summarized into three main themes: The first limitation reflects the challenges involving CMap data and the consequent underutilization of CMap as a resource for methods development. CMap data can be considered as a tensor (high-dimensional vector) (*21*) in five dimensions, consisting of genes, perturbations, cell lines, time points, and doses, typically with varying numbers of experiments in each dimension. This heterogeneity and multidimensionality of the CMap data present challenges in reliably defining cell typespecific gene-perturbation networks. In addition, despite its superior coverage, and partly due to its younger age, the more recent LINCS-CMap dataset is still underutilized compared to the extended version of the initial CMap release (CMap Build 2), which remains the prevalent CMap resource. As a result, few efforts exist that generate cell type-specific gene-perturbation networks from LINCS-CMap data at a large scale as a resource to facilitate network-based pharmacological approaches. The second limitation stems from the focus of CMap-based methods on a single input disease context. CMap-based approaches have typically focused on directly matching disease-associated gene signatures with genome-wide transcriptional responses to perturbations (*9*), ignoring the larger network of gene-perturbation interactions within multiple disease contexts, Given that diseases and drugs with similar molecular and clinical characteristics are likely to share phenotypes (*22*), these approaches potentially miss opportunities to identify polypharmacological effects and off-target disease pathways. The third limitation is CMap’s predominant use as a stand-alone resource. It has been suggested that integrating pharmacogenomic data such as CMap with cheminformatic data (e.g., bioactivity and chemical structure data) may be beneficial for drug discovery, but such efforts have so far been confined to quantifying the correlations of transcriptomic response with bioactivity and chemical structure (*23*). It remains to be investigated as to whether or not integrating the two sources of information directly benefits drug prioritization. While cheminformatic tools are very useful in their own right in drug discovery given their provision of structure-activity relationships, the cell type-specific extension of such tools using large-scale gene profiling experiments such as CMap has been noted as an unexplored future direction (*24*).

To tackle the above challenges, we have developed a computational framework, Pathopticon **(Figure 1)**. As the first component of Pathopticon, we propose a genecentric approach, named <u>QU</u>antile-based <u>I</u>nstance <u>Z</u>-score <u>C</u>onsensus (QUIZ-C). QUIZ-C uses LINCS-CMap data to build cell type-specific gene-perturbation networks for 60 cell lines, identifying the consistent and statistically significant relationships between genes and perturbations. The second component of Pathopticon, the <u>PA</u>thophenotypic <u>CO</u>ngruity <u>S</u>core (PACOS), measures the agreement between input and perturbation signatures within a global network of diverse disease phenotypes. After combining PACOS with pharmacological activity data, Pathopticon performs a nested prioritization that identifies the most suitable cell lines for an input gene signature, followed by the perturbations whose transcriptomic response best aligns with that of the input signature within each cell line. It surpasses current methods in terms of predictive performance and offers mechanistic clues into the shared intermediate phenotypes targeted by the predicted drugs. As such, Pathopticon offers the potential to be a feasible blueprint for a cell type-specific drug discovery and repositioning platform that integrates pharmacogenomics and cheminformatics. The Pathopticon algorithm and web app are available at https://github.com/r-duh/Pathopticon.

**Figure 1:**
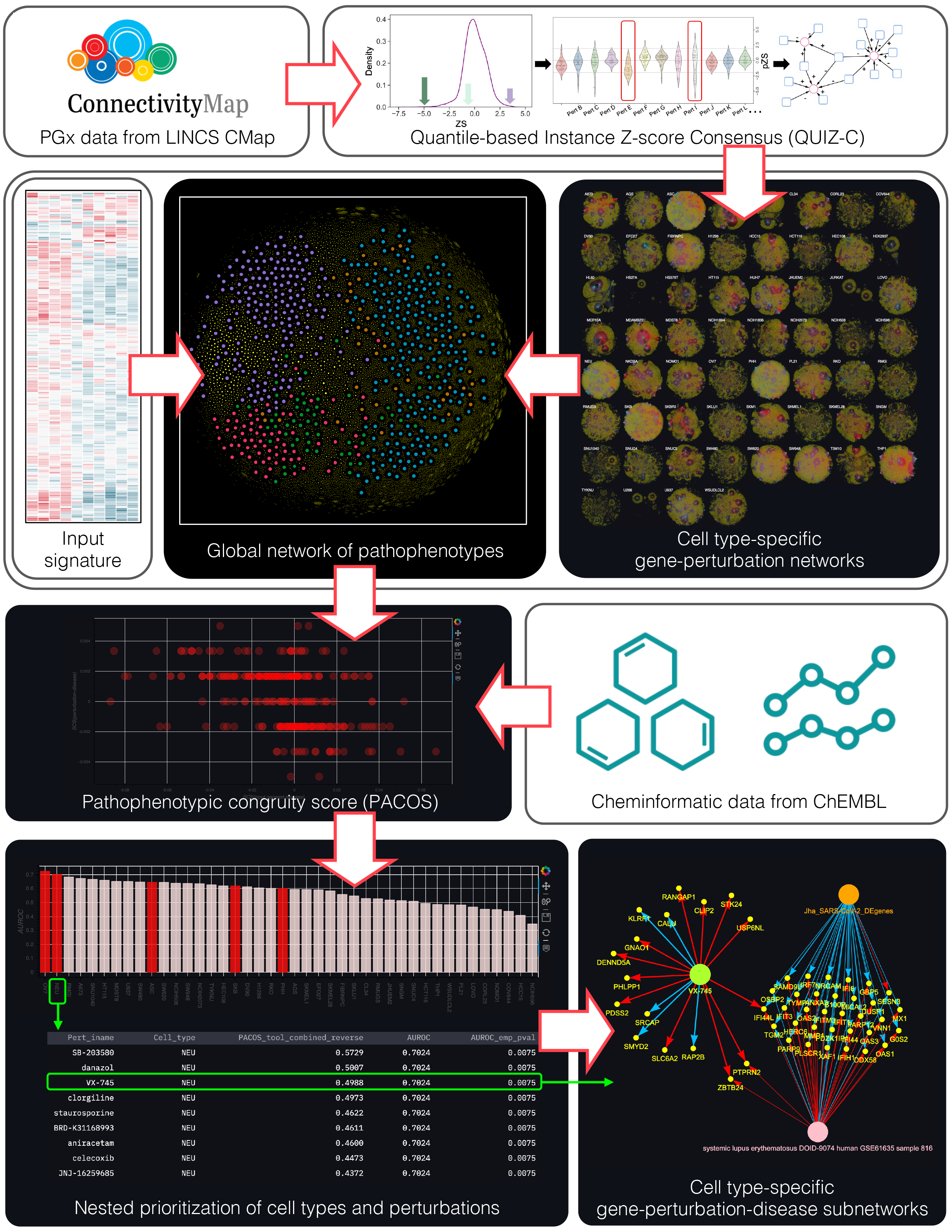
Overview of the Pathopticon framework.

## Results

### A gene-centric approach to building cell type-specific gene-perturbation networks

The recent large-scale CMap powered by the L1000 assay (referred to as LINCS-CMap or CMap 2.0) (*18*) contains gene expression measurements across the transcriptome in response to thousands of small molecule perturbations. These measurements were taken in a wide array of cell lines and over varying numbers of experiments (“instances”) encompassing different doses, time points, and replicate numbers. As a feature of this multidimensionality and heterogeneity, the LINCS-CMap data offer unique opportunities and challenges in building global, cell type-specific gene-perturbation networks for computational drug repositioning and discovery. However, efforts aimed at building these types of networks, while addressing the potential data biases adequately, remain limited. Notably, two recent approaches that aim to identify differential gene-perturbation pairs, namely the Moderated Z-score (MODZ) (*18*) and the Characteristic Direction (CD) measure (*25, 26*), have taken data heterogeneity into account to achieve a weighted consensus among experimental replicates of perturbations in LINCS-CMap. As such, they can be regarded as “perturbation-centric” approaches that can be extended to build cell type-specific gene-perturbation networks.

To address the challenges in reliably defining cell type-specific gene-perturbation networks while exploiting the full extent of the next generation CMap, we used LINCS-CMap Level 4 data **(Supplementary Methods)**, which retains the information on individual perturbation instances without collapsing them to a single value per perturbation **(Figure 2A)**. We devised a gene-centric network-building strategy, which we call <u>QU</u>antile-based <u>I</u>nstance <u>Z</u>-score <u>C</u>onsensus (QUIZ-C), that identifies the perturbations that (i) differentially and (ii) consistently modulate the expression of each gene **(Methods)**. Briefly, for each gene, we assign a “perturbation z-score,” *pZS*, to each perturbation instance against the background of all perturbation instances on that gene **(Figure 2B)**. The majority of perturbations in LINCS-CMap have multiple experimental instances **(Supplementary Figure 1)**, which permits variability between the *pZS* values of a perturbation due to different doses, time points, or experimental replicates. To prevent spurious gene-perturbation associations and capture high-confidence gene-perturbation pairs, we required a sufficient number of differential instances of a perturbation to view it as significantly affecting a gene. To determine this number, we defined a flexible consensus threshold based on the *pZS* distributions **(Figure 2C)**. Perturbations that passed this flexible consensus threshold were considered differentially affecting the expression of the gene in question **(Figure 2C-D)**. Repeating this procedure for all genes and perturbations in every cell line and aggregating all the differential gene-perturbation links **(Figure 2D)**, we built global cell type-specific gene-perturbation networks **(Supplementary Figure 2, Supplementary Table 1)**. To enable a direct comparison with our cell type-specific gene-perturbation networks (henceforth, referred to as QUIZ-C networks) in our downstream analyses, we also built cell type-specific gene-perturbation networks using the MODZ and CD approaches **(Supplementary Methods)**.

**Figure 2:**
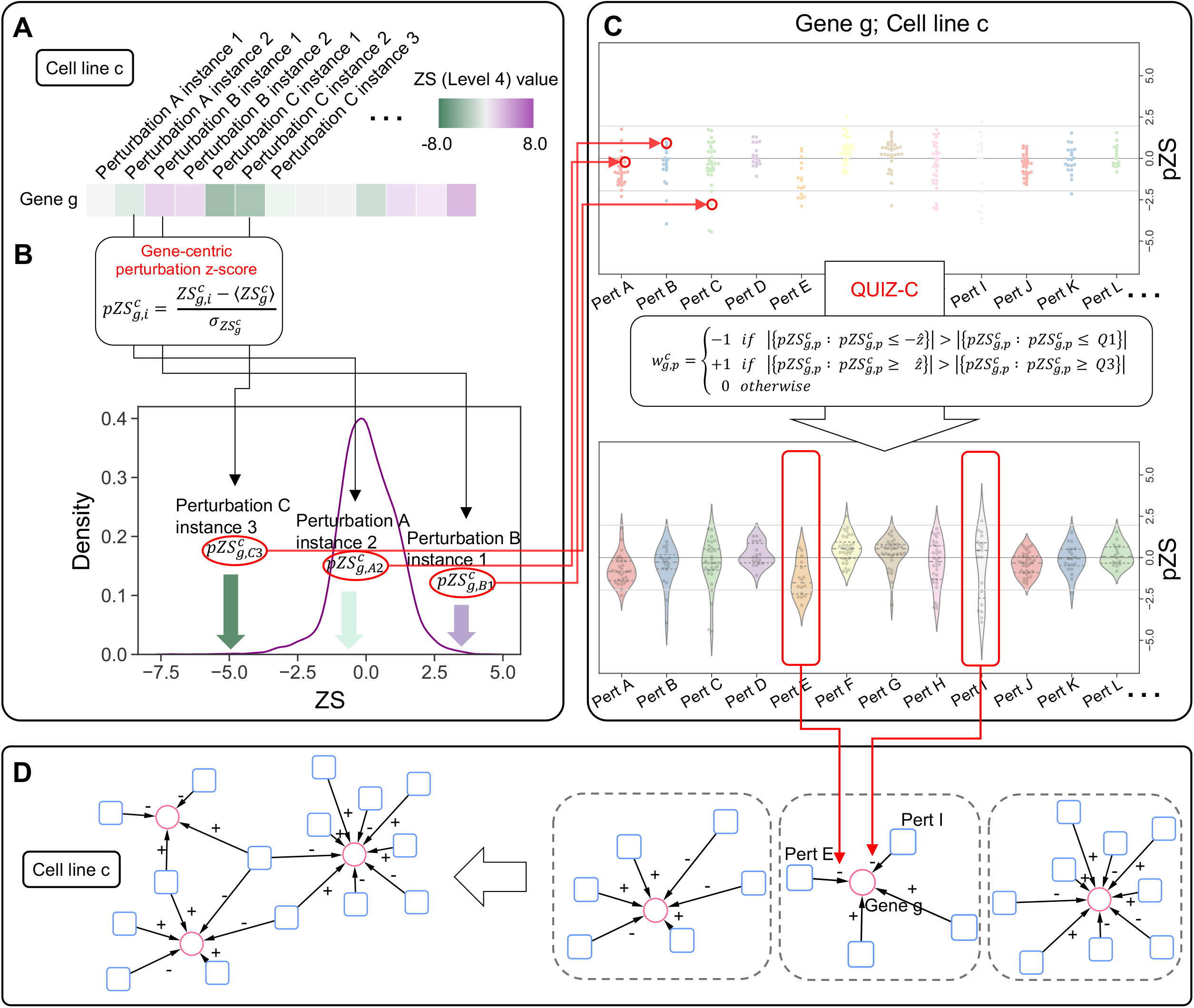
Overview of the QUIZ-C algorithm. **(A)** For every gene *g*, LINCS-CMap Level 4 data contains information on the expression value of all perturbation instances *i* tested in cell line *c*, denoted by 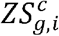. **(B)** We calculate the gene-centric perturbation z-score for each instance, denoted by 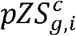, by comparing 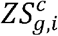 to the mean 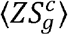 and standard deviation 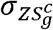 of all instances for gene *g* and cell line *c.* **(C)** For each gene and cell line, we select perturbations that pass a consensus threshold based on the 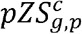 distributions of all of their instances. Q1 and Q3 indicate the first and third quartiles of 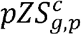. **(D)** We build cell type-specific gene-perturbation networks by repeating this procedure for all genes and perturbations in each cell line.

### QUIZ-C networks reflect the biological diversity of LINCS-CMap cell lines

QUIZ-C networks include the majority of the small-molecule perturbations tested on the CMap cell lines, with 34 out of 60 cell type-specific networks having over 80% of the perturbations tested in each respective cell line **(Figure 3A)**. We observed that the largest hub perturbations (i.e., perturbations modulating the expression of a large number of genes) are generally cell type-specific and are not shared by many cell lines, whereas perturbations with fewer targets can be present in a large number of cell lines (Pearson’s r = 0.013) **(Figure 3B)**. By contrast, genes targeted by many drugs were observed in a large number of cell lines, while genes targeted by fewer drugs were observed in fewer cell lines (Pearson’s r = 0.57) **(Figure 3C)**. These findings on hub drugs and targets **(Figures 3B and 3C)** were recapitulated in CD and MODZ networks **(Supplementary Figure 3)**, indicating that this feature is independent of the specific network-building approach and, rather, reflects the underlying biological differences between distinct cell lines and their transcriptional responses to small-molecule perturbations. In terms of individual gene-perturbation interactions, we found little overlap between pairs of cell types **(Figure 3D)**. While MODZ and CD networks also have little edge overlap between cell types **(Supplementary Figure 4A-B)**, their edge overlap is generally higher compared to QUIZ-C **(Supplementary Figure 4C)**. The lower edge overlap of QUIZ-C networks cannot simply be attributed to a lower network density (i.e., the number of existing edges out of all possible edges), as MODZ networks are the densest of the three sets of networks **(Supplementary Figure 4D)**, whereas CD networks have the highest edge overlap. In terms of the direction of effect (i.e., up- or down-regulation of the gene by the perturbation) of the overlapping edges across cell lines, we observed a high level of agreement between cell lines in QUIZ-C networks, which was supported by MODZ networks, and, to some degree, by CD networks **(Supplementary Figure 4E)**. This observation suggests that gene-perturbation pairs that are conserved across cell lines tend to have the same directional effect.

**Figure 3:**
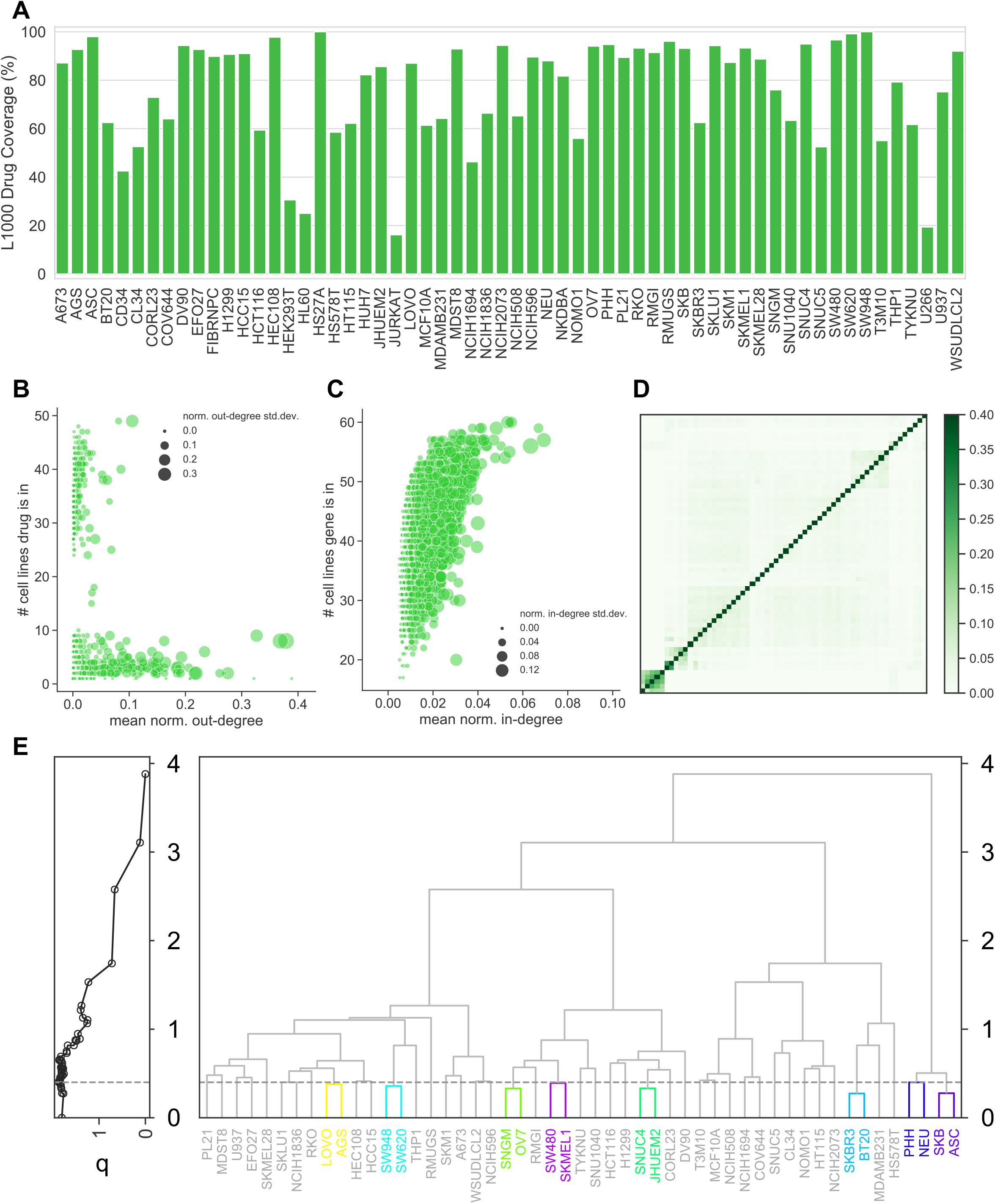
Properties of QUIZ-C networks. **(A)** The proportion of LINCS-CMap drugs tested in each cell line that was in the QUIZ-C network of that cell line. **(B)** The number of cell type-specific QUIZ-C networks a given drug appears in, as a function of the mean normalized out-degree of each drug across all QUIZ-C networks. Every circle represents a drug, and circle size is proportional to the standard deviation of the normalized out-degree. **(C)** The number of cell type-specific QUIZ-C networks a given gene appears in, as a function of the mean normalized in-degree of each gene across all QUIZ-C networks. Every circle represents a gene, and the circle size is proportional to the standard deviation of the normalized in-degree. **(D)** Hierarchically clustered heatmap qualitatively showing the edge overlap between QUIZ-C networks. Each row and column correspond to a cell line. The colors correspond to the Jaccard index of edge overlap. **(E)** Reducibility of QUIZ-C networks. Left panel shows the relative entropy quality function *q*(·) of each hierarchical clustering branch, with the maximum value of *q*(·) corresponding to the optimal configuration of the aggregation of layers. Right panel shows the hierarchical clustering dendrogram and the optimal clustering threshold. Cell lines belonging to the same cluster are in the same color, and clusters consisting of only one cell line are shown in light grey.

To supplement our findings on the structural uniqueness of QUIZ-C networks, we used a recent method that leverages information theoretic measures to quantify the degree of information redundancy (or “reducibility”) in multilayer networks **(Supplementary Methods)** (*27*). QUIZ-C, MODZ, and CD networks can each be represented as a multilayer network in which every layer is a cell type-specific network. Within this framework, QUIZ-C networks were highly non-redundant with a reducibility metric *χ* of 0.16 **(Figure 3E)**, whereas MODZ and CD were highly reducible with reducibility values of 0.92 and 0.74, respectively **(Supplementary Figures 5A and B)**. This finding suggests that, while most cell lines in MODZ and CD can be grouped together in terms of their topological similarity, the same cannot be said about QUIZ-C networks. Given the genetic and ontological diversity of the cell lines involved, which we have verified by quantifying their single tandem repeat (STR) profile similarity **(Supplementary Figure 6)** and Cell Line Ontology (CLO) similarity **(Supplementary Figure 7) (Supplementary Methods)**, this finding suggests that QUIZ-C networks remain sensitive to and thereby reflect differences among cell lines, a property that has important implications in disease-specific (rather than endophenotype-specific) drug development.

Overall, these results suggest that QUIZ-C networks have sufficient representation of drugs tested in CMap, are distinct in terms of their strongest perturbations, capture nonoverlapping sets of perturbation-gene pairs, and contain non-redundant information between cell types. These findings support the cell type-specificity of the QUIZ-C networks and suggest that they can serve as a good substrate on which to carry out *in silico* drug prediction.

### Drug repurposing and discovery potential of QUIZ-C networks

Using the Drug Repurposing Hub (DRH) (*28*), we inquired about the mechanism of action and clinical phase of drugs in LINCS-CMap-derived networks **(Supplementary Methods)**. Nearly 60% of drugs in most QUIZ-C networks had a known mechanism of action **(Figure 4A)**, indicating that ~60% of the drugs in our cell type-specific gene-perturbation networks have repurposing potential, while the rest are potentially novel therapeutic agents. These percentages were similar in MODZ and CD networks **(Supplementary Figure 8)**. In terms of clinical development, we found a balanced representation of experimental drugs (drugs in the preclinical phase), investigational drugs (drugs in clinical development phases 1 through 3), and approved (launched) drugs **(Figure 4B)**. This distribution was also observed in MODZ and CD networks **(Supplementary Figure 9A-B)**, suggesting that CMap-based gene-perturbation networks offer a combination of drug repurposing and novel drug discovery opportunities with drugs represented across all stages of drug development.

**Figure 4:**
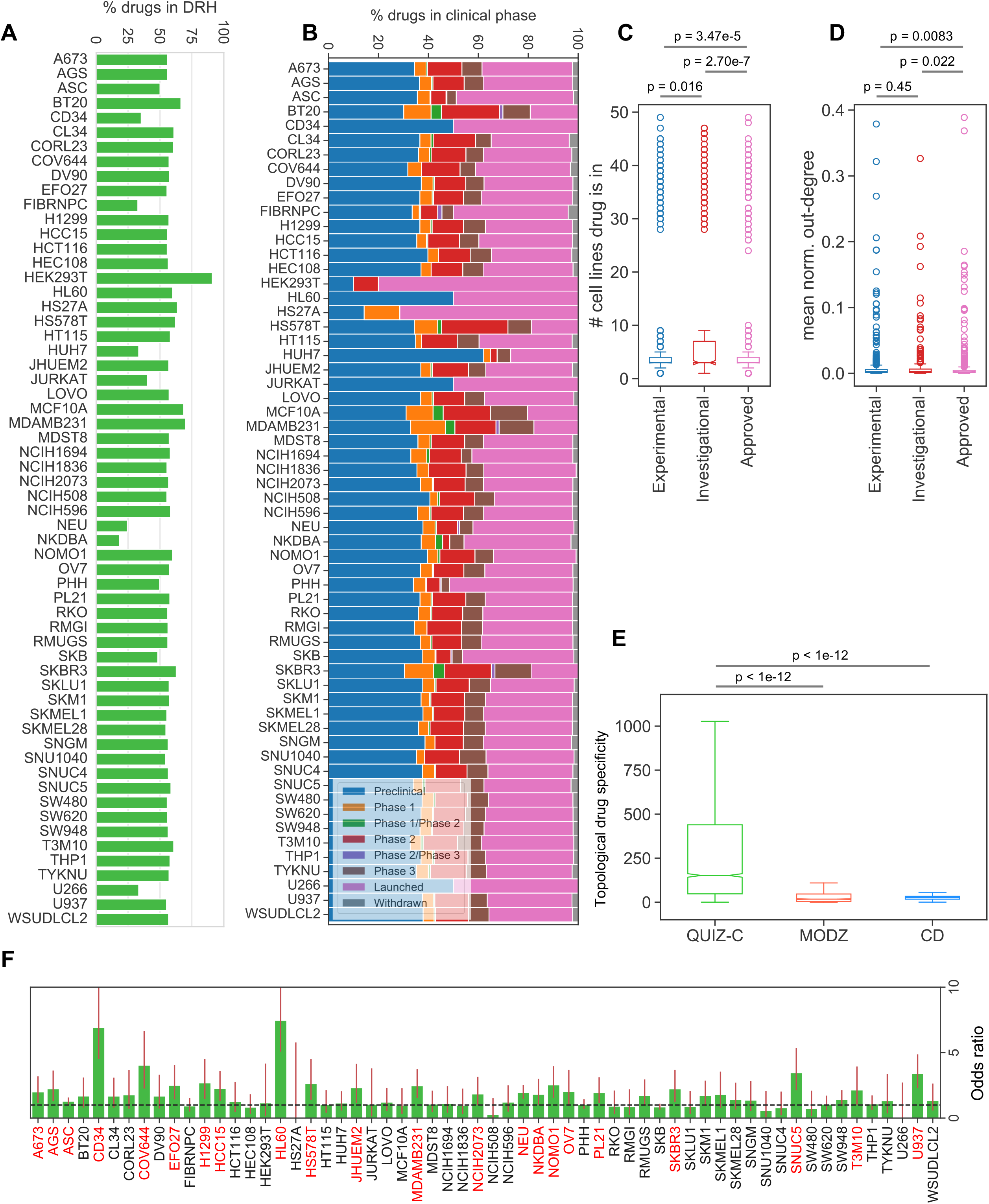
(A) Proportion of drugs in each QUIZ-C network that have MoA information in the Drug Repurposing Hub. **(B)** Clinical development phase breakdown of the drugs in each QUIZ-C network. **(C)** The number of cell type-specific QUIZ-C networks a given drug appears in for experimental, investigational, and approved drugs. **(D)** The mean normalized out-degree of a given drug across QUIZ-C networks, for experimental, investigational, and approved drugs. Reported p-values in (C) and (D) are from twosided Mann-Whitney U tests. **(E)** The topological drug specificity of QUIZ-C. MODZ and CD networks. **(F)** The odds ratio of enrichment of known drug-target interactions in each QUIZ-C network. Error bars indicate 95% confidence intervals and cell lines in red indicate a significant enrichment.

We next sought to answer the question as to whether or not the cell type representation (i.e., the number of cell type-specific networks in which a perturbation is present) and the out-degree (i.e., the number of genes significantly affected by the perturbation) of perturbations can help distinguish among clinical phases of drug development. In all three networks, investigational drugs were present in significantly more (p<0.05, twosided Mann-Whitney U test) cell lines than experimental and approved drugs **(Figure 4C, Supplementary Figures 9C and 9E)**. Comparing out-degrees across development phases revealed that investigational drugs have a significantly (p<0.05, two-sided Mann-Whitney U test) higher out-degree (i.e., higher number of target genes) compared to approved drugs in all three methods, **(Figure 4D**, **Supplementary Figures 9D and 9F)**. Finally, combining the number of cell lines and out-degree, we defined “topological drug specificity” as 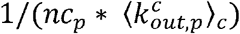 where *nc_p_* is the number of cell lines in which perturbation p is present and 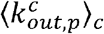 is the mean normalized out-degree of the perturbation over cell lines. Hence, drugs that are exclusive to a few cell types and have few target genes have high topological specificity. QUIZ-C networks had significantly higher topological drug specificity compared to both MODZ and CD networks (two-sided Mann-Whitney U test p-value <10^-12^ for both) **(Figure 4E)**.

### QUIZ-C networks have a higher enrichment of drug targets than other CMap-derived networks

We hypothesized that the degree of enrichment of known drug-target interactions would be an additional indicator of the utility of these networks for predicting novel and repurposed drugs. We used known drug-target interactions from the literature to determine the enrichment of known interactions among the gene-perturbation edges in CMap-derived networks. To maximize coverage of known drug-target interactions, we combined the DRH and the Drug-Gene Interaction Database (DGIdb) (*29*) **(Supplementary Methods)**. The addition of DGIdb data increased overall coverage **(Supplementary Figure 10A)**. Forty-eight percent of QUIZ-C cell type-specific networks had a significant enrichment (two-sided Fisher’s exact p-value < 0.05) of known drug-target interactions **(Figure 4F)** compared to 15% and 13% for MODZ and CD networks, respectively **(Supplementary Figure 10B-C)**. In terms of the difference of log odds-ratios, 74% and 65% of cell lines had a higher enrichment of literature-evidenced drug-target interactions in QUIZ-C compared to MODZ and CD, respectively **(Supplementary Figure 10D-E)**. Together, these results suggest the potential utility of QUIZ-C networks in drug discovery and repurposing.

### Using pathophenotypic profiles to determine the congruity between input signatures and perturbations

The discovery and repositioning of potentially therapeutic drugs using CMap methods have largely focused on the direct comparison of input gene signatures (which can be associated with a disease, treatment, or any other biological context) with perturbation-induced gene signatures (*9*). As such, this approach provides a local view of correlation between the two gene sets that is limited to the biological context of the input gene signature, and is, therefore, agnostic to the larger network of gene-perturbation interactions in other disease contexts. Gene-perturbation networks derived from LINCS-CMap data, on the other hand, enable us to investigate the global effect of perturbations on genes that are implicated in disease etiologies other than those of the input genes. Here, we exploited the fact that QUIZ-C and other CMap-derived networks can be used to study simultaneously the transcriptional response of multiple sets of disease-associated genes to a perturbation in a cell type-dependent manner. In particular, we hypothesized that using a comprehensive network of human disease signatures (pathophenotypes) as a common basis for similarity and dissimilarity between perturbations and input signatures will facilitate and provide additional insights into the prioritization of novel and repositioned drugs.

Here, we describe the PAthophenotypic COngruity Score (PACOS), which uses CMap-derived gene-perturbation networks to predict small molecule and cell line combinations that best enhance or repress a given input signature. PACOS is summarized in **Figure 5** and described in detail in the **Methods** section. In brief, we define the Signature Congruity Score (SCS) to quantify the agreement between two gene signatures in terms of the direction of effect (i.e., an increase or decrease in expression). SCS enables us to assign a scalar value in the range [-1, 1] to any perturbation signature (i.e., gene-perturbation edges of a given perturbation in QUIZ-C or other CMap networks)-disease signature or input signature-disease signature pair **(Figure 5A)**. Extending the SCS calculation to all diseases in a large-scale, multi-sample disease-gene network consisting of 569 disease signatures **(Methods)**, we define Pathophenotypic Congruity Profiles (PCPs), which are vectors composed of SCS values **(Figure 5B)**. We then calculate PCPs for each perturbation in each cell type-specific CMap-derived network, as well as for the input signature. Finally, we calculate PACOS, which quantifies the similarity/dissimilarity between the PCP values of any input signature and cell line-perturbation pair **(Figure 5B)**. A positive PACOS indicates an “enhancing” perturbation whose transcriptional effect is congruent with that of the input signature, whereas a negative PACOS indicates a “repressing” perturbation whose transcriptional effect is incongruent with that of the input signature **(Figure 5B)**. Given an input signature, we rank PACOS values across all perturbations in each cell line, for which we calculate the area under the receiver operating characteristic curve (AUROC). These AUROC values are compared with those of randomized rankings to determine an empirical p-value for each cell line. This procedure enables us to create a nested ranking of cell lines and perturbations whereby the cell lines with the highest statistically significant AUROC values can be queried for the best performing perturbations **(Figure 5C)**.

**Figure 5:**
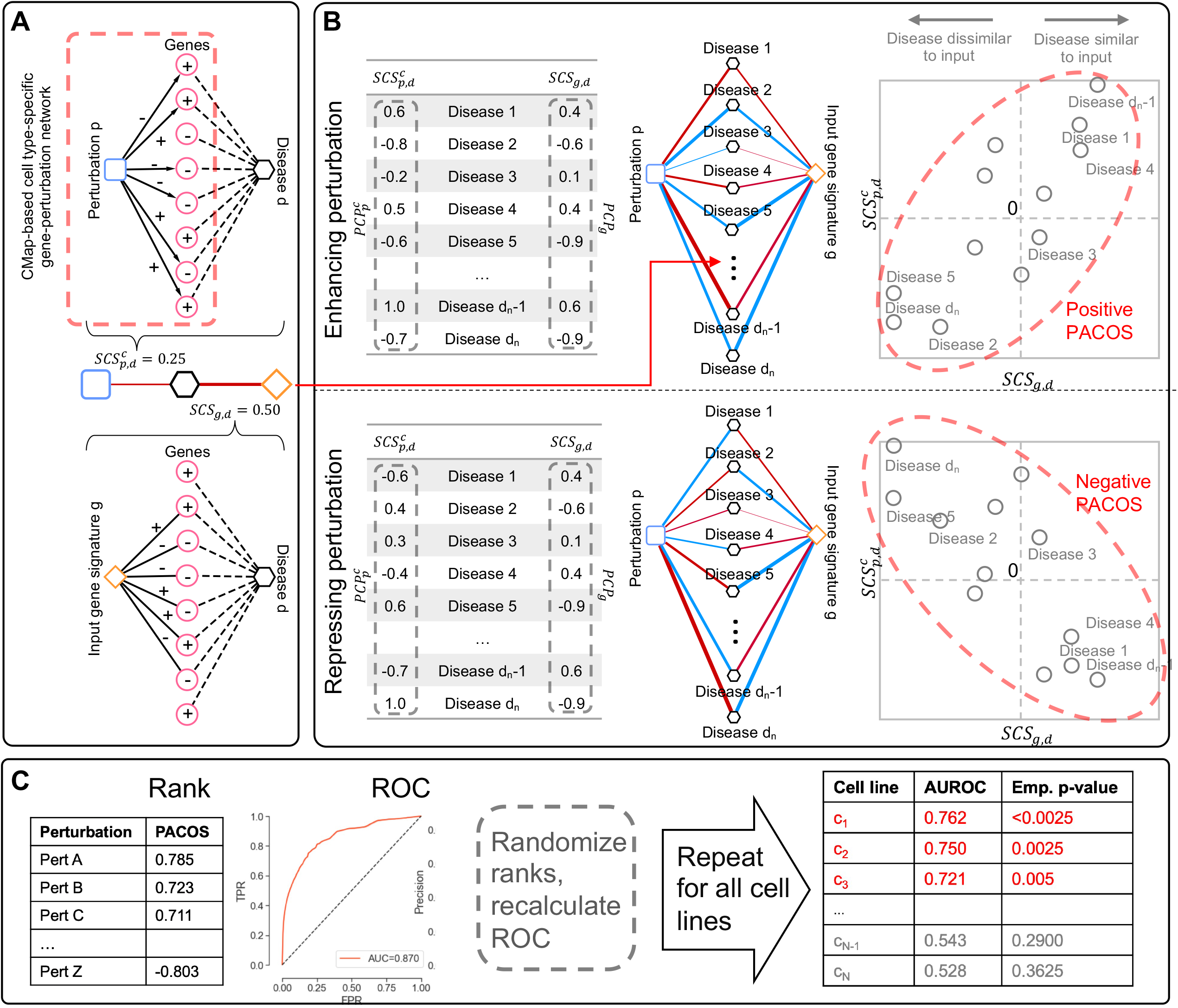
Overview of the PACOS algorithm. Blue squares indicate perturbations, black hexagons indicate intermediary disease phenotypes, orange diamonds indicate input signatures, and pink circles indicate genes. **(A)** The signature congruity score, 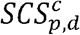, between each disease *d* and each perturbation *p* in cell line *c* is calculated. The same procedure is repeated for the input signature *g* and each disease *d*, resulting in 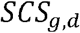. **(B)** Aggregating 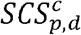 and 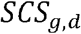 values for all intermediary diseases, we form the 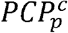 and 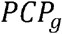 vectors, respectively. The Spearman correlation between 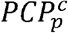 and *PCP_g_* yields the PACOS value for each perturbation for the given input signature in the given cell line. **(C)** Given the PACOS ranking of perturbations in each cell line, receiver operating characteristic (ROC) curves are generated for each cell line. This procedure is repeated for randomized ranks to yield an empirical p-value for each cell line, which can in turn be used to rank cell lines and then perturbations within each cell line.

### Benchmarking Pathopticon

To assess the prediction performance of our framework under various scenarios, we designed a benchmarking strategy that uses 192 gene signatures from the literature as input **(Supplementary Table 2, Supplementary Methods)**. We performed our nested prioritization using the Pathopticon framework: We calculated the AUROC values for all benchmark signatures and cell lines, resulting in 192×60 AUROC values, and identified the statistically significant cell lines. Across all benchmark signatures, we observed a balanced representation of significant cell lines, pointing towards the lack of bias towards certain cell lines in the nested prioritization. Over 80% of cell lines were significant (empirical p-value < 0.05) in at least one benchmark signature in both repressing and enhancing modes. **(Supplementary Figure 11)**. Moreover, repressing and enhancing modes were complementary to each other with non-overlapping sets of significant cell lines.

In our benchmark, we were particularly interested in testing (i) how using QUIZ-C as the underlying gene-perturbation network compares to using other CMap-derived networks; (ii) how PACOS compares with purely cheminformatic, yet input- and cell type-agnostic, measures; (iii) how PACOS compares with other recent CMap-based prioritization methods such as L1000CDS2 (*26*); and (iv) how each of the above comparisons fare in terms of chemical diversity of the top predicted drugs. Below we present the results of these tests.

### QUIZ-C networks surpass other CMap-derived networks in predictive performance within the Pathopticon framework

To compare the predictive capability of PACOS across CMap-derived networks, we focused on the best-performing cell line (i.e., the cell line with the highest AUROC value among significant cell lines) for each benchmark input signature. PACOS-QUIZ-C showed, on average, higher maximum AUROCs compared to both PACOS-MODZ and PACOS-CD in both “enhancing” and “repressing” mode, and the difference was statistically significant (two-sided Wilcoxon signed-rank test p-value < 0.05) in three out of four comparisons **(Figure 6A)**. Due to its being agnostic to the specific input signature, tool score used alone performs poorly compared to PACOS, and similar to random expectation (two-sided Wilcoxon signed-rank test p-value = 0.22) **(Figure 6A)**. Overall, these data suggest that the underlying gene-perturbation network derived from CMap has a non-trivial function in drug prioritization performance and that the cell type-specificity of QUIZ-C networks may play a role in improving prioritization performance in comparison to other CMap-based networks.

**Figure 6:**
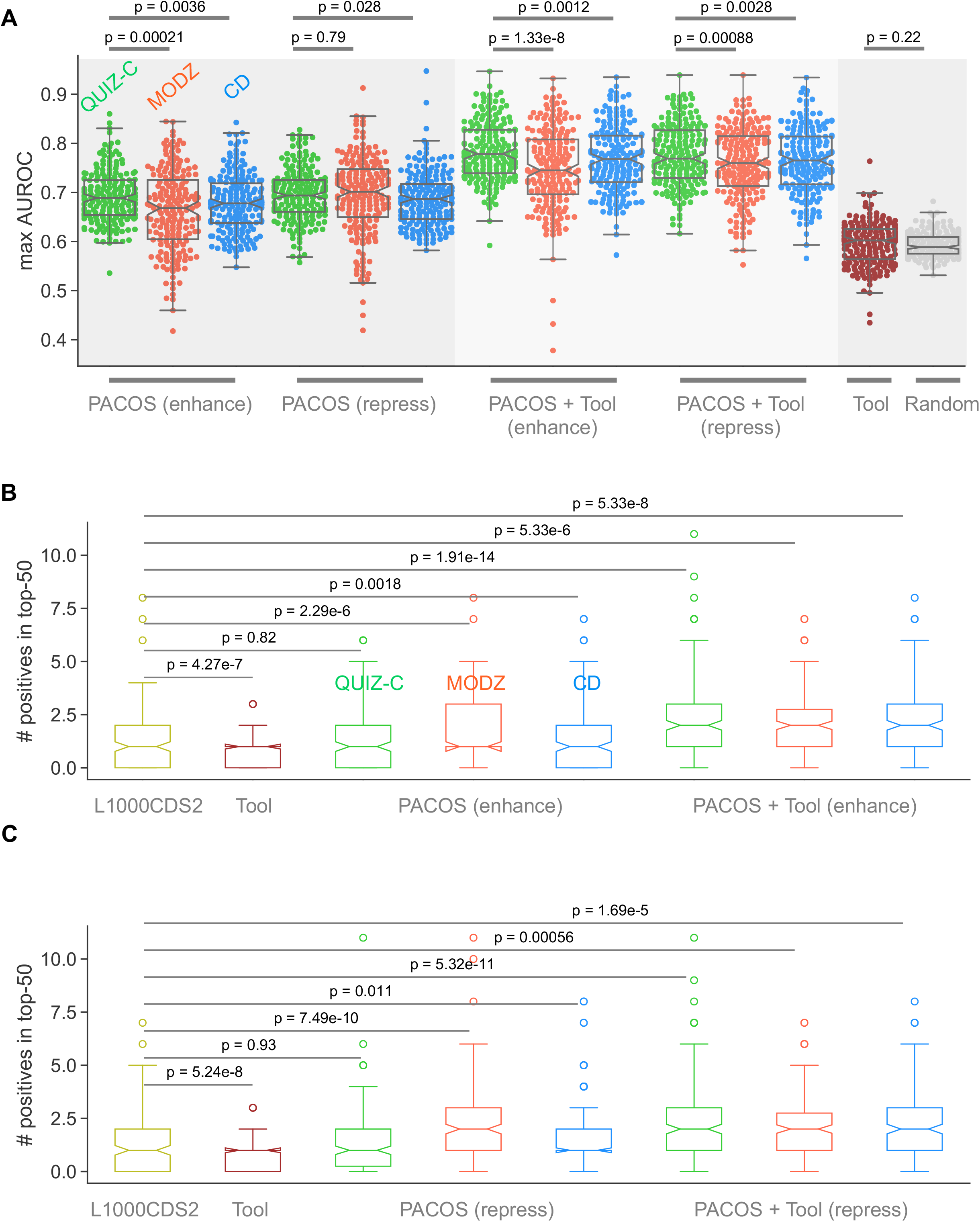
Benchmark results. **(A)** Boxplots of the maximum area under the ROC curves across all benchmark input gene sets for different LINCS-CMap networks and methods. Each dot corresponds to a benchmark input gene set. **(B)** Boxplots of the number of positives (known drug targets) in the top 50 predictions for the forward (L1000CDS2) / enhance (PACOS) mode. **(C)** Boxplots of the number of positives (known drug targets) in the top 50 predictions for the reverse (L1000CDS2) / repress (PACOS) mode.

### Integrating pharmacogenomic and cheminformatic data improves cell typespecific drug prediction performance

Despite being frequently used as reliable predictors of potential novel or repurposed small molecules (*30*), ligand-based measures such as binding selectivity and tool score (*31*) derived from bioactivity data (*32*) largely focus on single protein targets. As such, these measures prioritize compounds by taking as input single targets rather than multiple targets (*24*) – an approach that is at odds with recent bioinformatic findings that drugs have, on average, 32 targets in the proteome (*33*). In this sense, these cheminformatic measures are agnostic to disease context in the form of multiple disease-associated genes. They nevertheless provide useful information on chemical binding properties of a given drug, a feature that is crucial in drug development. Given the drawbacks of solely relying on pharmacogenomic data in drug discovery (*34, 35*), we hypothesized that cheminformatic and pharmacogenomic information are complementary, and that their integration will be more beneficial in drug discovery than either approach alone. To test this hypothesis, we used a heuristic measure that optimally combines PACOS with tool score **(Methods, Supplementary Figure 12)**. Supporting our hypothesis, the integration of PACOS and tool score significantly surpassed PACOS alone in all three cases in both enhancing and repressing mode (two-sided Wilcoxon signed-rank test p-value < 10^-12^ in all cases), with PACOS-QUIZ-C-Tool outperforming both PACOS-MODZ-Tool and PACOS-CD-Tool in both modes **(Figure 6A)**. The integration with cheminformatic data also increased the number of significant cell lines identified by Pathopticon **(Supplementary Figure 13)** compared to LINCS-CMap data alone **(Supplementary Figure 11)**.

As an additional external benchmark, we compared PACOS to a state-of-the-art method that also uses LINCS-CMap data to prioritize drugs, namely L1000CDS^2^ (*26*). Comparison with L1000CDS^2^ is important since L1000CDS^2^ is another tool that makes full use of LINCS-CMap data and simultaneously ranks drug candidates along with the relevant cell lines. In terms of the number of known targets captured in the top 50 candidates, L1000CDS^2^ performed significantly (two-sided Wilcoxon signed-rank test p-value < 0.05) better than tool score. On average, PACOS had a higher number of known targets in the top 50 candidates compared to L1000CDS^2^. While this difference was significant (two-sided Wilcoxon signed-rank test p-value < 0.05) for PACOS-MODZ and PACOS-CD, it was comparable to L1000CDS^2^ for PACOS-QUIZ-C. When combined with tool scores, all three networks significantly surpassed L1000CDS^2^. These sets of results were replicated for both the enhancing **(Figure 6B)** and repressing **(Figure 6C)** modes.

Together, these results demonstrate the predictive advantage of integrating cheminformatic data with LINCS-CMap data for cell type-specific drug prioritization, and suggest that QUIZ-C networks best complement cheminformatic data compared to the other CMap-based networks. Moreover, the disease pathophenotypes embedded in PACOS can provide important insights about which disease processes the input gene signature and the perturbation signatures simulate. In the following section, we highlight this point with a biological example.

### High chemical diversity of the PACOS-prioritized drugs

While the top-prioritized compounds can be selected as potential candidates for downstream validation studies, a larger set of highly ranked drug candidates can also be thought of as a screening set of compounds. In this case, it is desirable that the set of drugs chosen for screening are structurally diverse rather than similar to ensure the exploration of a larger portion of chemical space (*24, 30, 36*). We, therefore, sought to quantify the structural diversity of the top-ranked drugs for each benchmark signature. Overall, the chemical structure similarity, measured by the Tanimoto coefficient **(Supplementary Methods)**, was low for all methods with a median value between 0.2 and 0.4 **(Supplementary Figures 14-17)**, in agreement with a recent report that suggested that setting a Tanimoto structural similarity threshold of 0.2 was an effective proxy for chemical diversity (*24*). For the majority of benchmark signatures, QUIZ-C had significantly lower (two-sided Mann-Whitney U p-value <= 0.05) Tanimoto similarity (therefore higher structural diversity) in the top-50 predicted drugs than MODZ, CD, or L1000CDS2 for both enhance and repress modes and with or without tool scores **(Supplementary Table 3, Supplementary Figures 14-17)**. These results suggest that the top drugs predicted by Pathopticon hold the potential for being used as cell type-aware compound screening libraries focused on a specific input signature.

### Pathopticon identifies potential therapeutic candidates to counter macrophage-driven vascular inflammation

To demonstrate the utility of Pathopticon for discovering potential therapeutic agents, we used omic data from our previous preclinical study of vein graft disease in mice (*37*). Vein graft failure remains an unmet clinical burden, with its high initial-year event rates and no effective medical treatment (*38, 39*), among patients with coronary or peripheral artery disease who subsequently underwent myocardium sparing or limb-saving vein bypass surgery. For our use case, we set out to predict candidate drugs that may alter the state of inflammation, which is an important contributor to this disease. To this end, we used proteomic data derived from diseased vein graft tissue (plaque enriched- and disease time-course-) and sought to repress two gene signatures corresponding to early and late stages of the disease **(Methods).** The majority of the predicted drugs (13 out of the top-20 predictions for both stages combined) had literature evidence of clinical investigation in the context of vascular medicine **(Supplementary Table 4; see Supplementary Information for an overview of the available literature).** For downstream *in vitro* validation, we focused on a subset of these 13 candidates that were predicted to reverse significantly the late-stage vein graft disease signature in adipose stromal cells (ASCs), a mixed population of cells containing adipose progenitor cells (APCs) and fibro-inflammatory precursors, including macrophages, important in a variety of vascular diseases **(Figure 7A, see Supplementary Information for additional references)**. The diseases that had a high signature overlap with vein graft disease for each of these top candidates were enriched in inflammation related pathways **(Figures 7A, 7B, Supplementary Files 1-2)**. To demonstrate the direct and quantifiable modulation of inflammation by these candidates, we conducted gene expression studies by real-time polymerase chain reaction (qPCR) **(Methods).** The majority of the predicted drugs decreased the expression of at least one of the important inflammatory genes (*37, 40*) in lipopolysaccharide (LPS)-stimulated human primary macrophages, M(LPS), such as CCL2, CCL5, CCL7, CCL8, GBP1, RELB and S100A10 **(Figure 7C, Supplementary Figure 18)**. Known macrophage antiinflammatory genes such as CCL22, APOE, CHI3L1, PPARA, and ARG1, however, remained unaltered for the majority of the tested drugs, with the exception of the PPAR-α agonist GW-7647, the hormone estriol, and the p38 MAPK inhibitor SB-203580 **(Figure 7C, Supplementary Figure 19)**, all of which have been shown to be antiinflammatory and athero-protective (*37, 41–43*). Similar gene expression signature transitions from pro- to anti-inflammatory states may also benefit, by way of therapy, other atherothrombotic vascular diseases such as restenosis following stenting for coronary artery disease (*44*), arterio-venous fistula disease, and peripheral artery disease.

**Figure 7:**
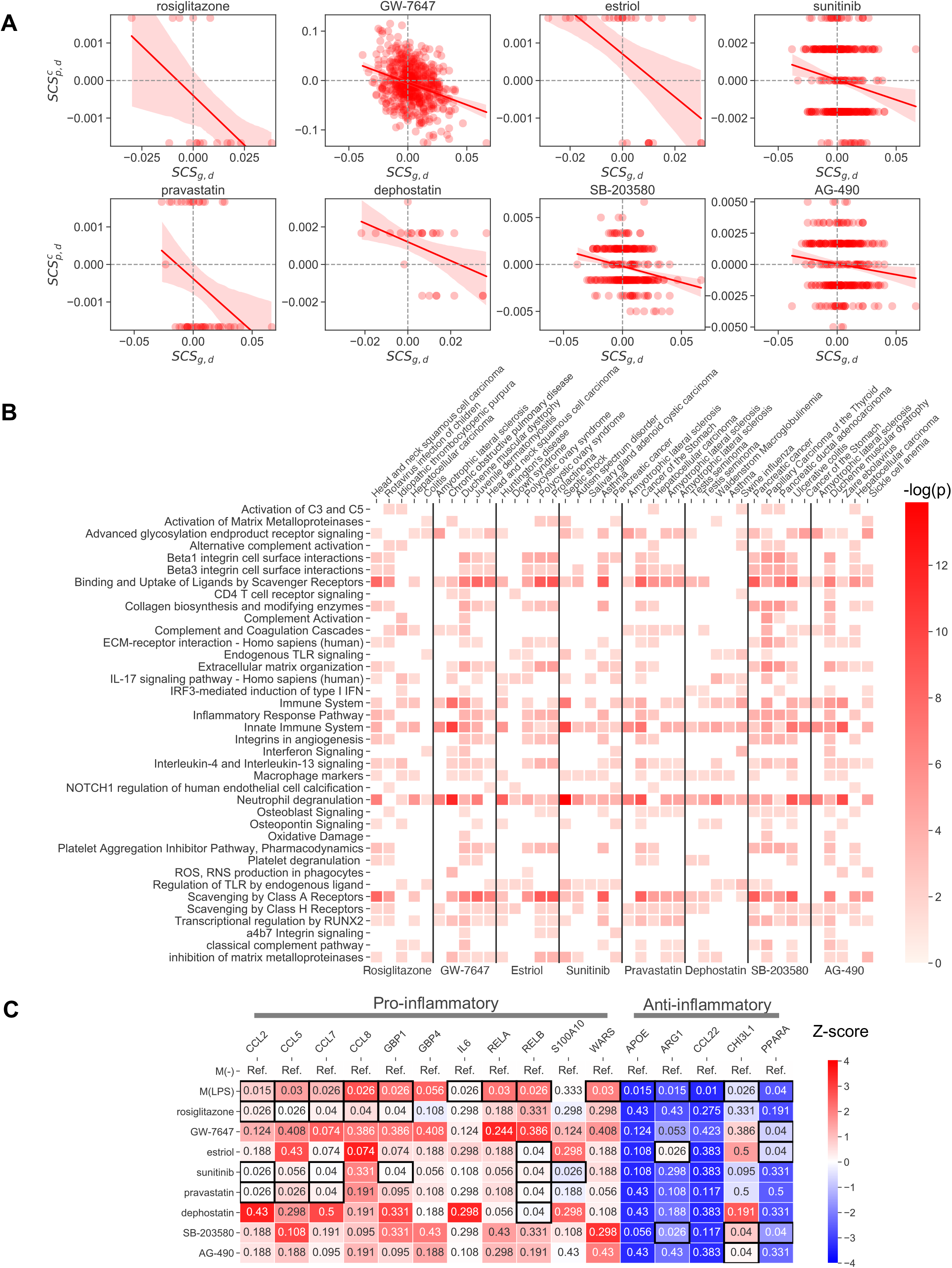
*In vitro* validation. (A) The eight drug candidates repressing the vein graft disease signature that were selected for *in vitro* validation. The signature congruity score, 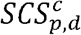, between each disease *d* and each perturbation *p* in cell line *c*, is shown on the y-axis and the signature congruity score for the input signature *g* and each disease *d, SCS_g,d_*, is shown in the x-axis. Each circle represents an intermediary disease phenotype. The lines indicate the estimated regression model and the shaded regions indicate the 95% confidence intervals. (B) The pathway enrichment of the top five disease phenotypes with the highest signature overlap with the vein graft disease signature (i.e., the leftmost and rightmost diseases on the x-axis). Darker colors indicate a higher enrichment in terms of the -log(p) value and empty cells indicate instances where the enrichment was not significant (p>0.05). (C) Summary of the results of the q(PCR) experiments. Z-scores are calculated separately for each column (i.e., each gene) and centered around the reference state (i.e., the undifferentiated macrophage M(-)). Black framed cells indicate statistically significant changes (p<0.05, two-sided Mann-Whitney U test) with respect to M(LPS) for the candidate drugs, and with respect to the baseline (i.e., M(-)) for M(LPS).

## Discussion

In this study, we approach cell type-guided computational drug discovery as a two-stage prioritization task that involves the simultaneous ranking of cell lines and perturbations. Based on our broad overview of the LINCS-CMap data, we propose a statistical consensus approach to generate cell type-specific drug-perturbation networks. We designed a drug discovery framework, Pathopticon, that integrates these drug-perturbation networks with diverse disease phenotypes and compound bioactivity data. Through extensive benchmarks, we demonstrate that Pathopticon surpasses currently available methods in terms of its prediction performance and the chemical diversity of its predicted candidates. We provide our framework as a user-friendly web app through which users can explore each cell type-specific network, make predictions based on their omics experiments, and analyze the results.

The scope of our gene-perturbation networks is complementary to that of the state-of-the-art and makes maximal use of the LINCS-CMap data: unlike the majority of CMap-based approaches that focus on the nine cell lines in the LINCS-CMap reference (“Touchstone”) dataset, we generate networks for 60 cell lines outside the Touchstone set. While cell lines have proven to be versatile instruments that allow for the study of multiple disease contexts (*45*), they are also often highly specific in terms of their transcriptional and phenotypic response to perturbations by drugs (*46*). In line with this observation, our extensive benchmarks showed the advantage of QUIZ-C over other CMap-based gene perturbation networks in terms of reflecting the unique transcriptional response of each cell line to perturbations.

Our signature congruity score (SCS), which is the base scoring component underlying PACOS, is not novel by itself. It is akin to methods previously introduced such as XSum, XCos (*47*), or more recently, EMUDRA (*48*), in that it is a simple metric quantifying the agreement between an input gene set and a perturbation signature (*49, 50*). One characteristic of SCS that can arguably make it advantageous to these similar scores is that it does not require the fold-change values of the input signature, which may not always be available to the researcher. What makes PACOS unique as a prioritization method, rather, is that candidate drug predictions are made based on the agreement of the transcriptional signatures of the drug and input signature on a large ensemble of diseases, as opposed to predictions directly based on the similarity of input and drug signatures. This feature becomes highly informative as it introduces an additional layer of information about which disease signatures are similarly affected by the perturbation and the input signature, thereby giving mechanistic clues about the shared intermediary pathophenotypes (endophenotypes) (*51*) targeted by the predicted candidates. For example, in our use case, we showed that all diseases implicated in the top predictions have a common inflammatory component **(Figure 7B)**, suggesting that the inflammation endophenotype lies at the core of these predictions.

Despite having an ample amount of compound bioactivity information on single targets in humans, only 12% of this portion of ChEMBL contained information about cell lines at the time these data were retrieved. We hypothesized that a way to maximize the utility of ChEMBL in the context of cell type-specific drug discovery is to integrate it with resources that have complete cellular context, such as LINCS-CMap. In favor of this hypothesis, our benchmarks showed that these resources combined results in a prediction performance that surpasses that of each resource alone.

Our study has several limitations due to its reliance on large public databases, the first one of which concerns LINCS-CMap data itself. Although Pathopticon is a generalizable computational model whose primary goal, like other CMap-based approaches, is to expedite drug discovery and repurposing, we acknowledge the recent concerns about the reproducibility within and between different CMap versions (*52*). We, therefore, recommend general caution in interpreting the results and encourage biological followup studies for the identified drug candidates. In particular, if candidates have been identified using consensus methods such as ours, a careful scrutiny of the time points and doses of the prioritized drugs in the CMap data should accompany subsequent *in vitro* studies. A second limitation pertains to the ChEMBL data, which, being an enormous database in which the level of annotation of compounds varies widely, is not without data quality issues and possible erroneous entries, such as negative nanomolar values (*53*). Diligent curation and data cleaning efforts are, therefore, warranted in studies utilizing this resource.

While LINCS-CMap’s cell line coverage is remarkable for a high-throughput genomic resource, it is still a limited subset of cell types available to researchers, which may lead to differences with the cell types on which omics data were generated. Should this discordance prevail, one can consider leveraging heterologous cell types that may colocalize in the same organ with the cell type of primary interest. For example, distinct cell types have been shown to act in concert in vascular diseases via heterologous cellcell interactions mediated by metabolic exchange (*54*). Therefore, compounds that target “biochemically adjacent” cell types can be used in lieu of the available cell type. Alternatively, single-cell RNA-seq can help shed light on similar cell subpopulations, leading to the identification of “transcriptionally homologous” cell types to the LINCS-CMap cell lines. Finally, since the cell types available in LINCS-CMap are a mixture of immortalized cell lines, primary cells, and end-differentiated cells, it is important to be cognizant of the challenges in interpreting the perturbation signatures in these distinct and often vastly different genomic contexts.

The mechanisms of action of the majority of the top-ranked drugs predicted by Pathopticon in our use case were supported by previous investigations in the context of vascular disease. By contrast, three predictions appeared to be relevant but effective in the opposite direction, and four predictions were approved for other indications not directly relevant to vascular disease (**see Supplementary Information for an extended discussion and more references)**. While we gave priority to a subset of the drugs with literature evidence to validate *in vitro*, this latter set of drugs may have repurposing potential and warrants a closer look at their mechanisms of action and tissue-specific targets.

Extensions to our work might include increased integration of bioactivity and cell typespecific multi-omic data, especially as the latter become more widely available with multi-institutional efforts such as the Trans-Omics for Precision Medicine (TOPMed) program. Different omic modalities can then be incorporated in our pathophenotypic congruity scoring schema, which would allow for the query of therapeutic agents that invoke similar signatures in terms of multiple omic types. The current boon in single-cell resolution sequencing methods and resources will undoubtedly help refine further celltype guided drug discovery in the near future.

## Materials and Methods

### <u>Qu</u>antile-based <u>I</u>nstance <u>Z</u>-score <u>C</u>onsensus (QUIZ-C)

We collected the Level 4 plate-normalized expression values (referred to as ZS (https://clue.io/connectopedia/data_levels) or ZSPC values (*18)*) of all perturbation instances for each gene. We then calculated a gene-centric z-score for each of these perturbation instances, denoted as 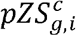, by comparing the Level 4 expression value of gene *g* when perturbed by instance *i* in cell line *c*, 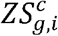, against the ZS values of all perturbation instances for the same gene and cell type,

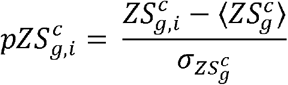

where 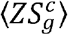 and 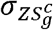 are the mean and standard deviation of ZS scores over all perturbation instances for the given gene *g* and cell type *c*, respectively. Thus, 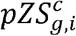 ensures that we capture the differential expression of perturbation instance *i* against the background of the entire population of perturbation instances in the given cell type *c*, in addition to the 384-well plate population. After calculating the 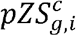 values for all perturbation instances, we aggregated these values at the perturbation (i.e., “pert_iname”) level to obtain the set of gene-centric z-scores for each perturbation such that

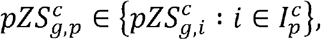

where 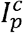 is the set of all instances for perturbation *p* in cell type *c*.

Once the gene-specific z-scores were calculated for all possible cell type, gene, and perturbation (*c, g, i*) triplets and aggregated into perturbation-level groups, we sought to identify perturbation-gene pairs in which the perturbation affects the expression of the target gene (i) significantly and (ii) consistently. The first condition for significance of effect is satisfied if 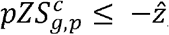, for 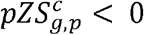 (decrease in expression) or 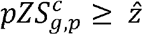, for 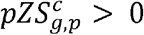 (increase in expression) where 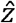 is the z-score at the chosen significance threshold. The second condition for consistency of effect, or consensus, is satisfied if a certain portion of 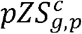 values meet the significance criterion based on their distribution. However, due to the variation in the number of perturbation instances across different cell lines **(Supplementary Figure 1)**, as well as the heterogeneity of expression values within and across perturbations due to varying doses and time points, it is not feasible to use a fixed number of perturbation instances as the consensus criterion. We, therefore, introduced a flexible consensus threshold that can accommodate the individual 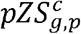 distribution of any given cell type, gene and perturbation triplet **(Figure 2)**, which we call the Quantile-based Instance Z-score Consensus (QUIZ-C) approach. Depending on whether perturbation p induces a decrease or increase in the expression of gene *g* in cell type *c*, we designate the 1^st^ or the 3^rd^ quartile (25^th^ or 75^th^ %ile) of the 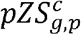 distribution, respectively, as the consensus threshold. If the number of significant instances (i.e., above a certain z-score threshold 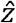, taken to be 1.96 here) for the given perturbation is greater than the number of instances below or above this threshold, then the consensus is satisfied **(Figure 2C)**, and the perturbation is deemed to affect the expression of the gene significantly and consistently in the given cell type. A positive or negative edge 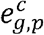 with a weight 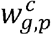 equal to +1 or −1, respectively, is then established between the perturbation-gene in the cell type of interest pair such that

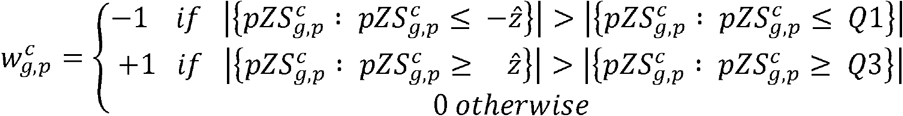

The positive and negative edges 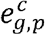 satisfying the above conditions were then collected into edgelists to be used in the building of the cell type-specific drug-gene networks **(Figure 2D)**. Due to the large scale of the LINCS-CMap Level 4 data (>1.3M perturbation instances, >12k edges and 60 cell lines), we used the *multiprocess* package in Python (https://pypi.org/project/multiprocess/) to perform the identification of edges in parallel for each cell line (coda available at https://github.com/r-duh/Pathopticon/blob/main/generate_QUIZC_networks.py). Our sensitivity analysis showed that the resulting QUIZ-C networks were largely robust to the choice of quantiles used under various regimes and model choices **(Supplementary Methods),** and different quantiles yielded similar network topologies, as quantified by the edge overlap **(Supplementary Figure 20).**

### Calculating the PAthophenotypic COngruity Score (PACOS)

We obtained a comprehensive list of disease signatures curated from the Gene Expression Omnibus (GEO) through the Enrichr web server (*55*), namely the “Disease_Perturbations_from_GEO_up” and “Disease_Perturbations_from_GEO_down” gene set libraries (https://maayanlab.cloud/Enrichr/#stats, accessed September 2020). These gene sets contain genes induced (“up”) and repressed (“down”) in 569 human disease samples, representing a broad array of diseases. Overall, these 569 gene sets comprise 21,787 genes, covering the majority of the protein-encoding genome.

To measure the degree of local agreement in terms of the direction of effect between a perturbation signature and a disease signature in a given cell line, we define the signature congruity score (SCS) as

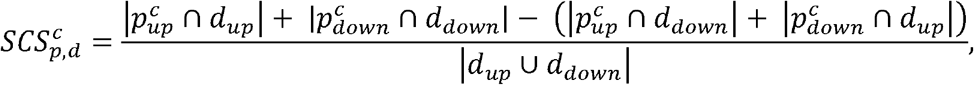

where 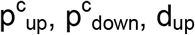, and d_down_ are the sets of up and down genes connected to perturbation *p* in the QUIZ-C/MODZ/CD network for cell type *c*, and the sets of up- and down-regulated genes for disease *d*, respectively.

We similarly define the *SCS* between an input gene signature and a disease signature as

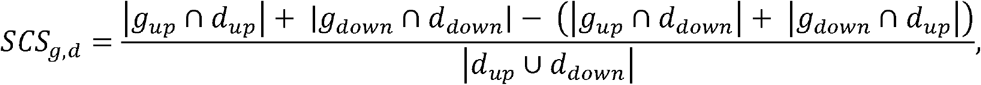

where g_up_, g_down_, d_up_, and d_down_ are the sets of up- and down-regulated genes for input gene set *g* and disease *d*, respectively.

To differentiate between the case where *SCS* is zero due to the congruent (same direction) and incongruent (opposite direction) effects canceling each other out, and the case where *SCS* is zero simply due to zero overlap between the input/perturbation signature and the disease signature, we calculate 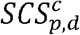 and 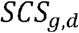 only if the perturbation signature and disease signature (or input gene signature and disease signature) have at least one common gene. In the case of no overlap between the input/perturbation signature and the disease signature, we assign a null value (NaN in Python) to *SCS.*

*SCS* values are, thus, bound to be in the range [-1, 1] where −1 and 1 represent complete incongruity and congruity of the given perturbation signature/input gene set with the given disease signature, respectively. Complete congruity and incongruity are hard to achieve since disease signatures have on average a few hundred up- and down-regulated genes whereas the number of genes affected by a perturbation or the input gene sets can be much smaller **(Figure 2C)**.

Expanding the signature congruity concept to all 569 disease signatures in our dataset, we generate pathophenotypic congruity profiles (PCP) for the given perturbation p or input gene set *g*, which are vectors of length *n_d_* = 569 (including null values)

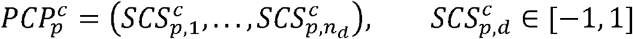

and

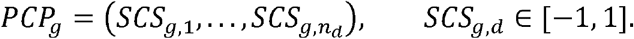

For each cell type-specific CMap-based gene-perturbation network, we precompute 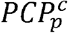 values for all perturbations within that cell type. This computation is performed only once per network and stored as a binary file (see https://github.com/r-duh/Pathopticon/ for file location) to speed up downstream candidate prioritization and benchmarking steps and reduce computational burden.

Finally, we define the pathophenotypic congruity score (PACOS) between input gene set *g* and perturbation *p* in cell line *c* as the Spearman correlation between 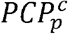 and 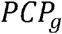

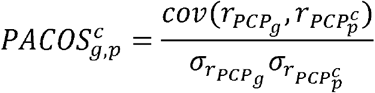

where *r_PCP_g__* and 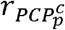 denote the corresponding rank vectors of *PCP_g_* and 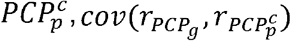 denotes the covariance of the rank vectors, and *σ_r_PCP_g___* and 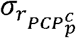 denote the standard deviations of the rank vectors. Hence, 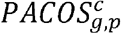 is a measure of how well a perturbation’s global effect across a wide range of pathophenotypic signatures enhances or represses that of an input gene signature. To ensure that the Spearman rank correlation yields reliable results, we calculate 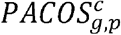 only if the number of diseases that are common between the input gene signature and the perturbation-cell type pair is above a threshold, which is set to 10 by default.

### Pathopticon: An analytical framework for drug prioritization that integrates CMap-based gene-perturbation networks with PACOS and tool scores

In the Pathopticon framework **(Figure 1)**, the *PCP_g_* vector of a given input gene signature *g* is compared across all cell types and perturbations to 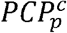, resulting in 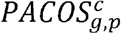 values for all perturbation-cell type pairs. These 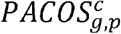 values are then integrated with tool scores of perturbation *p* to generate a ranked list of perturbation-cell type pairs, the top ranks of which offer potential viable therapeutic candidates with high selectivity that target the input gene signature. We scale the tool scores between −1 and 1 to be compatible with PACOS scores and then impute the missing values using the median value across all perturbations. We then collapse all drug-target pairs for a given drug to a single value by selecting the highest selectivity/tool score for that drug **(see Supplementary Methods and Supplementary File 3)**.

Since we hypothesize that incorporating binding selectivity information will increase candidate drug prediction performance for a given input gene signature *g*, we determine our final prioritization rankings based on a weighted average of PACOS and tool scores

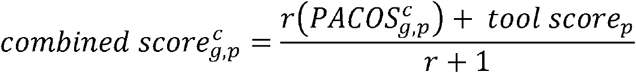

However, using tool scores by itself inherently limits the number of drugs that can be highly prioritized since not all drugs in the CMap-based networks have bioactivity information available on ChEMBL **(see Supplementary Methods and Supplementary File 4)**. Therefore, the drugs that do not have this information are pushed down the ranks if tool scores are weighed heavily, effectively constraining the number of drugs that can be ranked highly. To counter this bias, we give more weight to PACOS when combining PACOS and tool scores with the primary objective of expanding the pool of therapeutic candidates that can be top-ranked. In particular, to address the tradeoff between expanding the pool of drugs to prioritize and getting a high prediction performance, we take a heuristic approach in determining how to weigh each component in the final combined score. For different values of the ratio of PACOS to tool score, *r*, we calculate the median AUROC over all benchmark signatures and simultaneously record the number of unique drugs highly ranked in the top 50 ranks across all benchmarks to represent the diversity of the perturbation space explored in the prioritization. Varying *r* over a range of values from 0 to 6, we choose a value that retains a high value of unique drugs in the top ranks while not compromising a high AUROC. Based on this heuristic test, we set a combined score ratio of r = 2 where the number of unique drugs prioritized plateaus while the corresponding AUROC (0.69) is not substantially different than the maximum AUROC achieved (0.70) **(Supplementary Figure 13)**.

### Benchmarking and cell type-specific performance assessment of Pathopticon

As the input signatures for our benchmark, we used 192 gene sets from the Molecular Signatures Database (MSigDB) (*56*) that have 20 or more known drugs targeting them in the Therapeutic Target Database (TTD) database (*57*) **(Supplementary Table 2, Supplementary Methods)**. Since the number of perturbations in each cell type is highly heterogeneous **(Supplementary Figure 1B)**, we devised, and applied on each benchmark signature, the following cell type-specific performance assessment: We first ranked the 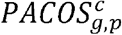 (or 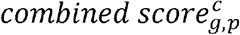, depending on which model was chosen) values separately within each cell line *c*. We then mapped the true positives corresponding to the input gene signature on this list of perturbations ranked by 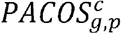 to determine the true positive rate (TPR) and false positive rate (FPR) values. We used the TPR and FPR values to plot the receiver operating characteristic (ROC) and finally measured the area under the ROC curves (AUROC). This procedure resulted in AUROC values for each cell type given the input gene signature. Next, to assess whether these AUROC values are statistically significantly higher than random expectation, we generated AUROC curves that correspond to permuted rankings for N=400 random permutations. Finally, we calculated empirical p-values such that

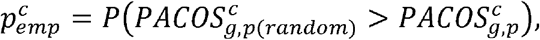

where 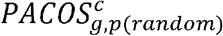 is the score of the randomized instance. This procedure allows a nested prioritization in which each cell line is ranked based on its corresponding 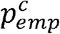 value and each perturbation within that cell line is ranked based on its 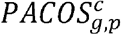 value.

Since our method is intended as a tool to recommend the best cell type and compound combinations given an input gene signature, when comparing AUROCs for our benchmark, we used the AUROCs of the best-performing cell lines, i.e., the maximum AUROC value across all cell lines. Following this approach, we calculated the AUROC for each input signature for (i) PACOS ranking in both repress and enhance mode, (ii) combined (PACOS + Tool) ranking in both repress and enhance mode, and (iii) tool score ranking only. As an additional, network-agnostic negative control, we calculated AUROCs by randomizing the drug rankings 400 times, for which we used the best AUROC value among the random instances in the benchmark. Overall, we had 14 sets of AUROCs across different prioritization and network building methods for performance comparison in our benchmark **(Figure 6A)**. We note that tool scores also take into account the best tool score for a drug-target pair **(Supplementary Methods)**, implying that the corresponding AUROC value for the tool score rankings reflect the best possible AUROC. Thus, further consistency and fair comparison in the benchmark was attained by ensuring the comparison of the best results from each method.

### Running Pathopticon on vein graft gene signatures

We used time-course proteomic measurements, described in detail in (*37*), to identify two gene signatures for vein graft disease, corresponding to early- and late-stage disease. As a proxy of early (late) disease, we used proteins measured on Day 1 and Day 3 (Day 7 and Day 28). We took the union of all proteins with false discovery rate (FDR)<0.05 and fold-change |FC|>2, at Days 1 and 3 and Days 7 and 28, respectively, for early and late signatures. We then used these as “up” (FC > 2) and “down” (FC < 0.5) genes and ran Pathopticon with the following parameters: Model (PACOS+Tool combined); Effect: Repress; r value: 2.0; Number of randomizations: 400.

### In vitro validation methods

An *in vitro* model of proinflammatory macrophages relevant to cardiovascular inflammation was used to test drug predictions. Primary human macrophages were derived from peripheral blood mononuclear cells sourced from buffy coats of healthy adults as previously described (*37*). After ten days of culture, sample sets of macrophages were exposed to 10 ng/mL of lipopolysaccharide, LPS (PeproTech), and co-incubated with either 0.1% (in culture media) of 1x phosphate buffered solution, PBS (ThermoFisher), containing 4 ppm of dimethylsulfoxide (DMSO) or one of the identified drugs at 10 μM concentration for 16 hours. RNA extraction (GE illustra RNAspin Mini, Cat# 25-0500-70) followed by gene expression studies using real-time qPCR (Standard BioTools Inc., Fluidigm BioMark HD) with TaqMan Gene Expression Assay probes, FAM (ThermoFisher) were employed. A sample set of unstimulated macrophages (PBS only, no LPS) was included to represent the baseline non-activated state. GAPDH and RPLP0 were used as endogenous controls to normalize gene expression data. Twotailed Mann-Whitney U tests were employed to assess the differences between treatments and controls. Pathway enrichment analysis was done by performing twotailed Fisher’s Exact tests on pathway data from the ConsensusPath Database (*58*) and adjusting the p-values for multiple testing using the Benjamini-Hochberg procedure.

## References

1. A. L. Hopkins, Network pharmacology, Nat. Biotechnol. 25, 1110–1111 (2007).

2. S. I. Berger, R. Iyengar, Network analyses in systems pharmacology, Bioinformatics 25, 2466–2472 (2009).

3. S. Zhao, R. Iyengar, Systems pharmacology: Network analysis to identify multiscale mechanisms of drug action, Annu. Rev. Pharmacol. Toxicol. 52, 505–521 (2012).

4. G. Hu, P. Agarwal, I. K. Jordan, Ed. Human Disease-Drug Network Based on Genomic Expression Profiles, PLoS One 4, e6536 (2009).

5. E. Guney, J. Menche, M. Vidal, A.-L. Barábasi, Network-based in silico drug efficacy screening, Nat. Commun. 7, 10331 (2016).

6. F. Cheng, C. Liu, J. Jiang, W. Lu, W. Li, G. Liu, W. Zhou, J. Huang, Y. Tang, R. B. Altman, Ed. Prediction of Drug-Target Interactions and Drug Repositioning via NetworkBased Inference, PLoS Comput. Biol. 8, e1002503 (2012).

7. F. Cheng, I. A. Kovács, A.-L. Barabási, Network-based prediction of drug combinations, Nat. Commun. 10, 1197 (2019).

8. J. Lamb, E. D. Crawford, D. Peck, J. W. Modell, I. C. Blat, M. J. Wrobel, J. Lerner, J.-P. Brunet, A. Subramanian, K. N. Ross, M. Reich, H. Hieronymus, G. Wei, S. A. Armstrong, S. J. Haggarty, P. A. Clemons, R. Wei, S. A. Carr, E. S. Lander, T. R. Golub, The Connectivity Map: Using Gene-Expression Signatures to Connect Small Molecules, Genes, and Disease, Science (80-.). 313, 1929–1935 (2006).

9. A. Musa, L. S. Ghoraie, S. D. Zhang, G. Glazko, O. Yli-Harja, M. Dehmer, B. Haibe-Kains, F. Emmert-Streib, A review of connectivity map and computational approaches in pharmacogenomics, Brief. Bioinform. 19, 506–523 (2018).

10. L. Huang, S. Garrett Injac, K. Cui, F. Braun, Q. Lin, Y. Du, H. Zhang, M. Kogiso, H. Lindsay, S. Zhao, P. Baxter, A. Adekunle, T.-K. Man, H. Zhao, X.-N. Li, C. C. Lau, S. T. C. Wong, Systems biology–based drug repositioning identifies digoxin as a potential therapy for groups 3 and 4 medulloblastoma, Sci. Transl. Med. 10, eaat0150 (2018).

11. J. T. Dudley, M. Sirota, M. Shenoy, R. K. Pai, S. Roedder, A. P. Chiang, A. A. Morgan, M. M. Sarwal, P. J. Pasricha, A. J. Butte, Computational repositioning of the anticonvulsant topiramate for inflammatory bowel disease, Sci. Transl. Med. 3, 96ra76 (2011).

12. M. Sirota, J. T. Dudley, J. Kim, A. P. Chiang, A. A. Morgan, A. Sweet-Cordero, J. Sage, A. J. Butte, Discovery and preclinical validation of drug indications using compendia of public gene expression data, Sci. Transl. Med. 3, 96ra77–96ra77 (2011).

13. M. Iskar, G. Zeller, P. Blattmann, M. Campillos, M. Kuhn, K. H. Kaminska, H. Runz, A.-C. Gavin, R. Pepperkok, V. van Noort, P. Bork, Characterization of drug-induced transcriptional modules: towards drug repositioning and functional understanding, Mol. Syst. Biol. 9, 662 (2013).

14. F. Iorio, R. Bosotti, E. Scacheri, V. Belcastro, P. Mithbaokar, R. Ferriero, L. Murino, R. Tagliaferri, N. Brunetti-Pierri, A. Isacchi, D. Di Bernardo, Discovery of drug mode of action and drug repositioning from transcriptional responses, Proc. Natl. Acad. Sci. U. S. A. 107, 14621–14626 (2010).

15. N. El-Hachem, D. M. A. Gendoo, L. S. Ghoraie, Z. Safikhani, P. Smirnov, C. Chung, K. Deng, A. Fang, E. Birkwood, C. Ho, R. Isserlin, G. D. Bader, A. Goldenberg, B. Haibe-Kains, Integrative cancer pharmacogenomics to infer large-scale drug taxonomy, Cancer Res. 77, 3057–3069 (2017).

16. J. A. Parkkinen, S. Kaski, Probabilistic drug connectivity mapping, BMC Bioinformatics 15, 113 (2014).

17. S. Lee, K. H. Lee, M. Song, D. Lee, Building the process-drug–side effect network to discover the relationship between biological Processes and side effects, BMC Bioinformatics 12, S2 (2011).

18. A. Subramanian, R. Narayan, S. M. Corsello, D. D. Peck, T. E. Natoli, X. Lu, J. Gould, J. F. Davis, A. A. Tubelli, J. K. Asiedu, D. L. Lahr, J. E. Hirschman, Z. Liu, M. Donahue, B. Julian, M. Khan, D. Wadden, I. C. Smith, D. Lam, A. Liberzon, C. Toder, M. Bagul, M. Orzechowski, O. M. Enache, F. Piccioni, S. A. Johnson, N. J. Lyons, A. H. Berger, A. F. Shamji, A. N. Brooks, A. Vrcic, C. Flynn, J. Rosains, D. Y. Takeda, R. Hu, D. Davison, J. Lamb, K. Ardlie, L. Hogstrom, P. Greenside, N. S. Gray, P. A. Clemons, S. Silver, X. Wu, W. N. Zhao, W. Read-Button, X. Wu, S. J. Haggarty, L. V. Ronco, J. S. Boehm, S. L. Schreiber, J. G. Doench, J. A. Bittker, D. E. Root, B. Wong, T. R. Golub, A Next Generation Connectivity Map: L1000 Platform and the First 1,000,000 Profiles, Cell 171, 1437–1452.e17 (2017).

19. X. A. Qu, D. K. Rajpal, Applications of Connectivity Map in drug discovery and development, Drug Discov. Today 17, 1289–1298 (2012).

20. A. B. Keenan, M. L. Wojciechowicz, Z. Wang, K. M. Jagodnik, S. L. Jenkins, A. Lachmann, A. Ma’ayan, Connectivity Mapping: Methods and Applications, Annu. Rev. Biomed. Data Sci. 2, 69–92 (2019).

21. M. Iwata, L. Yuan, Q. Zhao, Y. Tabei, F. Berenger, R. Sawada, S. Akiyoshi, M. Hamano, Y. Yamanishi, in Bioinformatics, (Oxford University Press, 2019), vol. 35, pp. i191–i199.

22. I. Vogt, J. Prinz, M. Campillos, Molecularly and clinically related drugs and diseases are enriched in phenotypically similar drug-disease pairs, Genome Med. 6, 1–17 (2014).

23. B. Chen, P. Greenside, H. Paik, M. Sirota, D. Hadley, A. J. Butte, Relating Chemical Structure to Cellular Response: An Integrative Analysis of Gene Expression, Bioactivity, and Structural Data Across 11,000 Compounds, CPT Pharmacometrics Syst. Pharmacol. 4, 576–584 (2015).

24. N. Moret, N. A. Clark, M. Hafner, Y. Wang, E. Lounkine, M. Medvedovic, J. Wang, N. Gray, J. Jenkins, P. K. Sorger, Cheminformatics Tools for Analyzing and Designing Optimized Small-Molecule Collections and Libraries, Cell Chem. Biol. 26, 765–777.e3 (2019).

25. N. R. Clark, K. S. Hu, A. S. Feldmann, Y. Kou, E. Y. Chen, Q. Duan, A. Ma’ayan, The characteristic direction: a geometrical approach to identify differentially expressed genes, BMC Bioinformatics 15, 79 (2014).

26. Q. Duan, S. P. Reid, N. R. Clark, Z. Wang, N. F. Fernandez, A. D. Rouillard, B. Readhead, S. R. Tritsch, R. Hodos, M. Hafner, M. Niepel, P. K. Sorger, J. T. Dudley, S. Bavari, R. G. Panchal, A. Ma’Ayan, L1000CDS2: LINCS L1000 characteristic direction signatures search engine, npj Syst. Biol. Appl. 2, 16015 (2016).

27. M. De Domenico, V. Nicosia, A. Arenas, V. Latora, Structural reducibility of multilayer networks, Nat. Commun. 6, 6864 (2015).

28. S. M. Corsello, J. A. Bittker, Z. Liu, J. Gould, P. McCarren, J. E. Hirschman, S. E. Johnston, A. Vrcic, B. Wong, M. Khan, J. Asiedu, R. Narayan, C. C. Mader, A. Subramanian, T. R. Golub, The Drug Repurposing Hub: A next-generation drug library and information resource Nat. Med. 23, 405–408 (2017).

29. K. C. Cotto, A. H. Wagner, Y. Y. Feng, S. Kiwala, A. C. Coffman, G. Spies, A. Wollam, N. C. Spies, O. L. Griffith, M. Griffith, DGIdb 3.0: A redesign and expansion of the drug-gene interaction database, Nucleic Acids Res. 46, D1068–D1073 (2018).

30. E. March-Vila, L. Pinzi, N. Sturm, A. Tinivella, O. Engkvist, H. Chen, G. Rastelli, On the Integration of In Silico Drug Design Methods for Drug Repurposing, Front. Pharmacol. 0, 298 (2017).

31. Y. Wang, A. Cornett, F. J. King, Y. Mao, F. Nigsch, C. G. Paris, G. McAllister, J. L. Jenkins, Evidence-Based and Quantitative Prioritization of Tool Compounds in Phenotypic Drug Discovery, Cell Chem. Biol. 23, 862–874 (2016).

32. D. Mendez, A. Gaulton, A. P. Bento, J. Chambers, M. De Veij, E. Félix, M. P. Magariños, J. F. Mosquera, P. Mutowo, M. Nowotka, M. Gordillo-Marañón, F. Hunter, L. Junco, G. Mugumbate, M. Rodriguez-Lopez, F. Atkinson, N. Bosc, C. J. Radoux, A. Segura-Cabrera, A. Hersey, A. R. Leach, ChEMBL: towards direct deposition of bioassay data, Nucleic Acids Res. 47, D930–D940 (2019).

33. M. Chartier, L. P. Morency, M. I. Zylber, R. J. Najmanovich, Large-scale detection of drug off-targets: Hypotheses for drug repurposing and understanding side-effects, BMC Pharmacol. Toxicol. 18, 1–16 (2017).

34. B. Haibe-Kains, N. El-Hachem, N. J. Birkbak, A. C. Jin, A. H. Beck, H. J. W. L. Aerts, J. Quackenbush, Inconsistency in large pharmacogenomic studies, Nat. 2013 5047480 504, 389–393 (2013).

35. M. Niepel, M. Hafner, C. E. Mills, K. Subramanian, E. H. Williams, M. Chung, B. Gaudio, A. M. Barrette, A. D. Stern, B. Hu, J. E. Korkola, C. E. Shamu, G. Jayaraman, E. U. Azeloglu, R. Iyengar, E. A. Sobie, G. B. Mills, T. Liby, J. D. Jaffe, M. Alimova, D. Davison, X. Lu, T. R. Golub, A. Subramanian, B. Shelley, C. N. Svendsen, A. Ma’ayan, M. Medvedovic, J. W. Gray, M. R. Birtwistle, L. M. Heiser, P. K. Sorger, A Multi-center Study on the Reproducibility of Drug-Response Assays in Mammalian Cell Lines, Cell Syst. 9, 35–48.e5 (2019).

36. Y. Hu, J. Bajorath, Compound promiscuity: what can we learn from current data?, Drug Discov. Today 18, 644–650 (2013).

37. J. L. Decano, S. A. Singh, C. Gasparotto Bueno, L. Ho Lee, A. Halu, S. Chelvanambi, J. T. Matamalas, H. Zhang, A. K. Mlynarchik, J. Qiao, A. Sharma, S. Mukai, J. Wang, D. G. Anderson, C. K. Ozaki, P. Libby, E. Aikawa, M. Aikawa, Systems Approach to Discovery of Therapeutic Targets for Vein Graft Disease: PPARα Pivotally Regulates Metabolism, Activation, and Heterogeneity of Macrophages and Lesion Development, Circulation 143, 2454–2470 (2021).

38. M. R. De Vries, K. H. Simons, J. W. Jukema, J. Braun, P. H. A. Quax, Vein graft failure: from pathophysiology to clinical outcomes, Nat. Rev. Cardiol. 13, 451–470 (2016).

39. I. Xenogiannis, M. Zenati, D. L. Bhatt, S. V. Rao, J. Rodes-Cabau, S. Goldman, K. A. Shunk, K. Mavromatis, S. Banerjee, K. Alaswad, I. Nikolakopoulos, E. Vemmou, J. Karacsonyi, D. Alexopoulos, M. N. Burke, V. N. Bapat, E. S. Brilakis, Saphenous Vein Graft Failure: From Pathophysiology to Prevention and Treatment Strategies, Circulation 144, 728–745 (2021).

40. A. Halu, J.-G. Wang, H. Iwata, A. Mojcher, A. L. Abib, S. A. Singh, M. Aikawa, A. Sharma, Context-enriched interactome powered by proteomics helps the identification of novel regulators of macrophage activation, Elife 7 (2018), doi:10.7554/eLife.37059.

41. P. Shashkin, B. Dragulev, K. Ley, Macrophage differentiation to foam cells, Curr. Pharm. Des. 11, 3061–3072 (2005).

42. W. Wharton, C. E. Gleason, V. M. Miller, S. Asthana, Rationale and design of the Kronos Early Estrogen Prevention Study (KEEPS) and the KEEPS Cognitive and Affective sub study (KEEPS Cog), Brain Res. 1514, 12–17 (2013).

43. J. Su, X. Cui, Y. Li, H. Mani, G. A. Ferreyra, R. L. Danner, L. L. Hsu, Y. Fitz, P. Q. Eichacker, SB203580, a p38 Inhibitor, Improved Cardiac Function but Worsened Lung Injury and Survival During Escherichia coli Pneumonia in Mice, J. Trauma 68, 1317 (2010).

44. H. Iwata, E. A. Osborn, G. J. Ughi, K. Murakami, C. Goettsch, J. D. Hutcheson, A. Mauskapf, P. C. Mattson, P. Libby, S. A. Singh, J. Matamalas, E. Aikawa, G. J. Tearney, M. Aikawa, F. A. Jaffer, Highly Selective PPARα (Peroxisome Proliferator-Activated Receptor α) Agonist Pemafibrate Inhibits Stent Inflammation and Restenosis Assessed by Multimodality Molecular-Microstructural Imaging, J. Am. Heart Assoc. 10 (2021), doi: 10.1161/JAHA.121.020834.

45. E. Poletto, G. Baldo, Creating cell lines for mimicking diseases, Prog. Mol. Biol. Transl. Sci. 181, 59–87 (2021).

46. M. Niepel, M. Hafner, Q. Duan, Z. Wang, E. O. Paull, M. Chung, X. Lu, J. M. Stuart, T. R. Golub, A. Subramanian, A. Ma’Ayan, P. K. Sorger, Common and cell-type specific responses to anti-cancer drugs revealed by high throughput transcript profiling, Nat. Commun. 2017 81 8, 1–11 (2017).

47. J. Cheng, L. Yang, Comparing gene expression similarity metrics for connectivity map, Proc. - 2013 IEEE Int. Conf. Bioinforma. Biomed. IEEE BIBM 2013, 165–170 (2013).

48. X. Zhou, M. Wang, I. Katsyv, H. Irie, B. Zhang, EMUDRA: Ensemble of Multiple Drug Repositioning Approaches to improve prediction accuracy, Bioinformatics 34, 3151 (2018).

49. J. Cheng, L. Yang, V. Kumar, P. Agarwal, Systematic evaluation of connectivity map for disease indications, Genome Med. 6 (2014), doi:10.1186/S13073-014-0095-1.

50. K. Samart, P. Tuyishime, A. Krishnan, J. Ravi, Reconciling multiple connectivity scores for drug repurposing, Brief. Bioinform. 22, 1–15.

51. S. D. Ghiassian, J. Menche, D. I. Chasman, F. Giulianini, R. Wang, P. Ricchiuto, M. Aikawa, H. Iwata, C. Müller, T. Zeller, A. Sharma, P. Wild, K. Lackner, S. Singh, P. M. Ridker, S. Blankenberg, A. L. Barabási, J. Loscalzo, Endophenotype Network Models: Common Core of Complex Diseases, Sci. Rep. 6 (2016), doi:10.1038/SREP27414.

52. N. Lim, P. Pavlidis, Evaluation of connectivity map shows limited reproducibility in drug repositioning, Sci. Reports 2021 111 11, 1–14 (2021).

53. G. Papadatos, A. Gaulton, A. Hersey, J. P. Overington, Activity, assay and target data curation and quality in the ChEMBL database, J. Comput. Aided. Mol. Des. 29, 885 (2015).

54. A. J. Marcus, M. J. Broekman, J. H. F. Drosopoulos, N. Islam, D. J. Pinsky, C. Sesti, R. Levi, Heterologous cell–cell interactions: thromboregulation, cerebroprotection and cardioprotection by CD39 (NTPDase-1), J. Thromb. Haemost. 1, 2497–2509 (2003).

55. M. V. Kuleshov, M. R. Jones, A. D. Rouillard, N. F. Fernandez, Q. Duan, Z. Wang, S. Koplev, S. L. Jenkins, K. M. Jagodnik, A. Lachmann, M. G. McDermott, C. D. Monteiro, G. W. Gundersen, A. Ma’ayan, Enrichr: a comprehensive gene set enrichment analysis web server 2016 update, Nucleic Acids Res. 44, W90–W97 (2016).

56. A. Liberzon, C. Birger, H. Thorvaldsdóttir, M. Ghandi, J. P. Mesirov, P. Tamayo, The Molecular Signatures Database Hallmark Gene Set Collection, Cell Syst. 1, 417–425 (2015).

57. Y. Wang, S. Zhang, F. Li, Y. Zhou, Y. Zhang, Z. Wang, R. Zhang, J. Zhu, Y. Ren, Y. Tan, C. Qin, Y. Li, X. Li, Y. Chen, F. Zhu, Therapeutic target database 2020: Enriched resource for facilitating research and early development of targeted therapeutics, Nucleic Acids Res. 48, D1031–D1041 (2020).

58. A. Kamburov, K. Pentchev, H. Galicka, C. Wierling, H. Lehrach, R. Herwig, ConsensusPathDB: toward a more complete picture of cell biology, Nucleic Acids Res. 39, D712–D717 (2010).

