## Supplementary Information for "Integrating pharmacogenomics and cheminformatics with diverse disease phenotypes for cell type-guided drug discovery"

*Discussion of drug candidates identified by Pathopticon for vein graft disease*

Rosiglitazone, a PPAR-gamma agonist, (ranked 5^th^ in the early vein graft (VG) disease signature) is an FDA-approved anti-diabetic. The VICTORY trial for anti-atherosclerotic effects found no evidence for a statistically significant effect of rosiglitazone on atherosclerosis (1). RS-100329 (ranked 4^th^ in the late vein graft disease signature) is an α1A-adrenoceptor antagonist. Α1-AR antagonists (norepinephrine) were previously investigated for vein grafts (2), however, not much is known on RS-100329 itself. GW-7647, a PPAR-alpha agonist, is the only drug that appears in the top-10 predictions for both early and late VG time-course data (7th in early, 3rd in late) was previously investigated for atherosclerosis and non-alcoholic steatohepatitis (NASH) (3,4). Estriol (1st in late), is a type of estrogen. Its atheroprotective effects were investigated in the early 2000s. The clinical trial, Estrogen and Graft Atherosclerosis Research (EAGAR), showed no CVD benefit with increased risk of CAD in the remaining healthy coronary vessels (5,6). Betamethasone (1st in early) is a corticosteroid. Several studies on the protective effects of corticosteroids in coronary artery bypass graft surgery were conducted, but mostly on dexamethasone (7,8). Sunitinib (5th in late) is a multitarget RTK inhibitor, and its effect on neointimal hyperplasia in arteriovenous grafts was investigated recently (9). Nitrendipine (8th in early) is a calcium channel blocker (CCB), Improved radial artery outcomes were reported for CCBs (10,11), with nothing in particular for nitrendipine. AG-490 (3^rd^ in early) and Gefitinib (8^th^ in late are EGFR tyrosine kinase inhibitors. There are many studies investigating anti-inflammatory properties in nephropathy and liver ischemia/reperfusion. Improved outcomes were seen in myoblast transplantation. In general, reducing EGFR activity resulted in reduced neointimal hyperplasia (12–15). Pravastatin (9th in late) is an HMG CoA reductase inhibitor, which, like other statins, is prescribed to post-operative CABG patients (16). Recent studies showed protective effects of dephostatin (2nd in late), a protein tyrosine phosphatase 1B inhibitor. against atherosclerotic plaque formation (17), with a possible role in peripheral artery disease (PAD) as well (18). SB-203580 (10th in late), a p38 kinase inhibitor, was Investigated as a potential treatment for VG failure in the early 2000s (19).

While the majority of the top-10 predictions appear to be relevant and with potential for further investigation in the context of vein graft disease, there a few drugs seem to have the opposite effect to what would be expected in the context of vein graft and atherosclerosis, e.g., inhibitors and antagonists when agonists are expected. Splitomycin (2nd in early) is a sirtuin (SIRT1) inhibitor. Sirtuins are believed to be protective against atherosclerosis (20). Calcitriol (9th in early) is the active vitamin D metabolite. Vitamin D is believed to be a promoter of vascular calcification (21). SCH-58261 (10th in early) is an adenosine receptor A2A antagonist. A2AAR is protective against inflammation and appears to have anti-atherosclerotic effects (22–24). The remaining four drugs returned no results relevant to vein graft diseases or atherosclerosis. Most of these drugs are approved for other uses: Loperamide (6th in early), an opioid-receptor agonist, is used as an anti-diarrheal; Formoterol (4th in early), a long-acting beta agonist, is used as a bronchodilator; Sulpiride (6th in late), dopamine/serotonin receptor antagonist, is used as an anti-psychotic; and Paroxetine (7th in late), a selective serotonin reuptake inhibitor, is used as an anti-depressant.

*Supplementary note on adipose stromal cells and macrophages*

Adipose stromal cells (ASCs) have been reported to work in close tandem with macrophages in times of pro-inflammatory activation (25). ASCs can mediate proinflammatory M1-like activation of macrophages (25), while macrophages can inhibit adipogenic differentiation of these cells by producing pro-inflammatory cytokines (26). As a result, adipose stromal cells can differentiate into fibro-inflammatory phenotype when adipogenic differentiation is suppressed (27). Crosstalk between macrophages and ASCs is found in many tissue microenvironments but is particularly important in obesity, a major CAD risk factor. Shared metaflammation pathways between ASCs and macrophages, due to many layers of evolutionarily conserved interactions occurring between immune response and metabolism, may provide key targets for therapeutic intervention in vascular inflammation (28). Hence, a systems approach targeting this relationship may reveal drugs that can access these shared pathways.

**Supplementary Methods**

*Processing the LINCS-CMap Level 4 data*

The LINCS-CMap Level 4 expression data were downloaded from the Touchstone Reference Dataset Supporting Files on the CLUE platform (clue.io) (version date 5/11/2018). This dataset, named “GSE92742_Broad_LINCS_Level4_ZSPCINF_mlr12k_n1319138x12328.gctx.gz” encompasses 1,319,138 perturbation instances (including experimental replicates), 473,647 unique perturbation signatures, 51,383 perturbations, 12,328 genes, and 76 cell lines. The .gctx data matrix file was parsed using the “parse” function (“cmapPy.pandasGEXpress.parse”) within the pandasGEXpress module of the CMapPy tool (29) in Python. The small molecule perturbations (chemical compounds) denoted by “trt_cp” were retained in the data, and the gene (“rid”)-by-perturbation instance (“inst_id”) data tables were written individually to file in the .feather format for each cell type. Out of the 76 cell types in LINCS-CMap Level 4 data, five had no small molecule compounds tested on them, but rather had control vectors or shRNAs used as perturbations. Due to the heterogeneity in the number of perturbations across cell lines **(Supplementary Figure 16)**, we did not consider 10 cell lines with more than 10,000 perturbations (A375: 52,707; A549: 57,775; HA1E: 31,478; HCC515: 26,492; HEPG2: 18,239; HT29: 51,377: MCF7: 99,059; NPC: 14,599; PC3: 91,954; VCAP: 120,468) for downstream analysis. Of the remaining cell lines, MCH58 yielded no significant edges. Overall, this procedure resulted in 60 cell lines with which to carry out the downstream analyses.

*Sensitivity analysis of the QUIZ-C approach*

To assess the response of cell type-specific gene-perturbation network topologies to changes in QUIZ-C parameters, we performed a sensitivity analysis that considers multiple models and consensus thresholds. We considered three models: (i) The core QUIZ-C approach that relies on the instance distribution described in detail in the Main methods. For this model, we considered the consensus thresholds of 70^th^, 75^th^, 80^th^ and 85^th^ quantiles; (ii) Model with an additional “fixed” threshold that divides drugs into two regimes: If the number of instances is over threshold, it calculates quantiles, else if the number of instances is below threshold, a fixed number of significant replicates (e.g., 2) is needed for an edge. For this model, we considered the consensus thresholds of 75^th^ and 80^th^ quantiles, instance thresholds of 3 and 4, and for two significant replicates; (iii) Model with an additional “percentage” threshold that divides it into two regimes: If the number of instances is over threshold, it calculates quantiles, if the number of instances below threshold, a percentage of significant replicates (e.g. 66%) is needed for an edge. For this model, we considered the consensus thresholds of 75^th^ and 80^th^ quantiles, instance thresholds of between 3 and 10, and for 66% significant replicates **(Supplementary Figure 20)**. We found that the second and third models corresponding to fixed and percent thresholds proved limiting for cell lines with fewer instances per drug, and that the quantile cutoff did not have a high impact on the network topologies across different regimes. More than two-thirds of the cell type-specific networks built by QUIZ-C with a quantile cutoff between 70% and 85% had over 75% edge overlap with each other. Based on these findings, we chose the simples model (model i) with a quantile threshold of 75%.

*Mappings between perturbation identifiers*

We used PubChem Compound IDs (CIDs) as the unifying identifier in all analyses to interface various pharmacological and cheminformatics databases. To map the LINCS-CMap perturbation names (“pert_inames”) to their corresponding CIDs, we first extracted the InChI and SMILES keys of all compounds (“trt_cp”) in the LINCS-CMap perturbation dataset (“GSE92742_Broad_LINCS_pert_info.txt”). We then used the InChI and SMILES keys as inputs in the PubChem Identifier Exchange service (<https://pubchem.ncbi.nlm.nih.gov/idexchange/idexchange.cgi>) to retrieve the corresponding CIDs. We merged the results from InChI keys and SMILES to maximize mapping coverage in cases where information about either key was missing. We then subset this global mapping to only include the drugs in our QUIZ-C networks for downstream analyses.

*Construction of MODZ- and CD-based networks for benchmarking*

To benchmark QUIZ-C as a method for gene-perturbation network building, we used two state-of-the-art methods which can determine significant gene-perturbation interactions from the LINCS-CMap data in a similar manner, and can therefore be used to build cell type-specific networks such as ours. The first one of these methods, namely the moderated z-score (MODZ), is the method used in the original LINCS-CMap study (30) and forms the basis of the LINCS-CMap Level 5 data. To build MODZ-based drug-gene networks, we used the differentially expressed genes identified through the MODZ procedure in the LINCS Canvas Browser (31), which we obtained though the Harmonizome database (32) (<https://maayanlab.cloud/Harmonizome/dataset/LINCS+L1000+CMAP+Signatures+of+Differentially+Expressed+Genes+for+Small+Molecules>, retrieved in October 2019). The second method, namely, the characteristic direction (CD) (33), has recently been used to identify differentially expressed genes under perturbations using the LINCS-CMap Level 3 data (34). We downloaded the database of differentially expressed gene signatures calculated using the CD method from the L1000CDS^2^ website (<https://maayanlab.cloud/public/L1000CDS_download/>, retrieved in October, 2019) and extracted the differentially expressed gene signatures for each cell type from the L1000CDS^2^ database using MongoDB. After obtaining the list of up- and down-regulated genes for each perturbation in each cell line for MODZ and CD, we created edge lists for both methods to be used in the construction of the cell type-specific drug-gene networks. Since both MODZ and CD measure differential expression and yield lists of up- and down-genes, they help us build unweighted networks with positive or negative edges similar to our QUIZ-C networks, thereby offering a direct means to benchmark our method and compare the resulting gene-perturbation networks.

*Structural reducibility analysis*

To determine the redundancy in the information content of the LINCS-CMap-based multilayer networks, where each network layer represents a cell type, we used the recently proposed structural reducibility concept (35). An information theoretic measure, structural reducibility $\chi$ quantifies how distinguishable, or non-redundant, each layer of the multilayer network is from a purely topological perspective. We used the Python implementation of the structural reducibility algorithm provided in the public GitHub repository by the authors of the study (<https://github.com/KatolaZ/multired>). To be compatible with the input format requirements of this Python script, we converted all the gene symbols and pert_inames to integer values. We restricted our attention to the 51 layers (51 cell types) that are common to QUIZ-C, MODZ and CD. We followed the simplified analysis workflow via the python script “run_multired_simple.py” and used the resulting Jensen-Shannon divergence matrices and relative entropy quality function $q(\cdot)$ vectors in our in-house scripts to generate hierarchical clusterings of cell types for each network-building method. We extracted the branching point corresponding to the maximum value of $q(\cdot)$ as the optimal configuration of the aggregation of layers (cell types). Using this clustering configuration (denoted by the vertical dashed lines), we calculated the structural reducibility $\chi$ as the ratio of the number of layer aggregation steps (i.e. branching points) to the number of total possible aggregation steps (i.e. the total number of layers minus one)

$$\chi=\frac{M-M_{opt}}{M-1},$$

where $M_{opt}$is the number of branching points at the optimal clustering configuration and $M=51$ is the total number of layers. A maximum at a low value of $q(\cdot)$ represents a highly distinguishable (i.e., not redundant or reducible), whereas a maximum at a high value of $q(\cdot)$ represents a highly redundant (i.e., reducible) multilayer network structure. In our context, this means that the cell type-specific networks are distinct in terms of their information content and cannot be merged without loss of substantial information.

*Single Tandem Repeat (STR) profile similarity*

We first used the cell line identifier and corresponding synonyms information in the Cellosaurus database (36) (Version 34.0, downloaded in March 2020 from <https://web.expasy.org/cellosaurus/>) to map the LINCS-CMap cell line names to Cellosaurus identifiers. Second, we queried Cellosaurus for the following 17 STR markers (excluding Amelogenin): CSF1PO, D2S1338, D3S1358, D5S818, D7S820, D8S1179, D13S317, D16S539, D18S51, D19S433, D21S11, FGA, Penta D, Penta E, TH01, TPOX and vWA. Of the 60 LINCS-CMap cell lines we used as the layers in our QUIZ-C gene-perturbation networks, 54 had STR profile information on Cellosaurus. We then used the Cell Line Authentication using STR (CLASTR) tool (37) (<https://web.expasy.org/cellosaurus-str-search/>) to calculate the STR similarity of these 54 cell lines using the default settings (“Algorithm” set to Tanabe and “Mode” set to non-empty markers). Since CLASTR calculates the STR profile similarity of a given cell line to all the cell lines in Cellosaurus with STR profile information, rather than calculating the pairwise STR similarity of two input cell lines, we obtained results for each cell line without a score filter and for the maximum number of results (typically around 1,000 top-ranked cell). This resulted in a list of cell lines, ranked from the most similar to the least similar, for each cell line. Since Cellosaurus has STR profile information on >6,000 cell lines and our similarity results return at most around 1,000 cell lines, some of the pairwise similarities between the 54 cell lines remained below “detection threshold,” meaning the lowest similarity value in the available CLASTR output. We therefore quantified (i) the similarity scores for the cell lines above the detection threshold (i.e. within the top ~1,000 cell lines), (ii) the similarity score corresponding to the detection threshold and the percentage of cell lines below this detection threshold (i.e. with lower similarity values than the threshold).

*Cell Line Ontology (CLO) semantic similarity*

To measure the ontological similarity of cell lines via the Cell Line Ontology (CLO) (38) (downloaded from [https://bioportal.bioontology.org/ontologies/CLO in January 2020](https://bioportal.bioontology.org/ontologies/CLO%20in%20January%202020)), we used a graph-based semantic similarity approach for biological ontology terms (39). We converted LINCS-CMap cell line identifiers to CLO identifiers via manual curation. We constructed the directed acyclic graph (DAG) from all the terms (cell lines) in CLO using their parent and child terms. We set the semantic contribution factor of parent-child relationship to 0.5 and calculated the S-value $S\left( A \right)$, semantic value $SV\left( A \right)$, and pairwise semantic similarity $S_{CLO}(A, B)$, as described in (39), for all cell lines in L1000.

*Parsing the ChEMBL database for bioactivity records*

In order to assess the selectivity and potency of the compounds in our gene-perturbation networks, we leveraged the large-scale bioactivity database ChEMBL. We downloaded all available bioactivity information from ChEMBL (Release 27, downloaded in June, 2020) and focused on the bioactivity information for compounds that target single proteins in humans by choosing “Homo Sapiens” as the Target Organism and “Single Protein” as the Target Type. These filtering steps resulted in 5,190,464 records out of a total of 16,066,124 bioactivity records in ChEMBL. After removing entries with missing Compound Keys, we converted the molecule and target ChEMBL IDs to PubChem CIDs and UniProt IDs, respectively. Molecule ChEMBL IDs were used as input on the PubChem Identifier Exchange service to retrieve the corresponding PubChem CIDs. Target ChEMBL IDs were converted to UniProt IDs by using the mappings from ChEMBL (“chembl_uniprot_mapping.txt,” downloaded from <ftp://ftp.ebi.ac.uk/pub/databases/chembl/ChEMBLdb/latest/>). The UniProt IDs were then converted to Entrez IDs and approved Gene Symbols using the HUGO Gene Nomenclature Committee website (<https://www.genenames.org/>, downloaded in May 2020). Overall, these ID conversions resulted in 5,107,962 activities between 1,035,402 compounds (in PubChem CIDs) and 3,448 targets (in Gene Symbols/Entrez IDs). Restricting these records to the compounds in the QUIZ-C networks resulted in 235,159 activities between 2,148 QUIZ-C compounds and 1,673 targets **(Supplementary File 4)**.

*Calculating selectivity and tool scores*

To calculate selectivity and tool scores, we focused on nanomolar (nM) activity by restricting the ChEMBL bioactivity data to records that have “nM” as the Standard Unit and “=” as the Standard Relation type, and removed erroneous records with negative nM values. For each drug (CID), we first identified their targets with known nM activity in ChEMBL. Within the set of targets, each target was sequentially taken as the "on-target", and the remaining targets were labeled as the "off-targets." This separation into on- and off-targets enables us to calculate a selectivity and tool score for each drug-target pair with known biomolecular activity. To assess the binding selectivity of a drug to a target, we followed the approaches taken in two recent studies (40,41) and defined selectivity as the combination of three factors (i) magnitude of selectivity (ii) significance of selectivity and (iii) data bias.

First, as a measure of the magnitude of selectivity, we calculated the “potency difference” as the difference between the lower (1st) quartiles of the log_10_(nM) activity distributions of the off- and on-targets, as originally described in (41)

$$Q1_{diff}= Q1(\log_{10} nM_{off-target})- Q1(\log_{10} nM_{on-target}).$$

Second, we calculated the Kolmogorov-Smirnov (KS) test p-value between the log_10_(nM) distributions between off- and on-targets. We introduced a lower threshold of KS p-value at 1e-16 and then calculated the negative logarithm of the KS p-values to serve as a measure of the significance of selectivity

$$selectivity significance= \frac{-\log_{10} KS pvalue}{16}$$

Third, we calculated the data bias as the ratio of the on-target measurements to all measurements for a given drug-target pair

$$data bias= \frac{N_{on-target activities}}{N_{on-target activities}+ N_{off-target activities}}.$$

Given this definition, if there are no off-target measurements, the measurement is severely biased, therefore the data bias is 100%. The higher the number of off-target measurements, the less biased the measurement becomes (i.e. data bias tends to zero). In the Selectivity and Tool Score calculations, we use (1-data bias) to favor drugs with low bias.

Bringing the three components together, we define the selectivity score as defined in (40)

$$selectivity score=\frac{{{Q1}_{diff}}/3+selectivity significance+\left( 1-data bias \right)}{3}.$$

We calculated overall tool scores for each drug-target pair as the product of the selectivity score and strength

$$tool score=selectivity score \times strength$$

where, if both on- and off-targets exist of a given drug exist, we divided binding affinities into three regimes (low affinity: >1000nM; medium affinity: >100 nM and <=1000nM; high affinity: <=100nM) and assigned strength values 7, 4 and 1 if there are more than one high-affinity measurements, if there are more than four medium-affinity measurements, or if neither of these two conditions are met, respectively, following the convention in (40).

We then removed outliers from selectivity score distribution by removing points outside of the range [Q1-4*IQR, Q3+4*IQR]. Since tool scores were skewed by the assigned strength values, we removed the outliers in tool Scores manually by excluding tool Scores above 7.5. We finally scaled the selectivity and tool scores to the range [-1, 1] to be compatible with the pathophenotypic congruity scores. The tool and selectivity scores are provided in **Supplementary File 3**.

*Calculating chemical structure similarity*

We used the cheminformatics toolkit ChemmineR (42) and its annotation package ChemmineDrugs to calculate the chemical structure similarity between drugs. We first downloaded the compound information in the form of structure-data files (SDFs) for QUIZ-C drugs using their PubChem CIDs as inputs. Using the SDFs, we computed the atom pair (AP) descriptors for each compound, which were subsequently used to generate their respective binary atom pair fingerprints (APFPs). The chemical structure similarity between compound pairs were then calculated by comparing the binary APFPs, using the Tanimoto coefficient. These calculations were performed on the O2 High Performance Compute Cluster, supported by the Research Computing Group, at Harvard Medical School. See https://it.hms.harvard.edu/our-services/research-computing for more information.

*Benchmarking datasets*

We used the curated gene sets from the Molecular Signatures Database (MSigDB) (43) as input gene sets for benchmarking. In particular, we used the Chemical and Genetic Perturbations (CGP) data set under the Curated Gene Sets (C2) collection (MSigDB ver 7.1., downloaded June 2020), which mainly consists of gene sets separated into those induced (“up”) and repressed (“down”) by the perturbation. Out of the 3,297 gene sets in CGP, 2,116 were tagged as with either “UP” or “DN,” resulting in 1,058 unique gene sets. Thus, each of these gene sets curated from publications included both “up” and “down” genes for the same study.

We next determined the set of drugs targeting each input gene set from MSigDB using the drug-target interactions from the Therapeutic Target Database (TTD) (44), which were used as true positives for each input gene set in the *benchmark*. We obtained the corresponding CIDs of all drugs with InChI keys in the TTD drug information file (“P1-02-TTD_drug_download.txt”) using the PubChem ID Exchange service, and used this InChI to CID mapping to convert the TTD drug identifiers (starting with “D”) to CIDs. To map the TTD target identifiers (starting with "T") to Gene Symbols, we parsed the TTD target information file (“P1-01-TTD_target_download.txt”) to extract the “GENENAME” tags. Using the TTD drug-target information file (“P1-07-Drug-TargetMapping.xlsx”) and the above two ID mappings we created, we extracted the drugs targeting the genes in each input gene set in terms of CIDs and Gene Symbols, respectively. Finally, we obtained the subset of drugs targeting each gene set that is in the QUIZ-C-, MODZ-, and CD-based networks. Overall, 1,044 drugs in the QUIZ-C-, MODZ-, and CD-based networks (out of the 3,819 drugs in all three sets of networks combined) were used as true positives for the benchmark.

To ensure that we have sufficient true positives in the performance assessment, we used input gene sets that have at least 20 targeting drugs in the QUIZ-C-, MODZ-, and CD-based networks. There were 194 such gene sets out of the 1,058 in the MSigDB CGP data, and we used these as our final benchmark set of input gene sets. We removed the two genesets (“DIAZ_CHRONIC_MEYLOGENOUS_LEUKEMIA” and “DODD_NASOPHARYNGEAL_CARCINOMA”) that resulted in excessively high signature congruity score (SCS) values ($SCS>\left| 0.5 \right|$), which turned out to be also included in the pathophenotype set. We removed these from the benchmark input genesets to ensure orthogonality. The final benchmark set had 192 input signatures **(Supplementary Table 2)**.

As independent datasets to quantify the enrichment of known drug targets in QUIZ-C-, MODZ-, and CD-based networks, we used the Drug Repurposing Hub (45) (<https://www.broadinstitute.org/drug-repurposing-hub>, version date 2018-09-07) and Drug-Gene Interaction Database (DGIdb) (46) (<https://www.dgidb.org/>, accessed April 2019) . We also used the Drug Repurposing Hub to annotate the mechanisms of action (MoA) and the clinical development stage of drugs in our gene-perturbation networks.

*Querying the L1000CDS^2^ database*

We performed automated queries to obtain the top candidates predicted by L1000CDS^2^ for all benchmark signatures through the L1000CDS^2^ web API (https://maayanlab.cloud/L1000CDS2/help/#api). We used the following configuration parameters: "aggravate": (False for Reverse; True for Forward), "searchMethod": "geneSet", "share": False, "combination": False, "db-version": "latest". We note that since the L1000CDS2 server only provides the top 50 results, our comparisons in the benchmark were limited to the top 50 results of each method.

**References**

1.	Bertrand OF, Poirier P, Rodés-Cabau J, Rinfret S, Title LM, Dzavik V, et al. Cardiometabolic effects of rosiglitazone in patients with type 2 diabetes and coronary artery bypass grafts: A randomized placebo-controlled clinical trial. Atherosclerosis [Internet]. 2010 Aug [cited 2022 Nov 17];211(2):565–73. Available from: https://pubmed.ncbi.nlm.nih.gov/20594555/

2.	Erami C, Zhang H, Ho JG, French DM, Faber JE. α1-adrenoceptor stimulation directly induces growth of vascular wall in vivo. Am J Physiol - Hear Circ Physiol [Internet]. 2002 [cited 2022 Nov 17];283(4 52-4):1577–87. Available from: https://journals.physiology.org/doi/10.1152/ajpheart.00218.2002

3.	Nakaya K, Tohyama J, Naik SU, Tanigawa H, MacPhee C, Billheimer JT, et al. Peroxisome proliferator-activated receptor-α activation promotes macrophage reverse cholesterol transport through a liver X receptor-dependent pathway. Arterioscler Thromb Vasc Biol [Internet]. 2011 Jun [cited 2022 Nov 17];31(6):1276–82. Available from: https://pubmed.ncbi.nlm.nih.gov/21441141/

4.	Okishio S, Yamaguchi K, Ishiba H, Tochiki N, Yano K, Takahashi A, et al. PPARα agonist and metformin co-treatment ameliorates NASH in mice induced by a choline-deficient, amino acid-defined diet with 45% fat. Sci Reports 2020 101 [Internet]. 2020 Nov 11 [cited 2022 Nov 17];10(1):1–11. Available from: https://www.nature.com/articles/s41598-020-75805-z

5.	Ali ES, Mangold C, Peiris AN. Estriol: emerging clinical benefits. Menopause [Internet]. 2017 Sep 1 [cited 2022 Nov 17];24(9):1081–5. Available from: https://pubmed.ncbi.nlm.nih.gov/28375935/

6.	Ouyang P, Tardif JC, Herrington DM, Stewart KJ, Thompson PD, Walsh MN, et al. Randomized trial of hormone therapy in women after coronary bypass surgery. Evidence of differential effect of hormone therapy on angiographic progression of disease in saphenous vein grafts and native coronary arteries. Atherosclerosis [Internet]. 2006 Dec [cited 2022 Nov 17];189(2):375–86. Available from: https://pubmed.ncbi.nlm.nih.gov/16442114/

7.	Schepers A, Pires NMM, Eefting D, de Vries MR, van Bockel JH, Quax PHA. Short-term dexamethasone treatment inhibits vein graft thickening in hypercholesterolemic ApoE3Leiden transgenic mice. J Vasc Surg. 2006 Apr 1;43(4):809–15.

8.	Donneyong MM, Kulik A, Gagne JJ. Trends and Patterns of Corticosteroid Use During Coronary Artery Bypass Grafting Surgery in the United States. J Cardiovasc Pharmacol Ther [Internet]. 2018 May 1 [cited 2022 Nov 17];23(3):226–36. Available from: https://pubmed.ncbi.nlm.nih.gov/29258391/

9.	Kwon SH, Li L, He Y, Tey JCS, Li H, Zhuplatov I, et al. Prevention of Venous Neointimal Hyperplasia by a Multitarget Receptor Tyrosine Kinase Inhibitor. J Vasc Res [Internet]. 2015 Mar 1 [cited 2022 Nov 17];52(4):244–56. Available from: https://pubmed.ncbi.nlm.nih.gov/26788996/

10.	Hall AB, Brilakis ES. Saphenous vein graft failure: seeing the bigger picture. J Thorac Dis [Internet]. 2019 [cited 2022 Nov 17];11(Suppl 9):S1441. Available from: /pmc/articles/PMC6560559/

11.	Gaudino M, Benedetto U, Fremes S, Hare DL, Hayward P, Moat N, et al. Effect of Calcium-Channel Blocker Therapy on Radial Artery Grafts After Coronary Bypass Surgery. J Am Coll Cardiol [Internet]. 2019 May 14 [cited 2022 Nov 17];73(18):2299–306. Available from: https://pubmed.ncbi.nlm.nih.gov/31072574/

12.	Gérard C, Dufour C, Goudenege S, Skuk D, Tremblay JP. AG490 improves the survival of human myoblasts in vitro and in vivo. Cell Transplant [Internet]. 2012 [cited 2022 Nov 17];21(12):2665–76. Available from: https://pubmed.ncbi.nlm.nih.gov/22963730/

13.	Freitas MCS, Uchida Y, Zhao D, Ke B, Busuttil RW, Kupiec-Weglinski JW. Blockade of Janus kinase-2 signaling ameliorates mouse liver damage due to ischemia and reperfusion. Liver Transpl [Internet]. 2010 May [cited 2022 Nov 17];16(5):600–10. Available from: https://pubmed.ncbi.nlm.nih.gov/20440769/

14.	Chan AK, Kalmes A, Hawkins S, Daum G, Clowes AW. Blockade of the epidermal growth factor receptor decreases intimal hyperplasia in balloon-injured rat carotid artery. J Vasc Surg [Internet]. 2003 Mar 1 [cited 2022 Nov 17];37(3):644–9. Available from: http://www.jvascsurg.org/article/S0741521402752336/fulltext

15.	Trieu VN, Narla RK, Myers DE, Uckun FM. EGF-genistein inhibits neointimal hyperplasia after vascular injury in an experimental restenosis model. J Cardiovasc Pharmacol [Internet]. 2000 [cited 2022 Nov 17];35(4):595–605. Available from: https://pubmed.ncbi.nlm.nih.gov/10774791/

16.	Makuuchi H, Furuse A, Endo M, Nakamura H, Daida H, Watanabe M, et al. Effect of pravastatin on progression of coronary atherosclerosis in patients after coronary artery bypass surgery. Circ J [Internet]. 2005 Jun [cited 2022 Nov 17];69(6):636–43. Available from: https://pubmed.ncbi.nlm.nih.gov/15914938/

17.	Thompson D, Morrice N, Grant L, Le Sommer S, Lees EK, Mody N, et al. Pharmacological inhibition of protein tyrosine phosphatase 1B protects against atherosclerotic plaque formation in the LDLR−/− mouse model of atherosclerosis. Clin Sci (Lond) [Internet]. 2017 Oct 10 [cited 2022 Nov 17];131(20):2489. Available from: /pmc/articles/PMC6365594/

18.	Mercier C, Rousseau M, Geraldes P. Growth Factor Deregulation and Emerging Role of Phosphatases in Diabetic Peripheral Artery Disease. Front Cardiovasc Med. 2021 Jan 7;7:380.

19.	Cornelissen J, Armstrong J, Holt CM. Mechanical Stretch Induces Phosphorylation of p38-MAPK and Apoptosis in Human Saphenous Vein. Arterioscler Thromb Vasc Biol [Internet]. 2004 Mar 1 [cited 2022 Nov 17];24(3):451–6. Available from: https://www.ahajournals.org/doi/abs/10.1161/01.ATV.0000116690.17017.8b

20.	Grootaert MOJ, Bennett MR. Sirtuins in atherosclerosis: guardians of healthspan and therapeutic targets. Nat Rev Cardiol 2022 1910 [Internet]. 2022 Mar 30 [cited 2022 Nov 28];19(10):668–83. Available from: https://www.nature.com/articles/s41569-022-00685-x

21.	Razzaque MS. The dualistic role of vitamin D in vascular calcifications. Kidney Int [Internet]. 2011 [cited 2022 Nov 28];79(7):708. Available from: /pmc/articles/PMC3120050/

22.	Johnston-Cox HA, Koupenova M, Ravid K. A2 Adenosine Receptors and Vascular Pathologies. Arterioscler Thromb Vasc Biol [Internet]. 2012 Apr [cited 2022 Nov 28];32(4):870. Available from: /pmc/articles/PMC5755359/

23.	Wang H, Zhang W, Tang R, Zhu C, Bucher C, Blazar BR, et al. Adenosine receptor A2a deficiency in leukocytes increases arterial neointima formation in apolipoprotein E-deficient mice. Arterioscler Thromb Vasc Biol [Internet]. 2010 May 1 [cited 2022 Nov 28];30(5):915–22. Available from: https://www.ahajournals.org/doi/abs/10.1161/ATVBAHA.109.202572

24.	Zernecke A, Bidzhekov K, Özüyaman B, Fraemohs L, Liehn EA, Lüscher-Firzlaff JM, et al. CD73/Ecto-5′-Nucleotidase Protects Against Vascular Inflammation and Neointima Formation. Circulation [Internet]. 2006 May 2 [cited 2022 Nov 28];113(17):2120–7. Available from: https://www.ahajournals.org/doi/abs/10.1161/CIRCULATIONAHA.105.595249

25.	Manferdini C, Paolella F, Gabusi E, Gambari L, Piacentini A, Filardo G, et al. Adipose stromal cells mediated switching of the pro-inflammatory profile of M1-like macrophages is facilitated by PGE2: in vitro evaluation. Osteoarthr Cartil [Internet]. 2017 Jul 1 [cited 2022 Dec 18];25(7):1161–71. Available from: https://pubmed.ncbi.nlm.nih.gov/28153787/

26.	Ma H, Li YN, Song L, Liu R, Li X, Shang Q, et al. Macrophages inhibit adipogenic differentiation of adipose tissue derived mesenchymal stem/stromal cells by producing pro-inflammatory cytokines. Cell Biosci [Internet]. 2020 Jul 20 [cited 2022 Dec 18];10(1). Available from: https://pubmed.ncbi.nlm.nih.gov/32699606/

27.	Joffin N, Paschoal VA, Gliniak CM, Crewe C, Elnwasany A, Szweda LI, et al. Mitochondrial metabolism is a key regulator of the fibro-inflammatory and adipogenic stromal subpopulations in white adipose tissue. Cell Stem Cell [Internet]. 2021 Apr 1 [cited 2022 Dec 18];28(4):702-717.e8. Available from: https://pubmed.ncbi.nlm.nih.gov/33539722/

28.	Hotamisligil GS. Inflammation, metaflammation and immunometabolic disorders. Nature [Internet]. 2017 Feb 8 [cited 2022 Dec 18];542(7640):177–85. Available from: https://pubmed.ncbi.nlm.nih.gov/28179656/

29.	Enache OM, Lahr DL, Natoli TE, Litichevskiy L, Wadden D, Flynn C, et al. The GCTx format and cmap{Py, R, M, J} packages: Resources for optimized storage and integrated traversal of annotated dense matrices. Bioinformatics [Internet]. 2019 Apr 15 [cited 2020 Oct 29];35(8):1427–9. Available from: www.hdfgroup.org/HDF5/.

30.	Subramanian A, Narayan R, Corsello SM, Peck DD, Natoli TE, Lu X, et al. A Next Generation Connectivity Map: L1000 Platform and the First 1,000,000 Profiles. Cell [Internet]. 2017 Nov 30 [cited 2020 Oct 29];171(6):1437-1452.e17. Available from: /pmc/articles/PMC5990023/?report=abstract

31.	Duan Q, Flynn C, Niepel M, Hafner M, Muhlich JL, Fernandez NF, et al. LINCS Canvas Browser: Interactive web app to query, browse and interrogate LINCS L1000 gene expression signatures. Nucleic Acids Res [Internet]. 2014 Jul 1 [cited 2020 Oct 31];42(W1). Available from: https://pubmed.ncbi.nlm.nih.gov/24906883/

32.	Rouillard AD, Gundersen GW, Fernandez NF, Wang Z, Monteiro CD, McDermott MG, et al. The harmonizome: a collection of processed datasets gathered to serve and mine knowledge about genes and proteins. Database (Oxford) [Internet]. 2016 Jan 1 [cited 2020 Oct 31];2016. Available from: https://academic.oup.com/database/article/doi/10.1093/database/baw100/2630482

33.	Clark NR, Hu KS, Feldmann AS, Kou Y, Chen EY, Duan Q, et al. The characteristic direction: a geometrical approach to identify differentially expressed genes. BMC Bioinformatics [Internet]. 2014 Mar 21 [cited 2020 Oct 31];15(1):79. Available from: http://bmcbioinformatics.biomedcentral.com/articles/10.1186/1471-2105-15-79

34.	Duan Q, Reid SP, Clark NR, Wang Z, Fernandez NF, Rouillard AD, et al. L1000CDS2: LINCS L1000 characteristic direction signatures search engine. npj Syst Biol Appl [Internet]. 2016 Aug 4 [cited 2020 Oct 31];2(1):16015. Available from: http://www.nature.com/articles/npjsba201615

35.	De Domenico M, Nicosia V, Arenas A, Latora V. Structural reducibility of multilayer networks. Nat Commun [Internet]. 2015 Apr 23 [cited 2020 Dec 17];6(1):6864. Available from: http://www.nature.com/articles/ncomms7864

36.	Bairoch A. The cellosaurus, a cell-line knowledge resource. J Biomol Tech [Internet]. 2018 Jul 1 [cited 2020 Dec 20];29(2):25–38. Available from: /pmc/articles/PMC5945021/?report=abstract

37.	Robin T, Capes‐Davis A, Bairoch A. CLASTR: The Cellosaurus STR similarity search tool ‐ A precious help for cell line authentication. Int J Cancer [Internet]. 2020 Mar 4 [cited 2020 Dec 20];146(5):1299–306. Available from: https://onlinelibrary.wiley.com/doi/abs/10.1002/ijc.32639

38.	Sarntivijai S, Lin Y, Xiang Z, Meehan TF, Diehl AD, Vempati UD, et al. CLO: The cell line ontology. J Biomed Semantics [Internet]. 2014 Aug 13 [cited 2020 Dec 20];5(1):37. Available from: /pmc/articles/PMC4387853/?report=abstract

39.	Wang JZ, Du Z, Payattakool R, Yu PS, Chen C-F. A new method to measure the semantic similarity of GO terms. Bioinformatics [Internet]. 2007 May 15 [cited 2020 Dec 20];23(10):1274–81. Available from: https://academic.oup.com/bioinformatics/article-lookup/doi/10.1093/bioinformatics/btm087

40.	Moret N, Clark NA, Hafner M, Wang Y, Lounkine E, Medvedovic M, et al. Cheminformatics Tools for Analyzing and Designing Optimized Small-Molecule Collections and Libraries. Cell Chem Biol [Internet]. 2019 May 16 [cited 2020 Nov 1];26(5):765-777.e3. Available from: https://doi.org/10.1016/j.chembiol.2019.02.018

41.	Wang Y, Cornett A, King FJ, Mao Y, Nigsch F, Paris CG, et al. Evidence-Based and Quantitative Prioritization of Tool Compounds in Phenotypic Drug Discovery. Cell Chem Biol [Internet]. 2016 Jul 21 [cited 2020 Nov 1];23(7):862–74. Available from: http://dx.doi.org/10.1016/j.chembiol.2016.05.016http://dx.doi.org/10.1016/j.chembiol.2016.05.016

42.	Cao Y, Charisi A, Cheng LC, Jiang T, Girke T. ChemmineR: A compound mining framework for R. Bioinformatics [Internet]. 2008 Aug [cited 2020 Dec 17];24(15):1733–4. Available from: /pmc/articles/PMC2638865/?report=abstract

43.	Liberzon A, Birger C, Thorvaldsdóttir H, Ghandi M, Mesirov JP, Tamayo P. The Molecular Signatures Database Hallmark Gene Set Collection. Cell Syst [Internet]. 2015 Dec 23 [cited 2018 May 22];1(6):417–25. Available from: http://www.ncbi.nlm.nih.gov/pubmed/26771021

44.	Wang Y, Zhang S, Li F, Zhou Y, Zhang Y, Wang Z, et al. Therapeutic target database 2020: Enriched resource for facilitating research and early development of targeted therapeutics. Nucleic Acids Res [Internet]. 2020 Jan 1 [cited 2020 Dec 13];48(D1):D1031–41. Available from: http://db.idrblab.net/ttd/

45.	Corsello SM, Bittker JA, Liu Z, Gould J, McCarren P, Hirschman JE, et al. The Drug Repurposing Hub: A next-generation drug library and information resource [Internet]. Vol. 23, Nature Medicine. Nature Publishing Group; 2017 [cited 2020 Dec 19]. p. 405–8. Available from: https://www.ncbi.nlm.nih.gov/pmc/articles/PMC5568558/

46.	Cotto KC, Wagner AH, Feng YY, Kiwala S, Coffman AC, Spies G, et al. DGIdb 3.0: A redesign and expansion of the drug-gene interaction database. Nucleic Acids Res [Internet]. 2018 Jan 1 [cited 2020 Dec 19];46(D1):D1068–73. Available from: https://academic.oup.com/nar/article/46/D1/D1068/4634012

**Supplementary Figure Legends:**

**Supplementary Figure 1:** The number of perturbation instances for each cell line in the LINCS-CMap data.

**Supplementary Figure 2:** Overview of QUIZ-C cell type-specific gene-perturbation networks. Green and yellow nodes indicate perturbations and genes, respectively. Up- and down-regulating interactions are shown in red and blue edges, respectively. The positions of the drugs (green nodes) are fixed across all networks. Instructions on how to access the zoomable and interactive versions of each QUIZ-C network can be found at <https://github.com/r-duh/Pathopticon>.

**Supplementary Figure 3:** **(A)** The number of cell type-specific MODZ networks a given drug appears in, as a function of the mean normalized out-degree of each drug across all MODZ networks. Every circle represents a drug, and circle size is proportional to the standard deviation of the normalized out-degree. **(B)** The number of cell type-specific MODZ networks a given gene appears in, as a function of the mean normalized in-degree of each gene across all MODZ networks. Every circle represents a gene, and the circle size is proportional to the standard deviation of the normalized in-degree. **(C)** The number of cell type-specific CD networks a given drug appears in, as a function of the mean normalized out-degree of each drug across all CD networks. Every circle represents a drug, and circle size is proportional to the standard deviation of the normalized out-degree. **(D)** The number of cell type-specific CD networks a given gene appears in, as a function of the mean normalized in-degree of each gene across all CD networks. Every circle represents a gene, and the circle size is proportional to the standard deviation of the normalized in-degree.

**Supplementary Figure 4:** **(A)** Hierarchically clustered heatmap qualitatively showing the edge overlap between MODZ networks. Each row and column correspond to a cell line. The colors correspond to the Jaccard index of edge overlap. **(B)** Hierarchically clustered heatmap qualitatively showing the edge overlap between CD networks. Each row and column correspond to a cell line. The colors correspond to the Jaccard index of edge overlap. **(C)** (Upper panel) Full boxplots showing the edge overlap Jaccard index of each cell line with all other cell lines. Green, red, and blue correspond to QUIZ-C, MODZ and CD networks, respectively. (Lower panel) Zoomed in version of the boxplot without fliers. **(D)** Edge density of CMap-based networks. Green, red, and blue correspond to QUIZ-C, MODZ and CD networks. **(E)** Boxplots showing percent edge concordance (i.e., the percentage of overlapping edges with the same direction of effect) of each cell line with respect to the other cell lines. Green, red, and blue correspond to QUIZ-C, MODZ and CD networks, respectively.

**Supplementary Figure 5:** **(A)** Reducibility of CD networks. **(B)** Reducibility of QUIZ-C networks. Left panel shows the relative entropy quality function $q(\cdot)$ of each hierarchical clustering branch, with the maximum value of $q(\cdot)$ corresponding to the optimal configuration of the aggregation of layers. Right panel shows the hierarchical clustering dendrogram and the optimal clustering threshold. Cell lines belonging to the same cluster are in the same color, and clusters consisting of only one cell line are shown in light grey.

**Supplementary Figure 6:** **(A)** Hierarchically clustered heatmap showing the single Tandem Repeat (STR) profile similarity between LINCS-CMap cell lines, calculated by CLASTR. **(B)** (Top) Boxplots of STR similarities showing the similarity values above the detection threshold. (Bottom) The similarity score corresponding to the detection threshold (red lines) and the percentage of cell lines that fall below this detection threshold (grey bars).

**Supplementary Figure 7:** **(A)** Hierarchically clustered heatmap showing the Cell Line Ontology (CLO) similarity between LINCS-CMap cell lines. **(B)** Boxplots of CLO similarities of each cell line with respect to the other LINCS-CMap cell lines.

**Supplementary Figure 8:** Percentage of drugs in CMap-based networks that have known mechanisms of action per the Drug Repurposing Hub. Green, red, and blue circles correspond to QUIZ-C, MODZ and CD networks, respectively.

**Supplementary Figure 9: (A)** Clinical development phase breakdown of the drugs in each MODZ network. **(B)** Clinical development phase breakdown of the drugs in each CD network. **(C)** The number of cell type-specific MODZ networks a given drug appears in for experimental, investigational and approved drugs. **(D)** The mean normalized out-degree of a given drug across MODZ networks, for experimental, investigational, and approved drugs. **(E)** The number of cell type-specific CD networks a given drug appears in for experimental, investigational, and approved drugs. **(F)** The mean normalized out-degree of a given drug across CD networks, for experimental, investigational and approved drugs. Reported p-values in (C)-(F) are from two-sided Mann-Whitney U tests.

**Supplementary Figure 10: (A)** The percentage of drugs in QUIZ-C networks with known targets in the Drug Repurposing Hub (DRH) and Drug-Gene Interaction Database (DGIdb). Light green indicates percentage in DRH only and dark green indicates the additions from DGIdb. **(B)** The odds ratio of enrichment of known drug-target interactions in each MODZ network. Error bars indicate 95% confidence intervals and cell lines in red indicate a significant enrichment. **(C)** The odds ratio of enrichment of known drug-target interactions in each CD network. Error bars indicate 95% confidence intervals and cell lines in red indicate a significant enrichment. **(D)** The difference of log odds-ratios between QUIZ-C and MODZ networks. Green and red denote cell lines for which QUIZ-C and MODZ have a higher log odds-ratio, respectively. **(E)** The difference of log odds-ratios between QUIZ-C and CD networks. Green and blue denote cell lines for which QUIZ-C and CD have a higher log odds-ratio, respectively.

**Supplementary Figure 11:** Circle heatmaps showing the AUROC values of benchmark signatures (rows) for each cell line (columns). Circle size reflects AUROC value and cell lines marked in darker shade are the significant cases for which empirical p-value < 0.05. Bars at the top show the total number of significant benchmark signatures per cell line and bars on the right show the total number of significant cell lines per benchmark signature. **(A)** PACOS in repress mode. **(B)** PACOS in enhance mode.

**Supplementary Figure 12: (A)** The total number of drugs in the top 50 predictions made by PACOS and PACOS combined with tool score, as a function of the PACOS-to-tool-score-ratio $r$. **(B)** The AUROC for PACOS and PACOS combined with tool score, as a function of the PACOS-to-tool-score-ratio $r$.

**Supplementary Figure 13:** Circle heatmaps showing the AUROC values of benchmark signatures (rows) for each cell line (columns). Circle size reflects AUROC value and cell lines marked in darker shade are the significant cases for which empirical p-value < 0.05. Bars at the top show the total number of significant benchmark signatures per cell line and bars on the right show the total number of significant cell lines per benchmark signature. **(A)** PACOS combined with tool score, in repress mode. **(B)** PACOS combined with tool score, in enhance mode.

**Supplementary Figure 14:** Boxplots of the chemical similarity of the top 50 predicted drugs for each benchmark signature. Each boxplot contains data on the similarity of the given cell line with all other cell lines. Chemical similarity is defined as the Tanimoto coefficient of similarity between the atom pair fingerprints (APFPs) of the top predicted drugs. **(A)** PACOS combined with tool score in repress mode, run on QUIZ-C networks. Font colors indicate comparisons of panel (A) with panels (B) and (C): Green, red, and blue font colors indicate the benchmark signatures for which QUIZ-C top candidates had lower chemical similarity (i.e., higher chemical diversity) than the top candidates of both MODZ and CD networks, and only CD networks, respectively. **(B)** PACOS combined with tool score in repress mode, run on MODZ networks. **(C)** PACOS combined with tool score in repress mode, run on CD networks.

**Supplementary Figure 15:** Boxplots of the chemical similarity of the top 50 predicted drugs for each benchmark signature. Each boxplot contains data on the similarity of the given cell line with all other cell lines. Chemical similarity is defined as the Tanimoto coefficient of similarity between the atom pair fingerprints (APFPs) of the top predicted drugs. **(A)** PACOS combined with tool score in enhance mode, run on QUIZ-C networks. Font colors indicate comparisons of panel (A) with panels (B) and (C): Green, red, and blue font colors indicate the benchmark signatures for which QUIZ-C top candidates had lower chemical similarity (i.e., higher chemical diversity) than the top candidates of both MODZ and CD networks, only MODZ networks, and only CD networks, respectively. **(B)** PACOS combined with tool score in enhance mode, run on MODZ networks. **(C)** PACOS combined with tool score in enhance mode, run on CD networks.

**Supplementary Figure 16:** Boxplots of the chemical similarity of the top 50 predicted drugs for each benchmark signature. Each boxplot contains data on the similarity of the given cell line with all other cell lines. Chemical similarity is defined as the Tanimoto coefficient of similarity between the atom pair fingerprints (APFPs) of the top predicted drugs. **(A)** PACOS combined with tool score in repress mode, run on QUIZ-C networks. Font colors indicate comparisons between panels (A) and (B): Green font color indicates the benchmark signatures for which QUIZ-C top candidates had lower chemical similarity (i.e., higher chemical diversity) than the top candidates of L1000CDS2. **(B)** L1000CDS2 predictions. Font colors indicate comparisons between panels (A) and (B): Magenta font color indicates the benchmark signatures for which L1000CDS2 top candidates had lower chemical similarity (i.e., higher chemical diversity) than the top candidates of PACOS combined with tool score in repress mode, run on QUIZ-C networks.

**Supplementary Figure 17:** Boxplots of the chemical similarity of the top 50 predicted drugs for each benchmark signature. Each boxplot contains data on the similarity of the given cell line with all other cell lines. Chemical similarity is defined as the Tanimoto coefficient of similarity between the atom pair fingerprints (APFPs) of the top predicted drugs. **(A)** PACOS combined with tool score in enhance mode, run on QUIZ-C networks. Font colors indicate comparisons between panels (A) and (B): Green font color indicates the benchmark signatures for which QUIZ-C top candidates had lower chemical similarity (i.e., higher chemical diversity) than the top candidates of L1000CDS2. **(B)** L1000CDS2 predictions. Font colors indicate comparisons between panels (A) and (B): Magenta font color indicates the benchmark signatures for which L1000CDS2 top candidates had lower chemical similarity (i.e., higher chemical diversity) than the top candidates of PACOS combined with tool score in enhance mode, run on QUIZ-C networks.

**Supplementary Figure 18:** q(PCR) results of the in vitro validation candidate drugs for the selected panel of pro-inflammatory/athero-genic genes.

**Supplementary Figure 19:** q(PCR) results of the in vitro validation candidate drugs for the selected panel of anti-inflammatory/athero-protective genes.

**Supplementary Figure 20:** Sensitivity of QUIZ-C networks to the choice of quantiles and other network building parameters. Similarity between network topologies is quantified by the Jaccard coefficient of overlap. All comparisons are made with the network configuration used in the manuscript, i.e., a 75^th^ quantile consensus threshold.

**Supplementary Table 1:** Properties of the QUIZ-C cell type-specific gene-perturbation networks.

**Supplementary Table 2:** Alphabetical list of benchmark signatures.

**Supplementary Table 3:** Comparison of methods in terms of their chemical structure similarity, measured by the Tanimoto coefficient.

**Supplementary Table 4:** Top therapeutic candidates to repress the early and late stage signatures of vein graft disease (VGD), predicted by pathopticon. Predictions that are significant for a given cell line (empirical p-value <=0.05) are marked in red. Lists are truncated after the top 10 significant hits.

**Supplementary File 1:** The pathway enrichment (nominal p-values) of the top five disease phenotypes with the highest signature overlap with the vein graft disease signature (i.e., the leftmost and rightmost diseases on the x-axis).

**Supplementary File 2:** The pathway enrichment (FDR values) of the top five disease phenotypes with the highest signature overlap with the vein graft disease signature (i.e., the leftmost and rightmost diseases on the x-axis).

**Supplementary File 3:** The tool and selectivity scores of the compounds in QUIZ-C networks.

**Supplementary File 4:** The ChEMBL bioactivities between the compounds and targets in QUIZ-C networks.
