## Supplementary Figures and Tables for "Integrating pharmacogenomics and cheminformatics with diverse disease phenotypes for cell type-guided drug discovery"

### Supplementary Figure 1

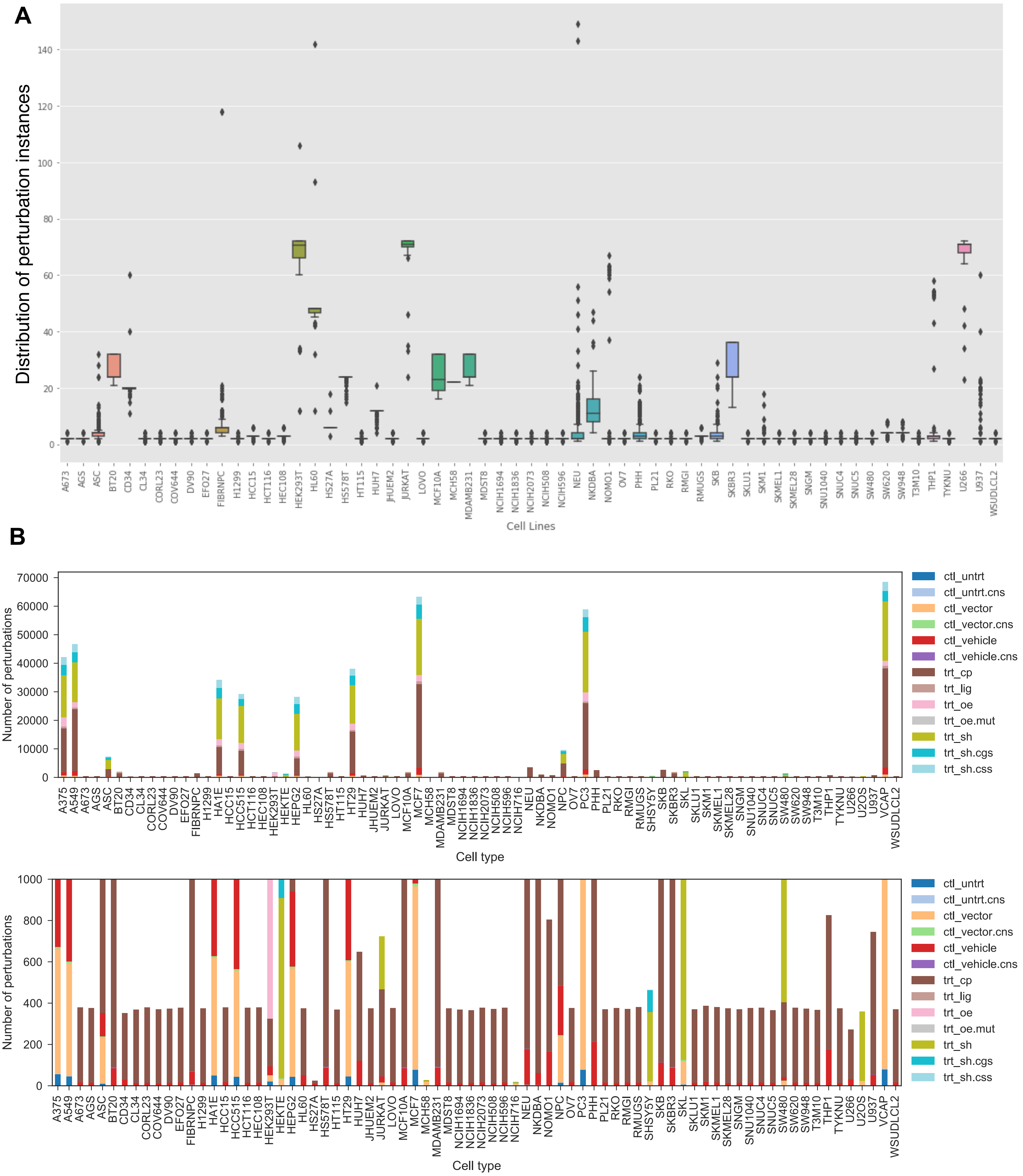

### Supplementary Figure 2

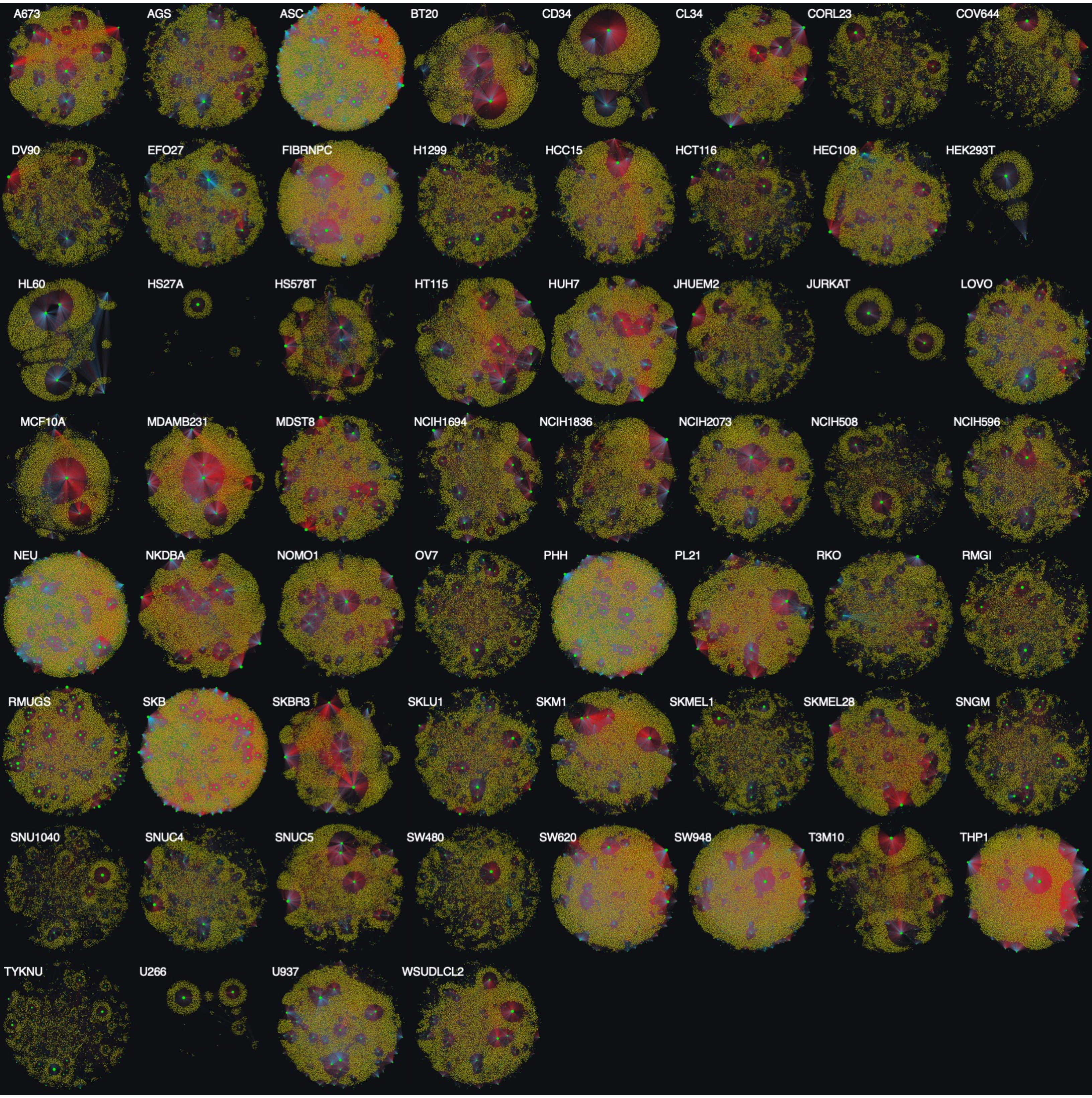

### Supplementary Figure 3

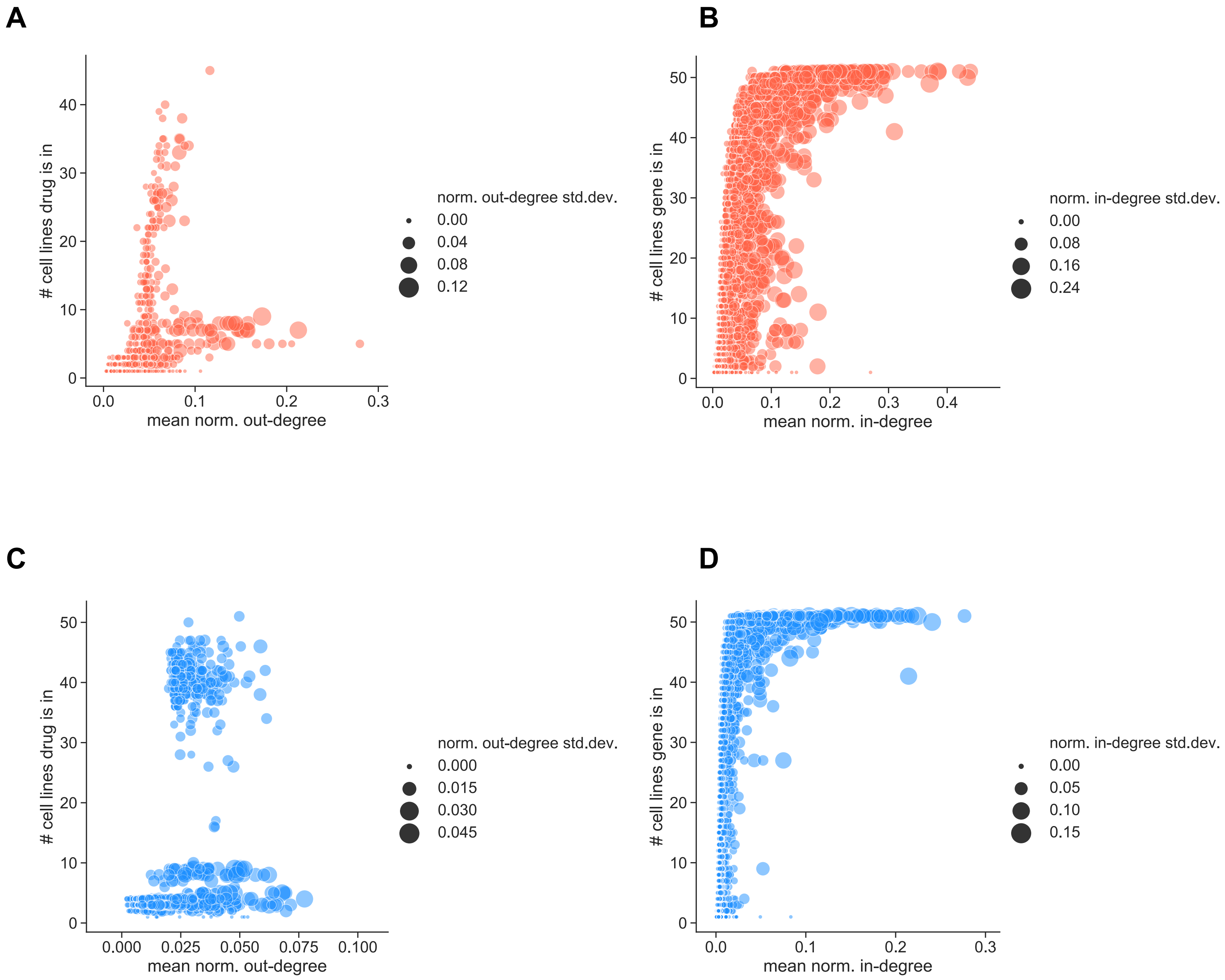

### Supplementary Figure 4

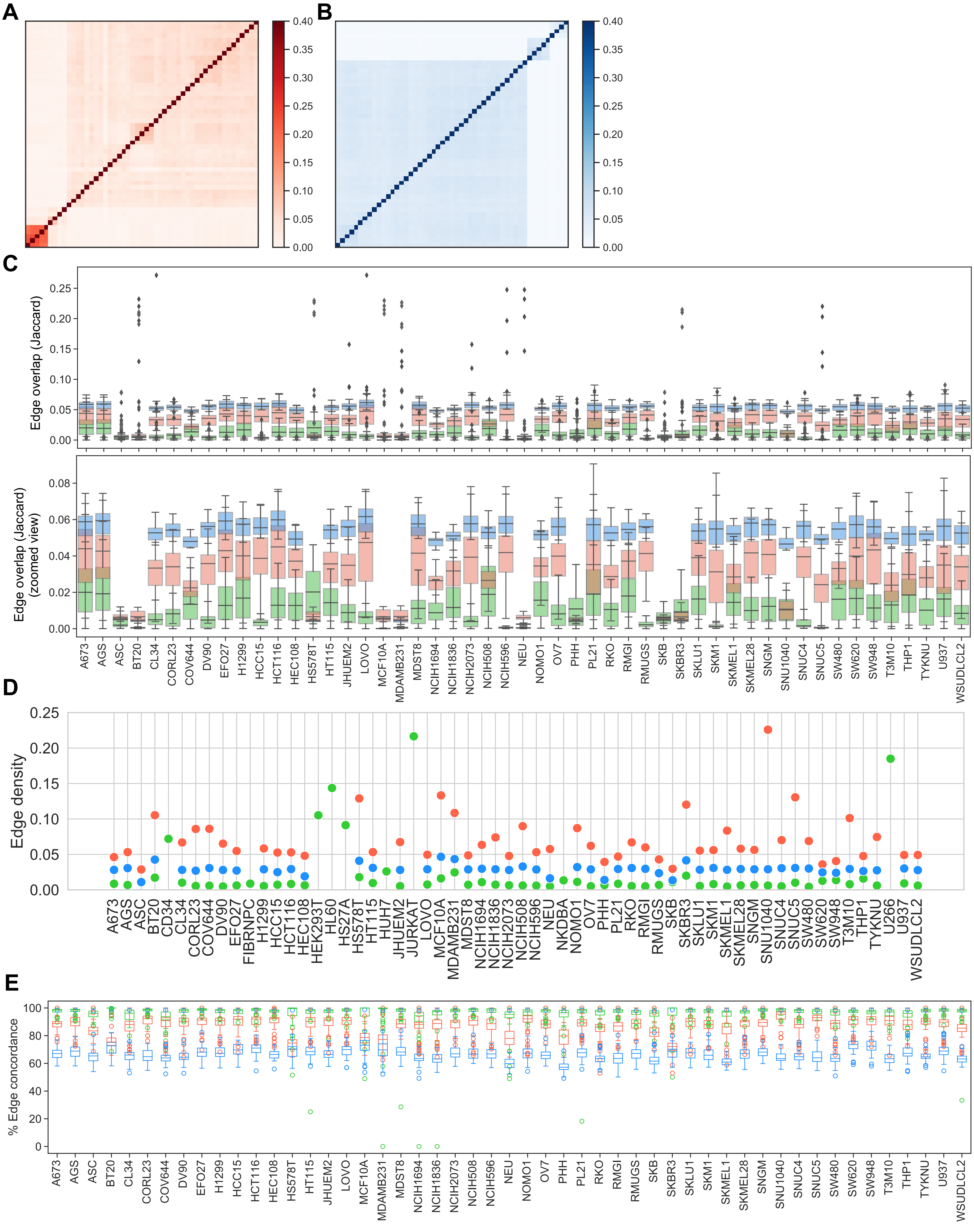

### Supplementary Figure 5

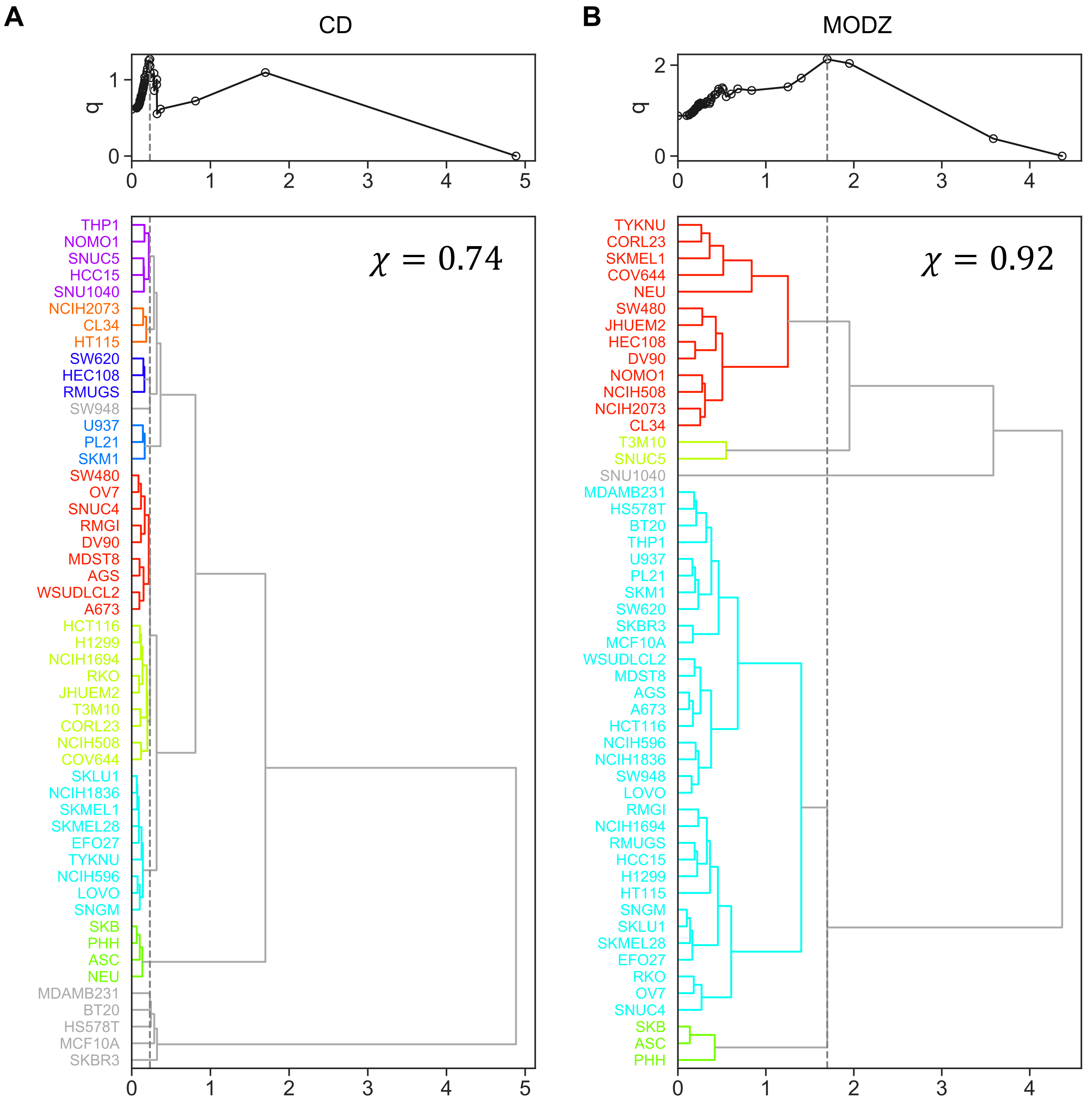

### Supplementary Figure 6

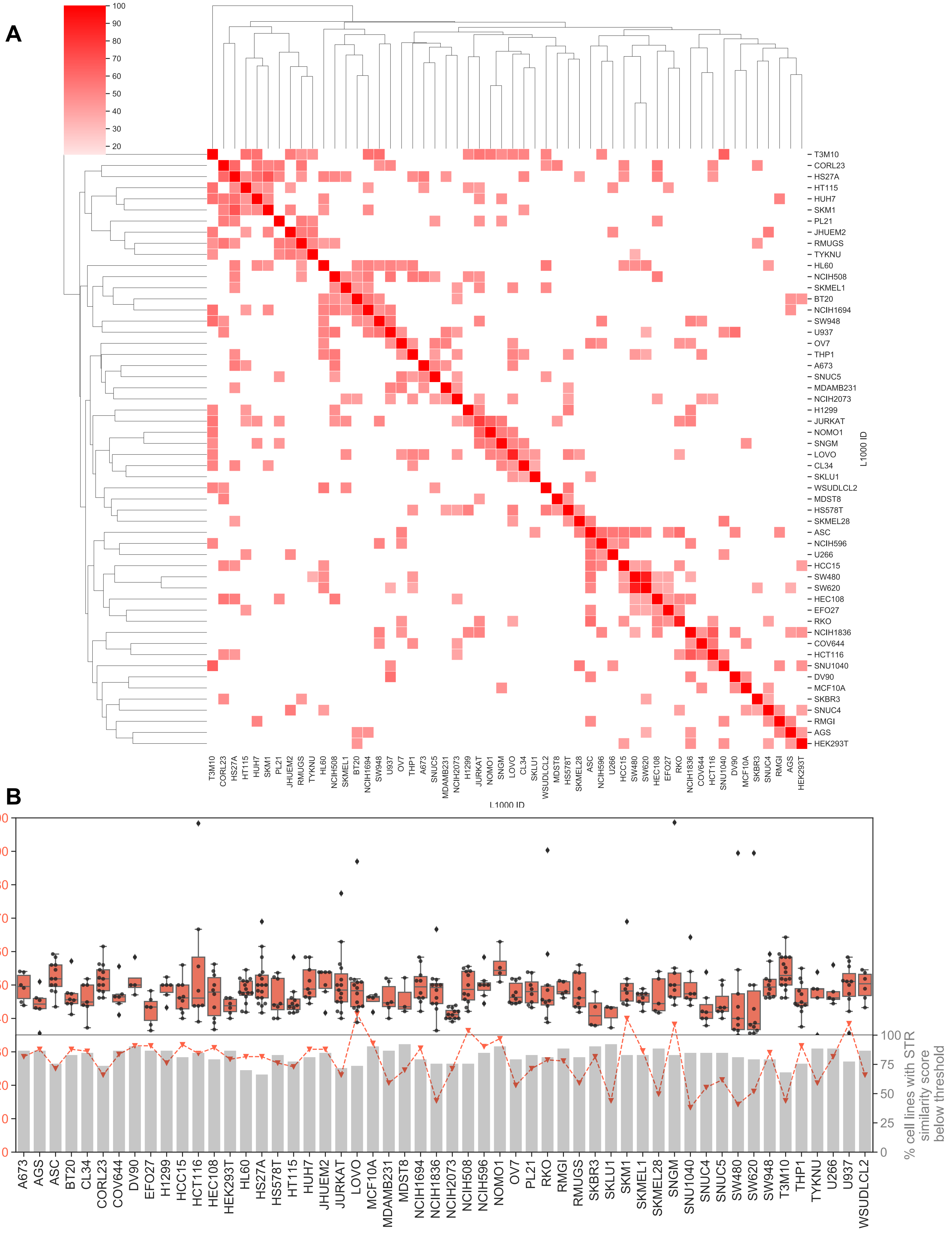

### Supplementary Figure 7

**A**

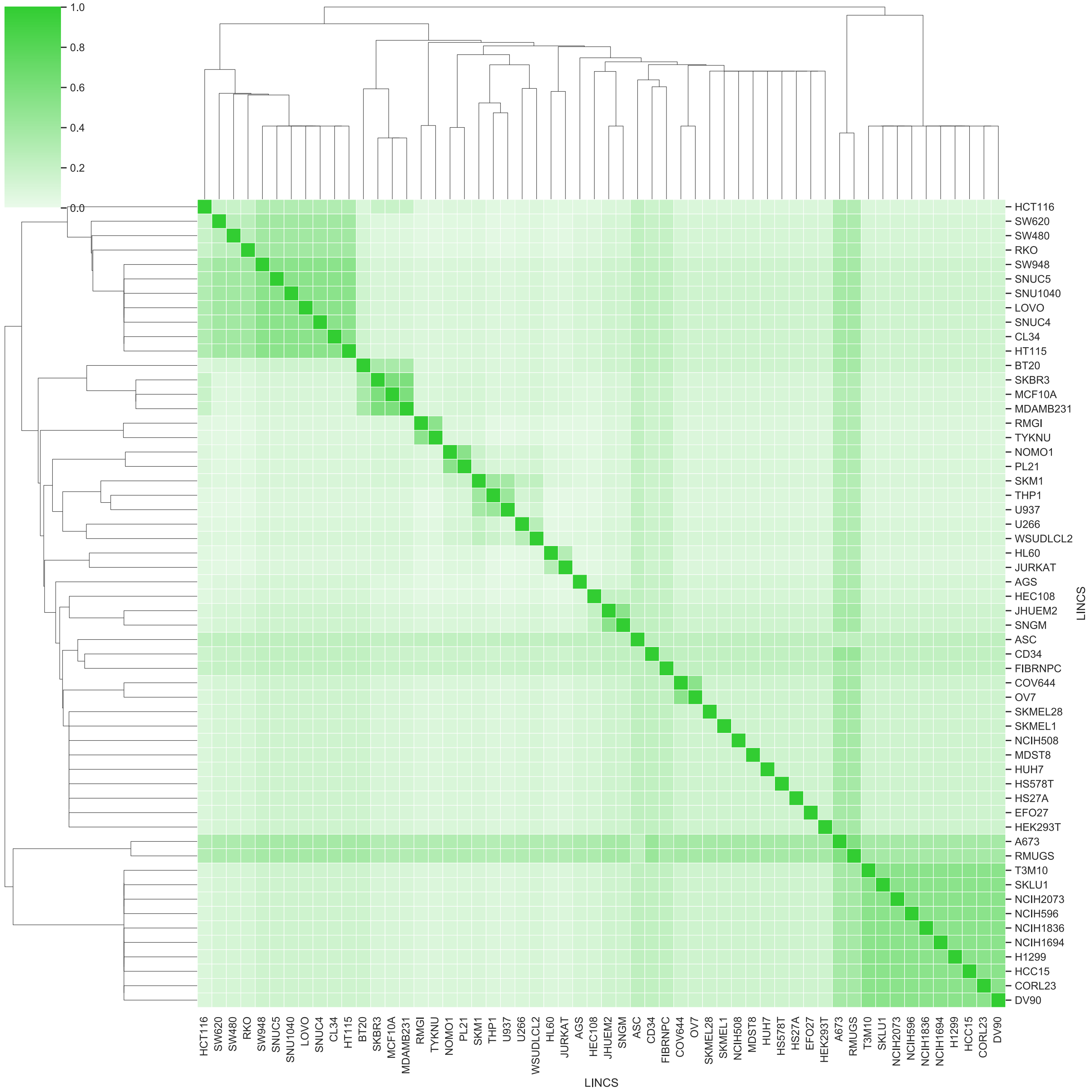


**B**

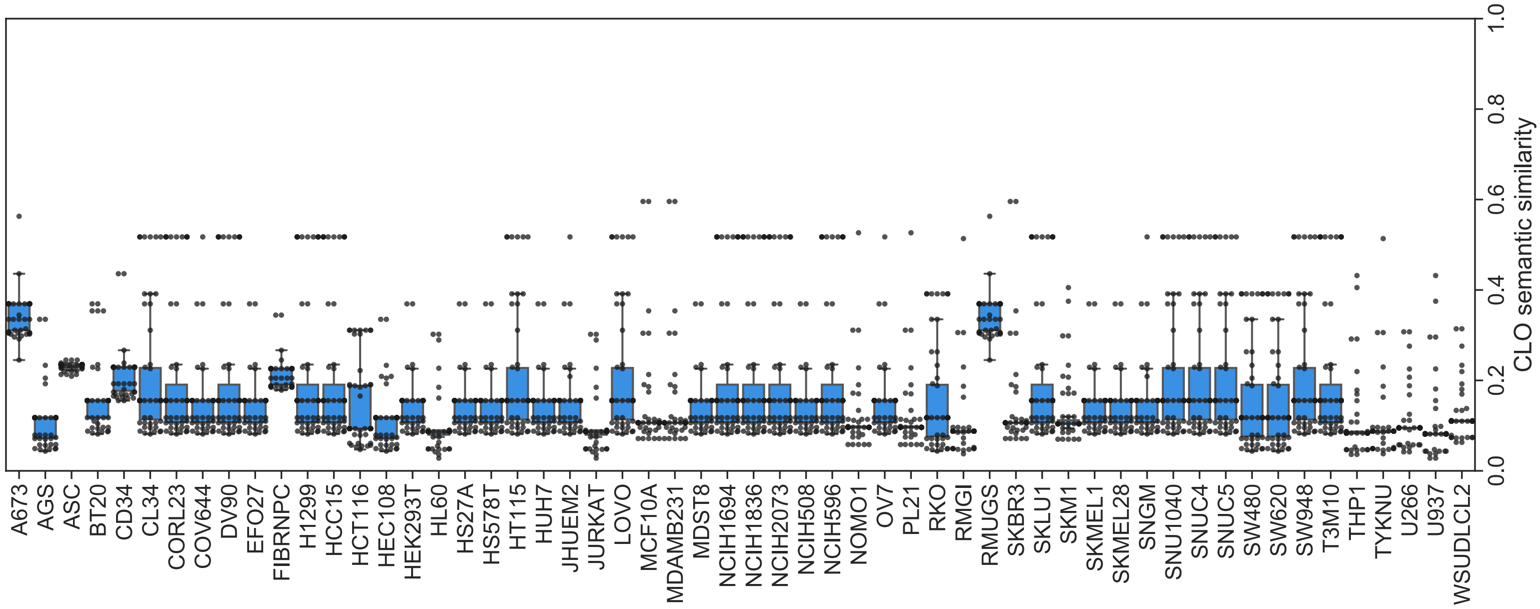

### Supplementary Figure 8

**A**

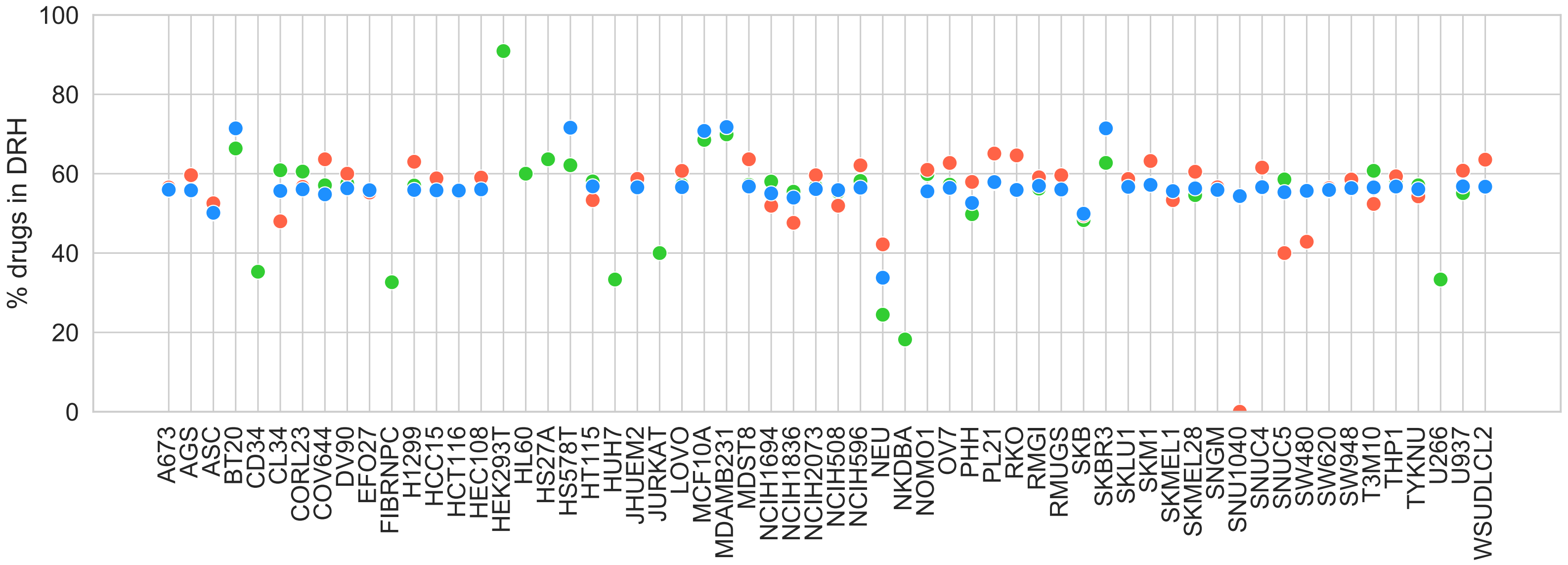

### Supplementary Figure 9

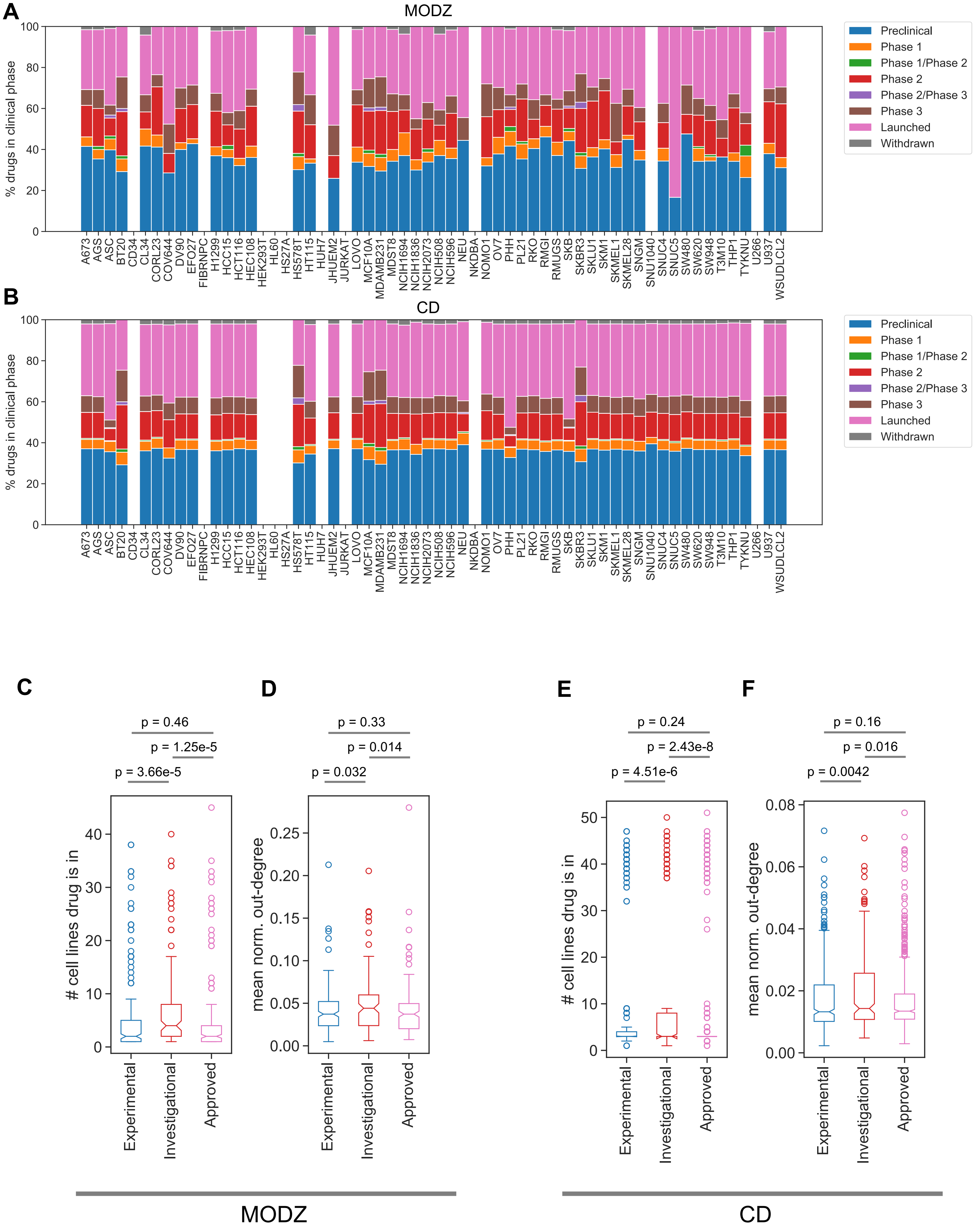

### Supplementary Figure 10

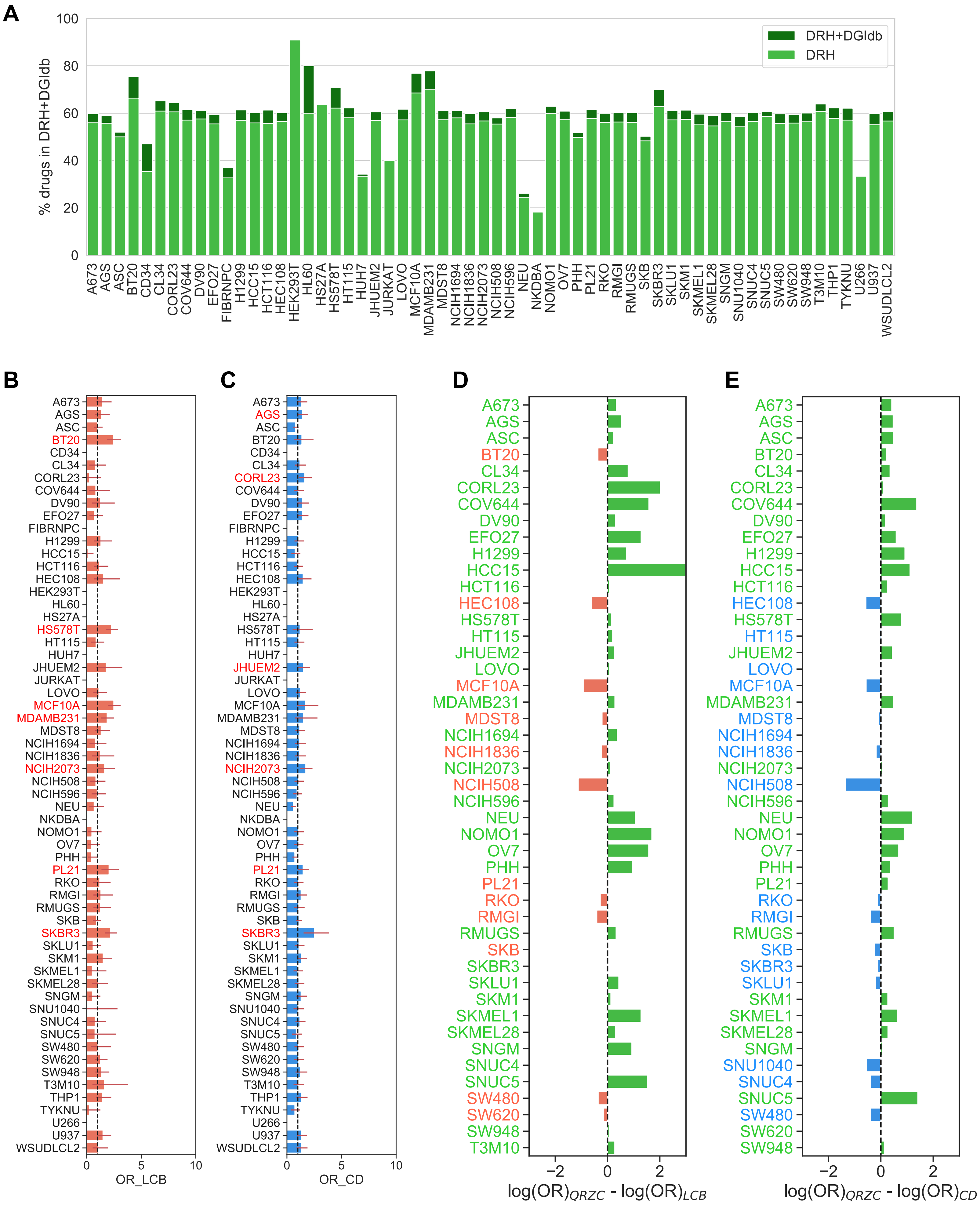

### Supplementary Figure 11

**A**

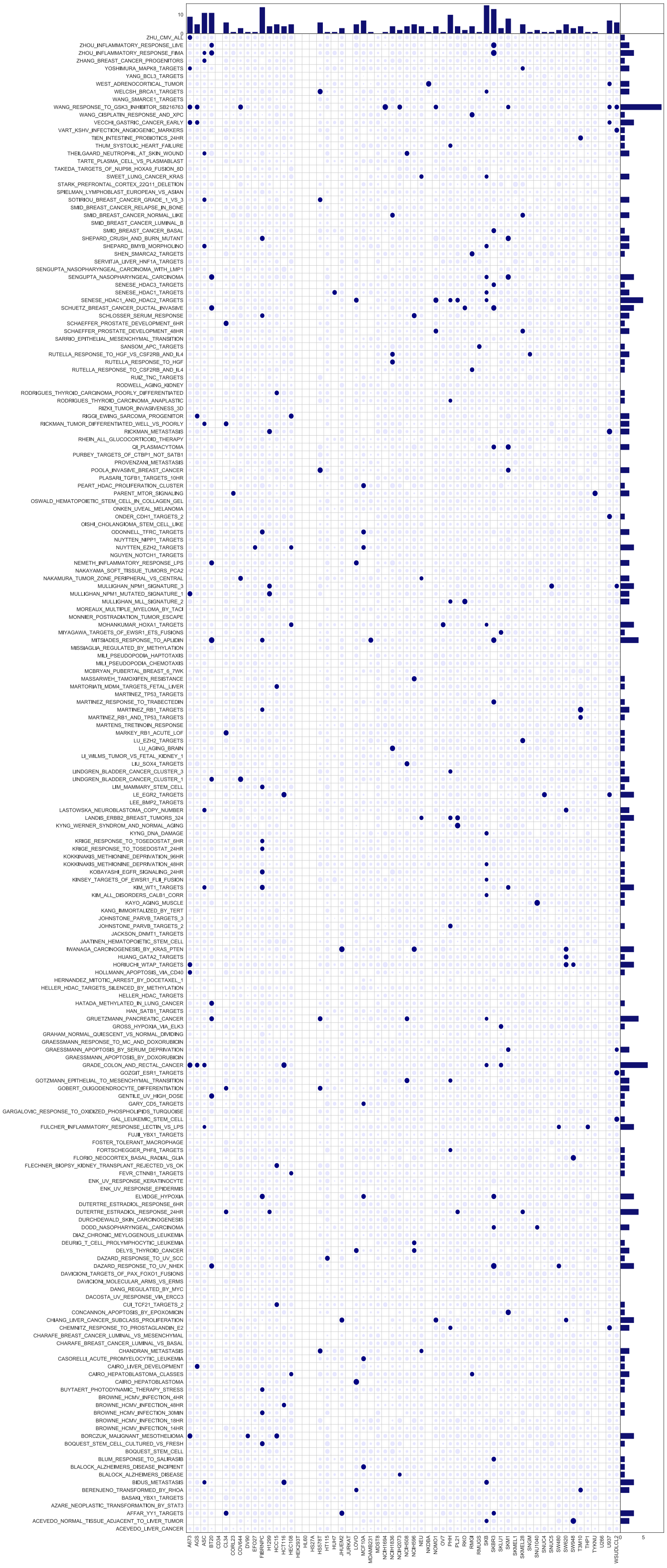

**B**

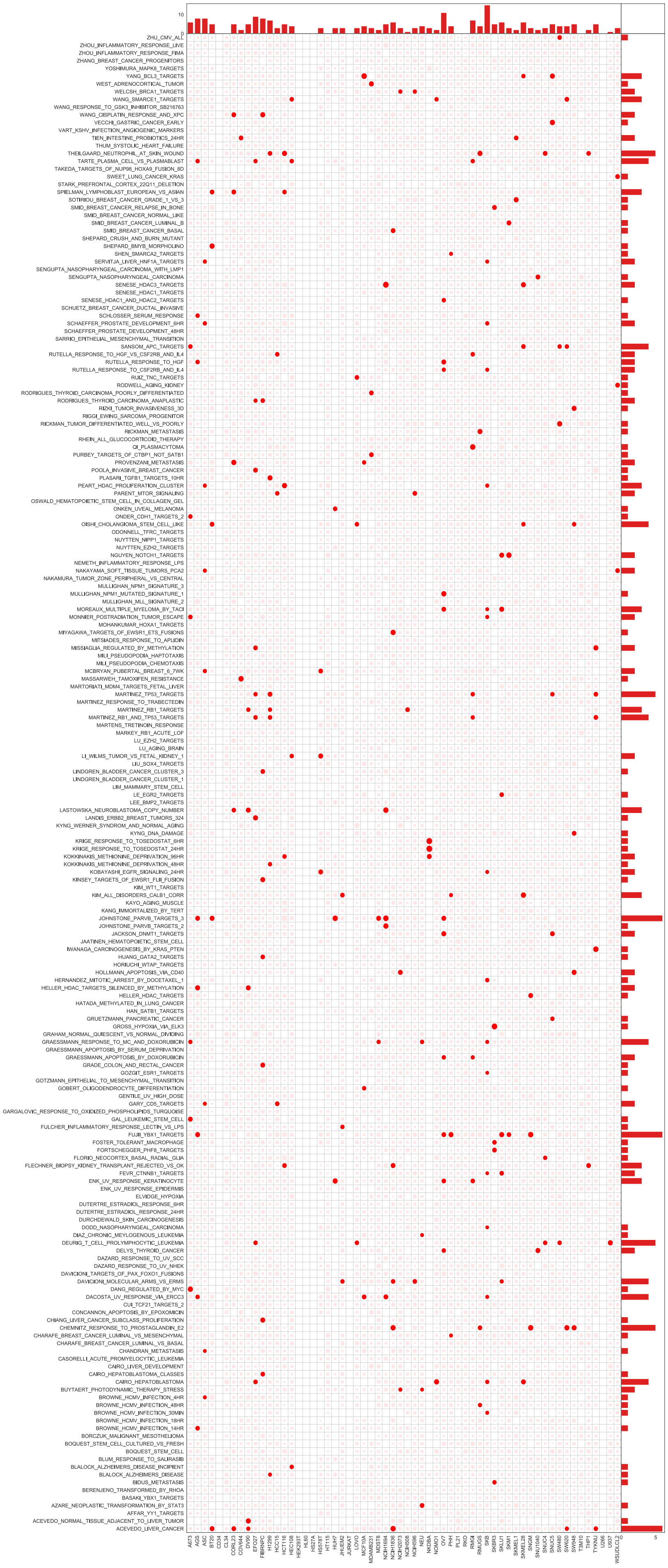

### Supplementary Figure 12

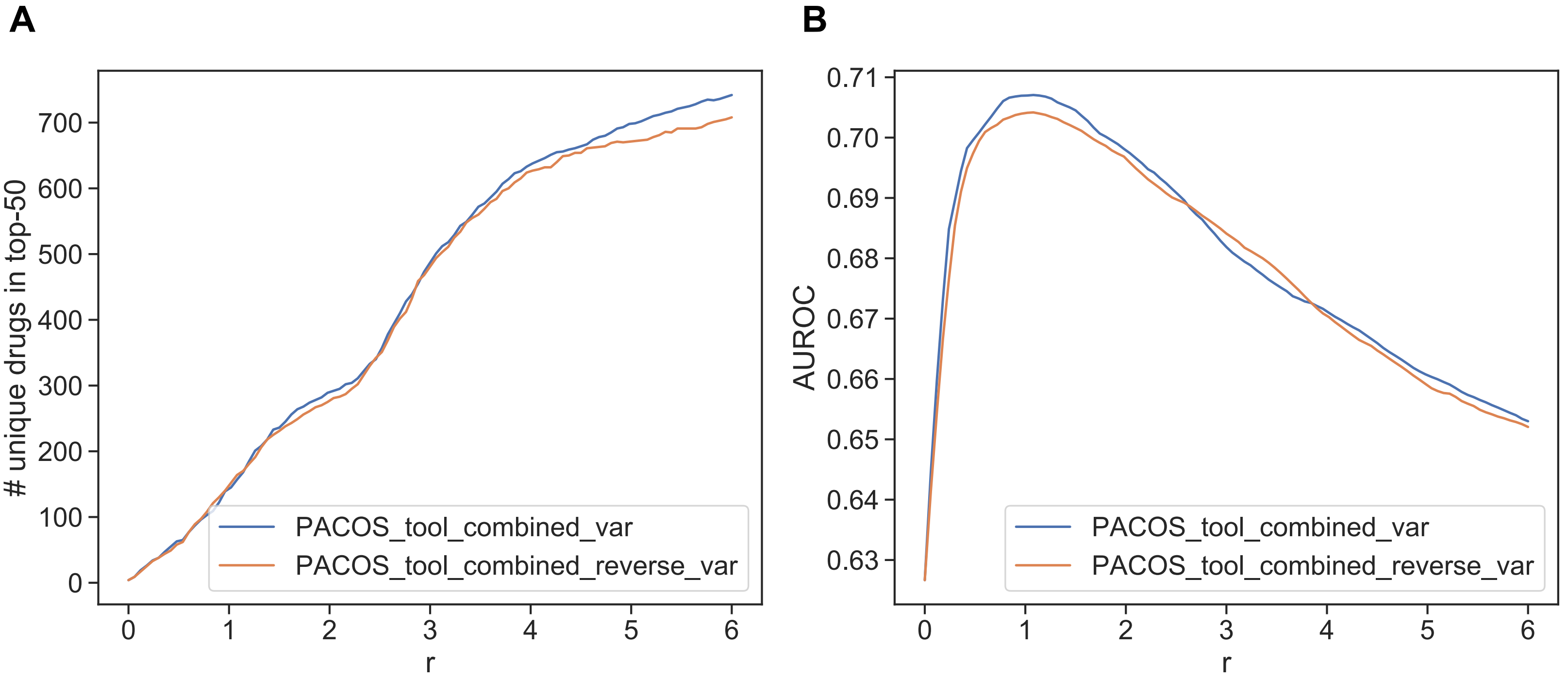

### Supplementary Figure 13

## A

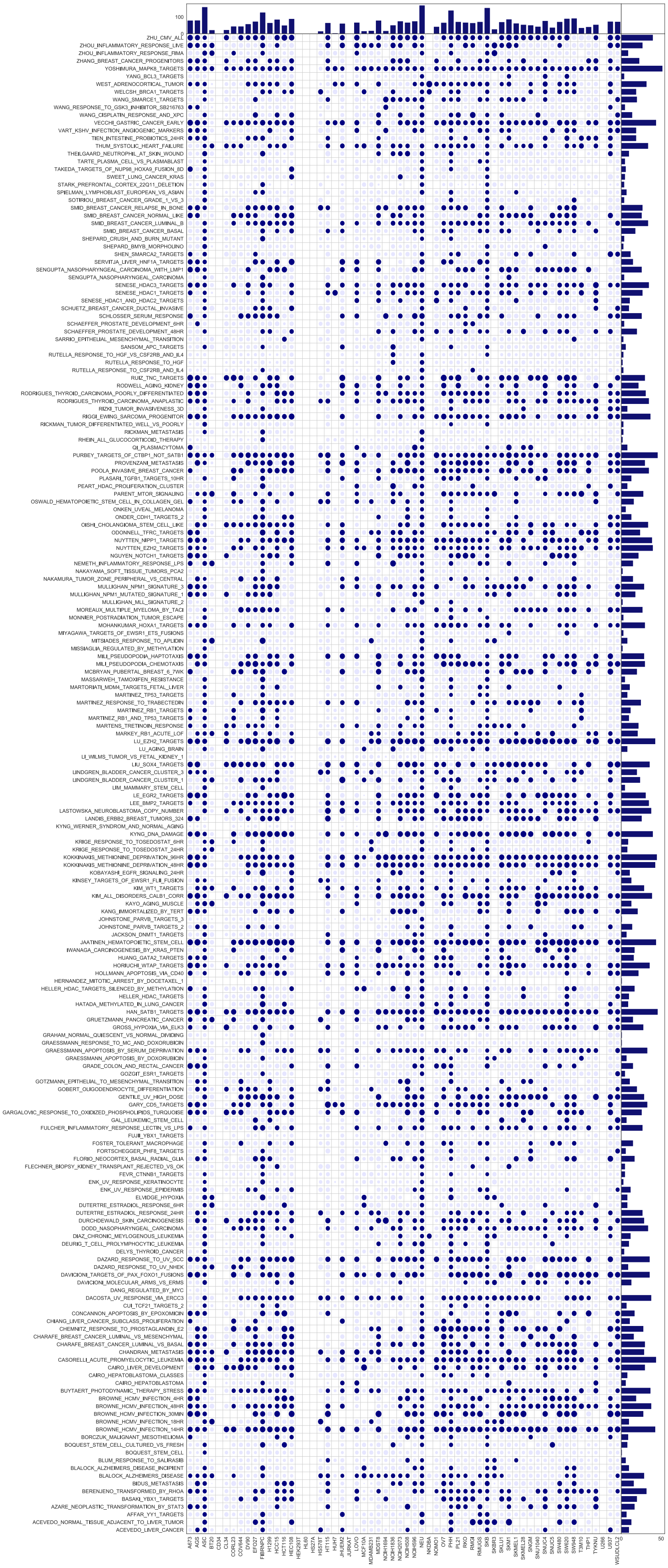

## B

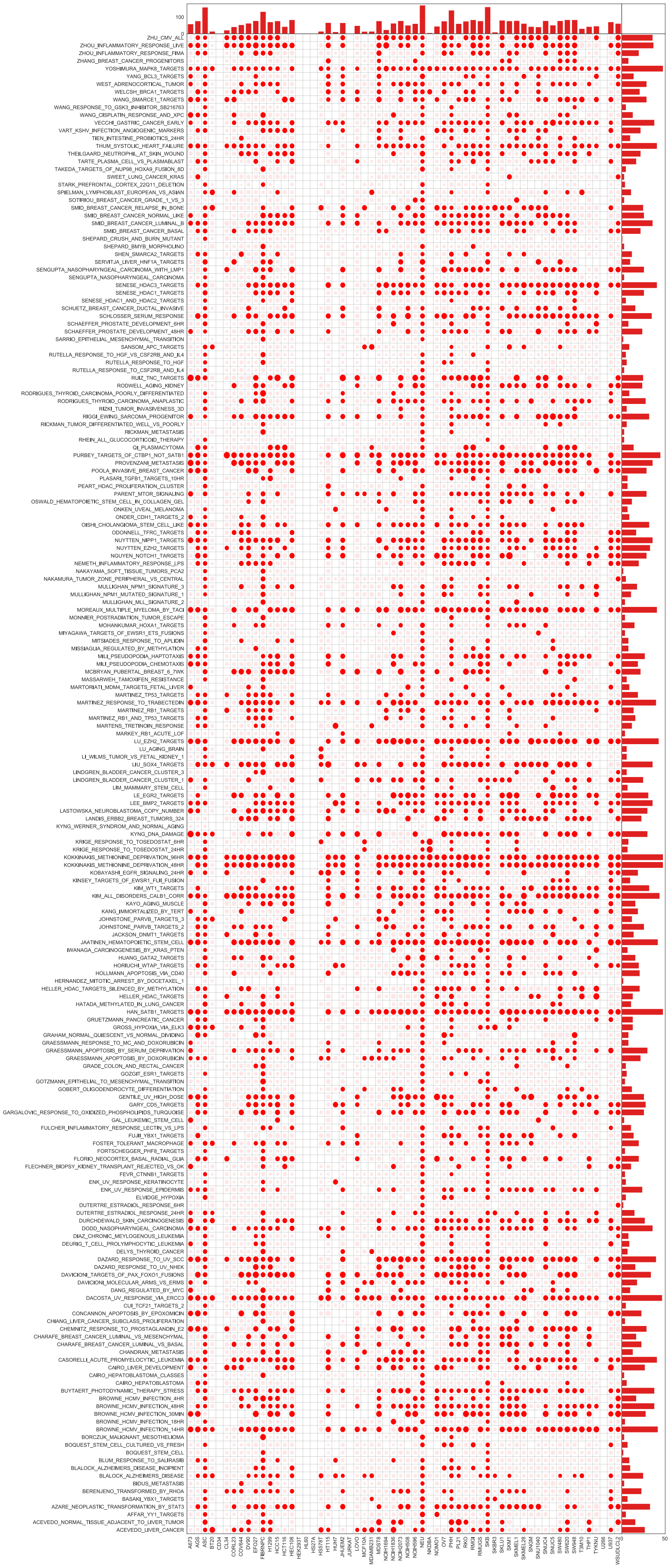

### Supplementary Figure 14

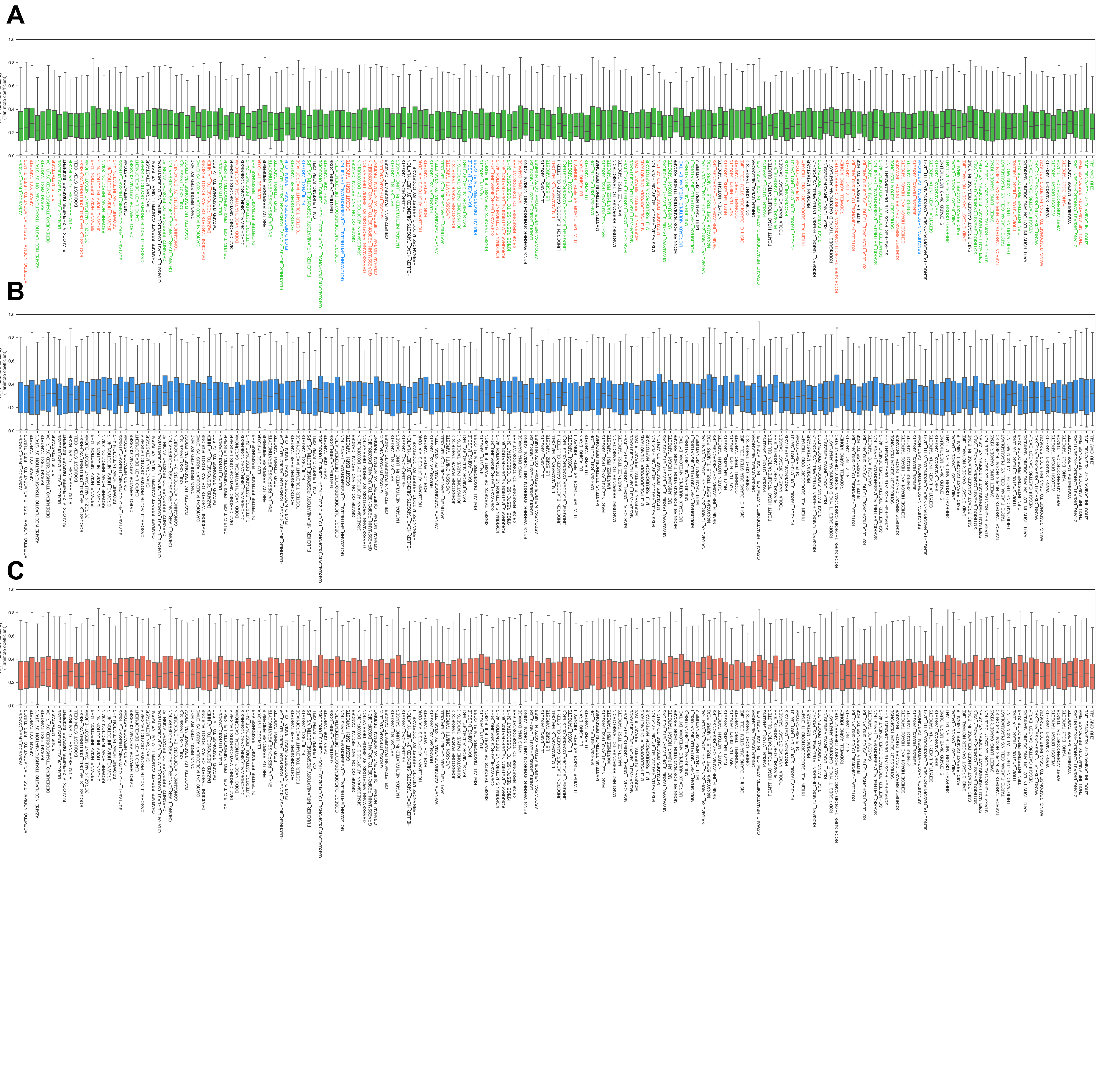

### Supplementary Figure 15

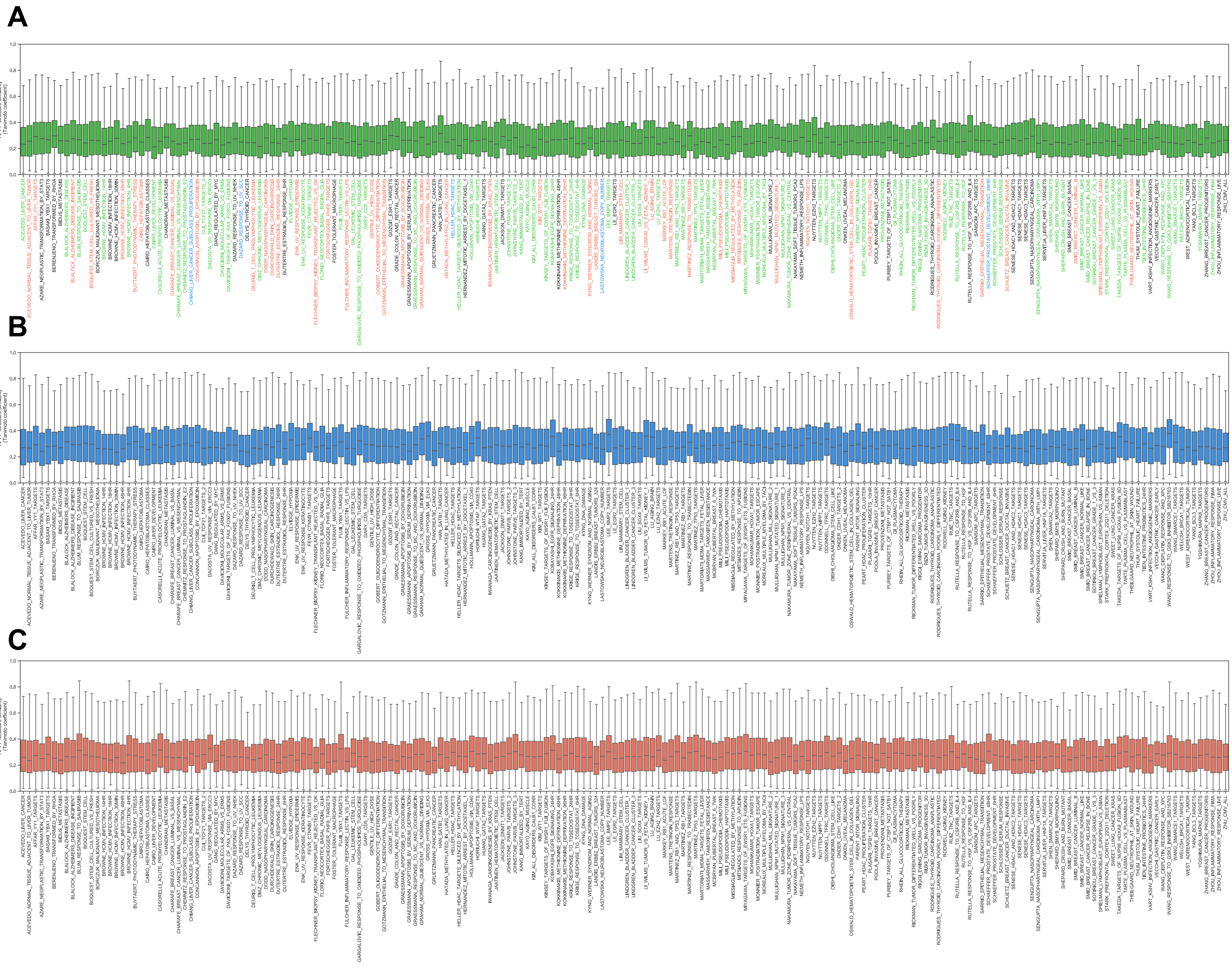

### Supplementary Figure 16

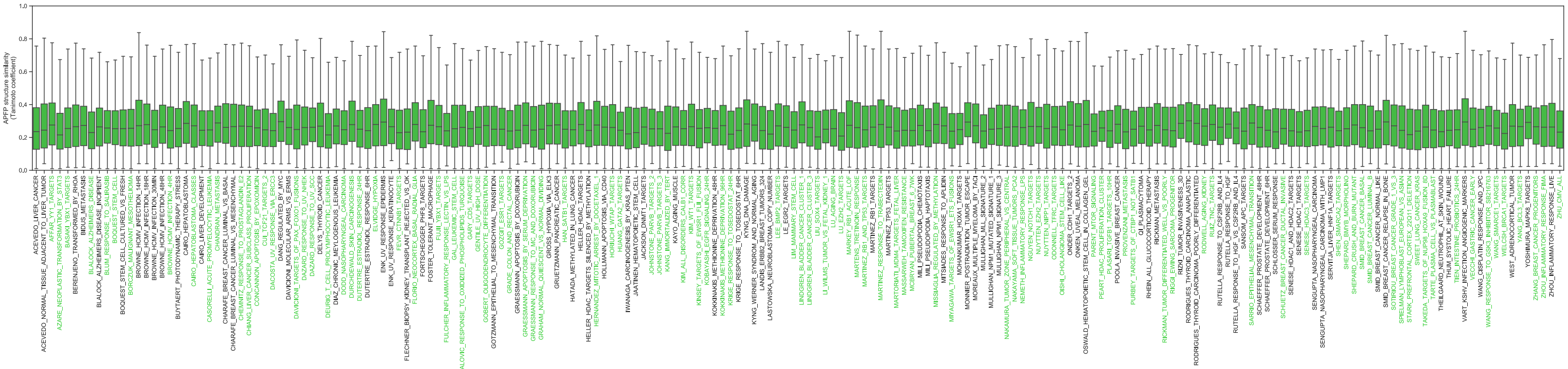


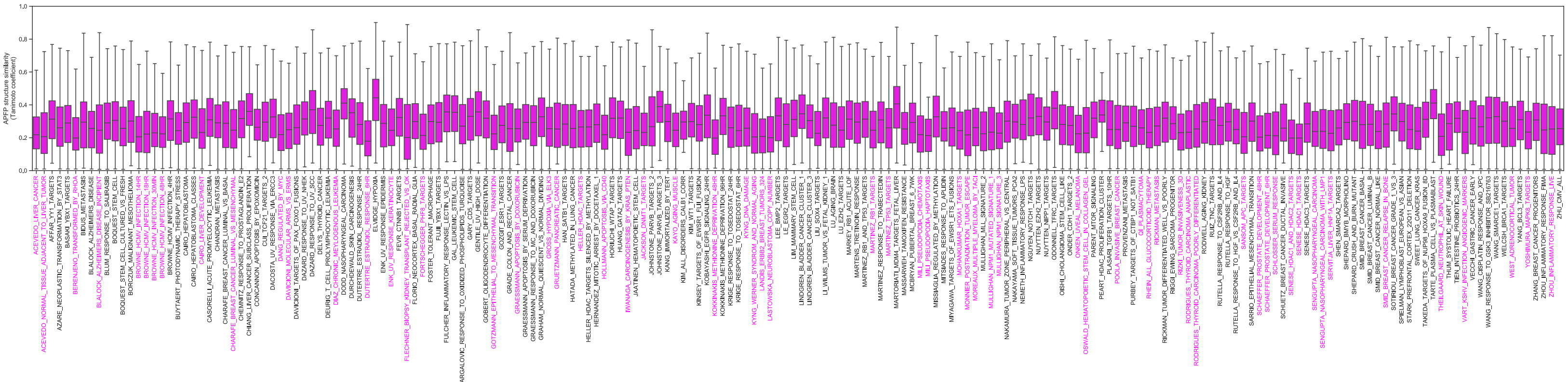

### Supplementary Figure 17

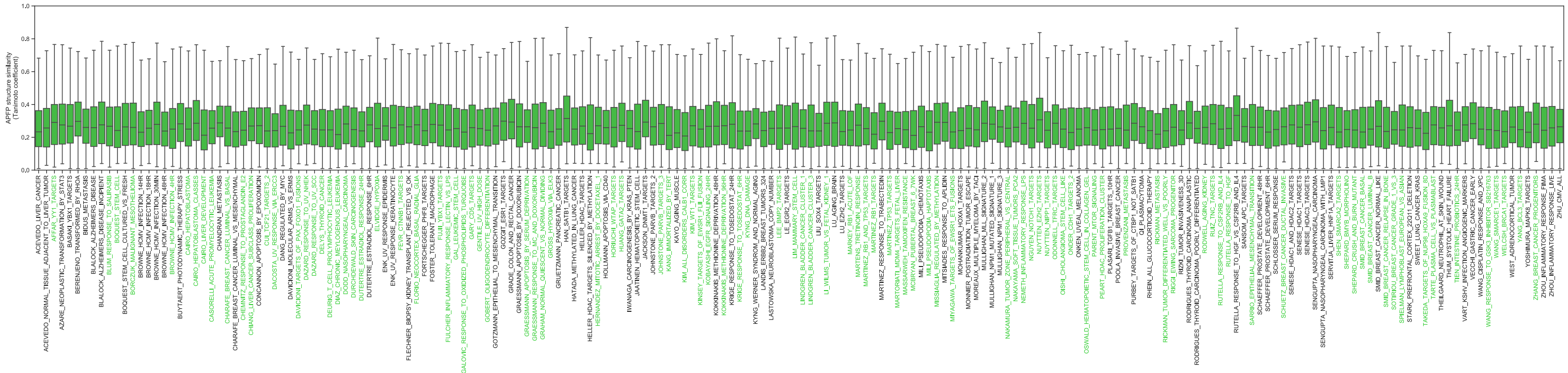


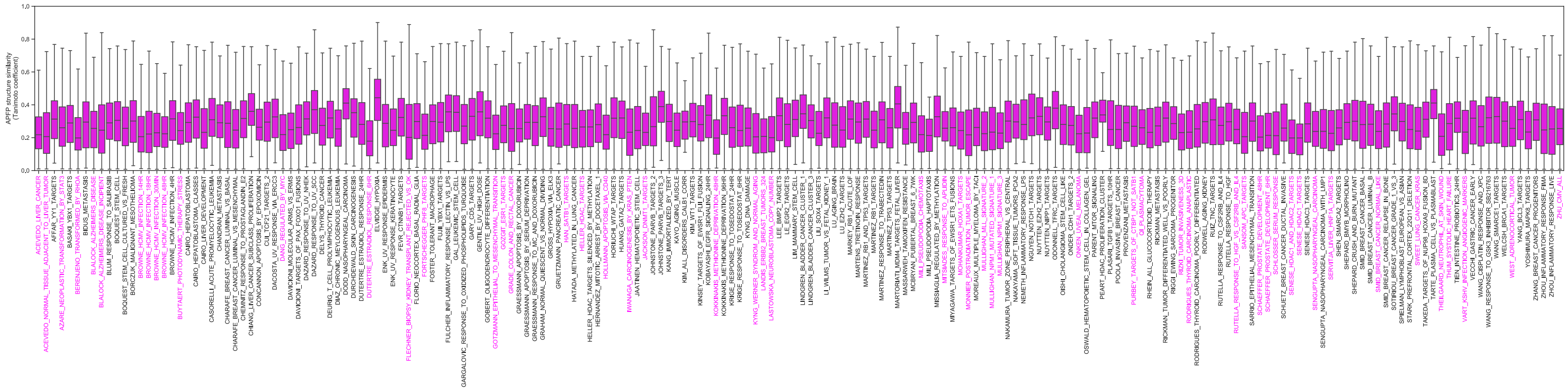

### Supplementary Figure 18

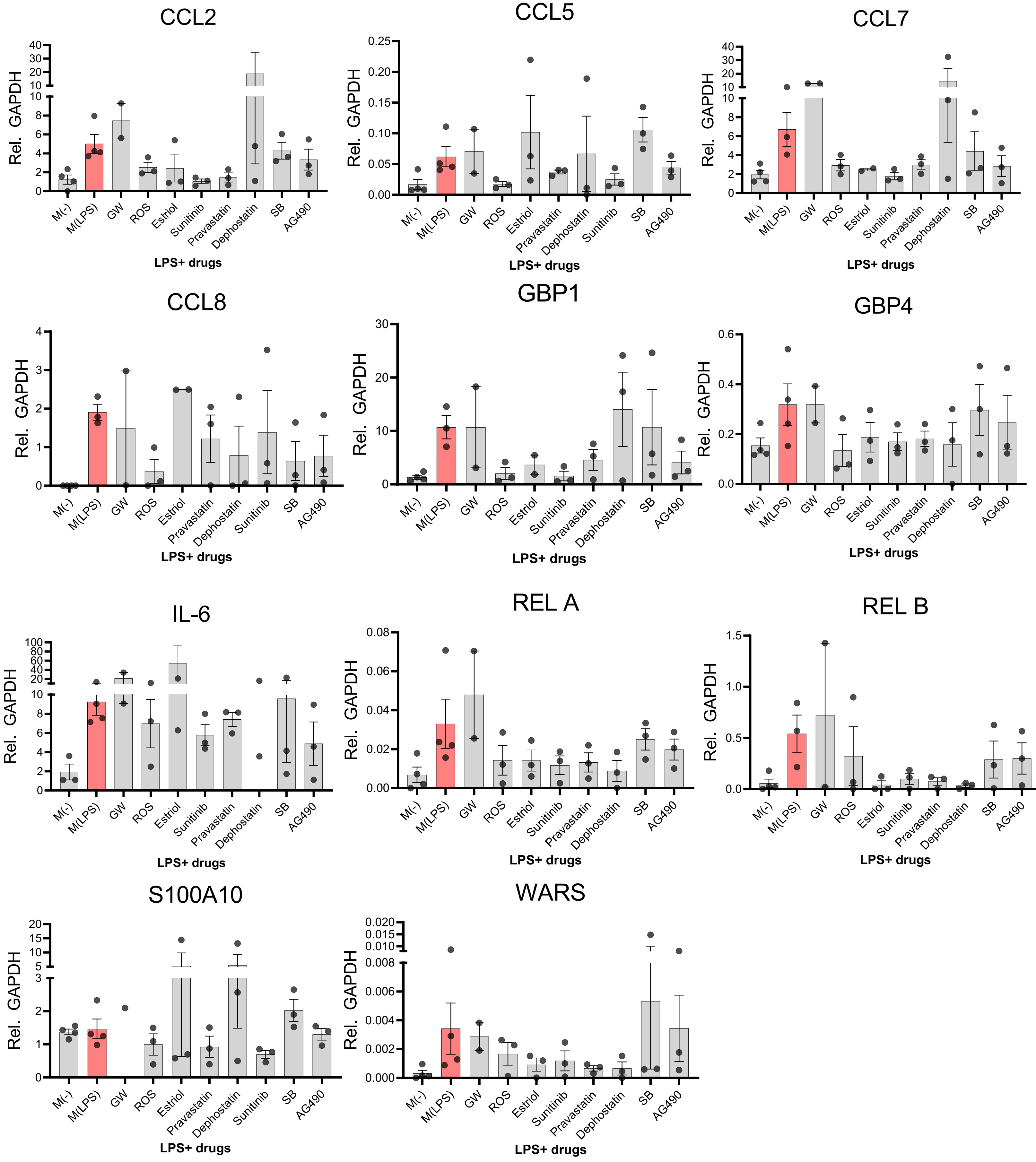

### Supplementary Figure 19

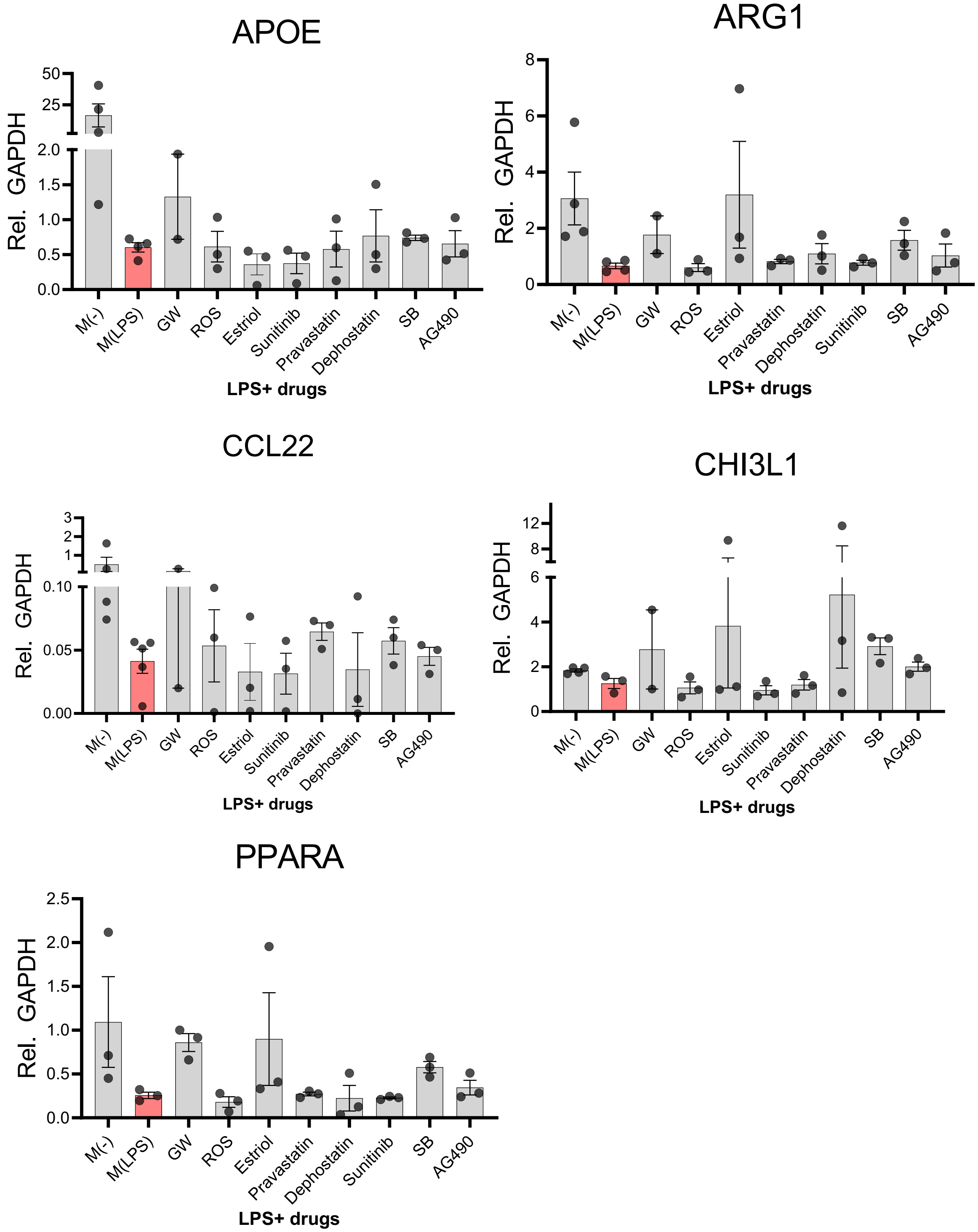

### Supplementary Figure 20

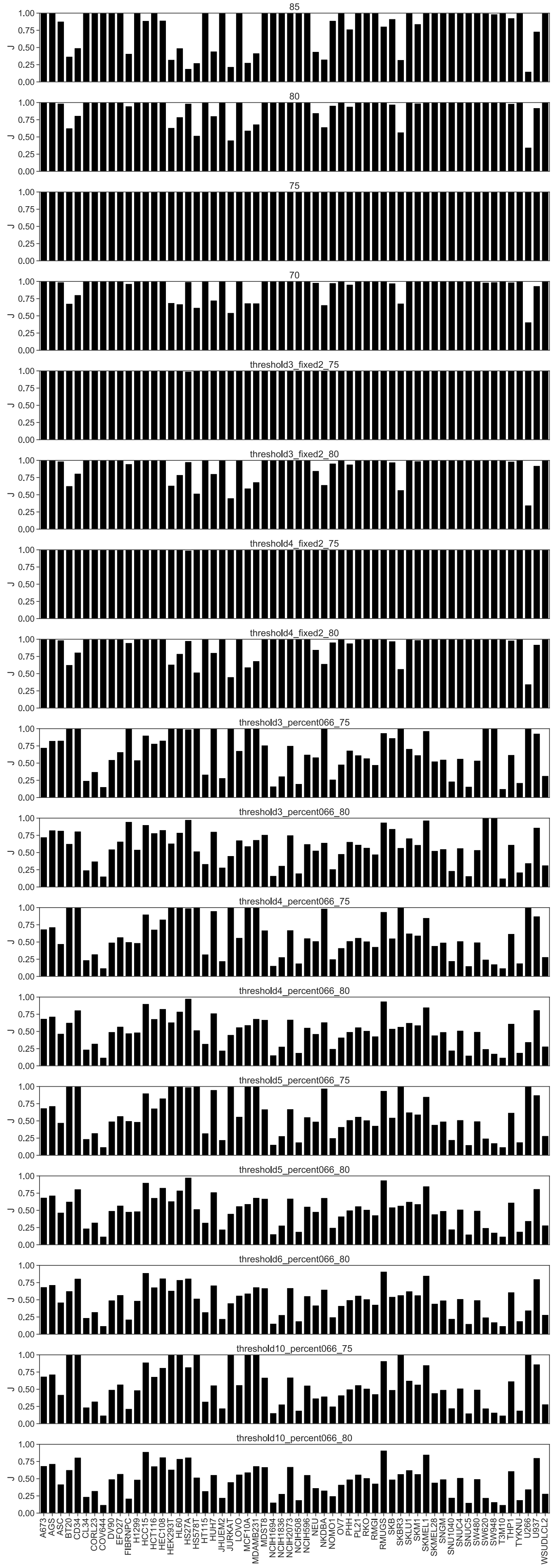

### Supplementary Table 1

| Cell_type | N_drugs | N_targets | M | Density |
|---|---|---|---|---|
| A673 | 311 | 10065 | 26669 | 0.00851986 |
| AGS | 330 | 9572 | 21579 | 0.00683148 |
| ASC | 2298 | 12323 | 340291 | 0.01201667 |
| BT20 | 110 | 8469 | 15978 | 0.01715132 |
| CD34 | 17 | 7567 | 9257 | 0.07196107 |
| CL34 | 184 | 9202 | 17116 | 0.01010886 |
| CORL23 | 261 | 5867 | 8460 | 0.00552476 |
| COV644 | 226 | 5889 | 7886 | 0.00592525 |
| DV90 | 332 | 7739 | 12364 | 0.00481212 |
| EFO27 | 330 | 9295 | 18979 | 0.00618743 |
| FIBRNPC | 539 | 12036 | 56767 | 0.00875034 |
| H1299 | 321 | 7867 | 13685 | 0.00541914 |
| HCC15 | 324 | 10132 | 23718 | 0.007225 |
| HCT116 | 212 | 5399 | 9550 | 0.00834361 |
| HEC108 | 349 | 9952 | 22329 | 0.00642885 |
| HEK293T | 11 | 1761 | 2037 | 0.10515719 |
| HL60 | 10 | 6431 | 9238 | 0.14364796 |
| HS27A | 11 | 572 | 574 | 0.09122695 |
| HS578T | 103 | 5317 | 9801 | 0.01789643 |
| HT115 | 217 | 10036 | 20912 | 0.0096023 |
| HUH7 | 111 | 10790 | 31166 | 0.02602176 |
| JHUEM2 | 304 | 7596 | 12139 | 0.00525684 |
| JURKAT | 5 | 2261 | 2448 | 0.21654135 |
| LOVO | 308 | 9667 | 22390 | 0.00751989 |
| MCF10A | 108 | 6518 | 11377 | 0.0161618 |
| MDAMB231 | 113 | 9004 | 25005 | 0.0245761 |
| MDST8 | 327 | 9804 | 22964 | 0.00716303 |
| NCIH1694 | 162 | 8037 | 14018 | 0.01076656 |
| NCIH1836 | 229 | 8304 | 13939 | 0.00733008 |
| NCIH2073 | 335 | 10162 | 22218 | 0.00652651 |
| NCIH508 | 231 | 5025 | 7123 | 0.00613642 |
| NCIH596 | 318 | 8082 | 15339 | 0.00596831 |
| NEU | 2183 | 12237 | 134986 | 0.00505312 |
| NKDBA | 192 | 10289 | 26398 | 0.01336277 |
| NOMO1 | 202 | 10587 | 24002 | 0.01122337 |
| OV7 | 332 | 7090 | 12122 | 0.0051498 |
| PHH | 1695 | 12295 | 139171 | 0.00667806 |
| PL21 | 312 | 10509 | 28157 | 0.00858757 |
| RKO | 332 | 8689 | 15565 | 0.00539562 |
| RMGI | 320 | 6906 | 11108 | 0.00502643 |
| RMUGS | 344 | 9585 | 22776 | 0.0069076 |
| SKB | 2178 | 12310 | 292285 | 0.01090161 |
| SKBR3 | 110 | 7747 | 16984 | 0.0199303 |
| SKLU1 | 329 | 8361 | 15322 | 0.00557008 |
| SKM1 | 310 | 10510 | 26402 | 0.0081035 |
| SKMEL1 | 334 | 6666 | 10385 | 0.00466439 |
| SKMEL28 | 315 | 9193 | 18395 | 0.00635231 |
| SNGM | 266 | 6480 | 11687 | 0.00678026 |
| SNU1040 | 225 | 4631 | 6372 | 0.00611531 |
| SNUC4 | 338 | 7429 | 12718 | 0.00506491 |
| SNUC5 | 181 | 8798 | 16096 | 0.01010777 |
| SW480 | 345 | 6516 | 9644 | 0.00429 |
| SW620 | 353 | 12126 | 54640 | 0.01276493 |
| SW948 | 353 | 12111 | 58536 | 0.01369205 |
| T3M10 | 191 | 8111 | 12267 | 0.00791828 |
| THP1 | 294 | 12029 | 56016 | 0.01583927 |
| TYKNU | 219 | 3766 | 4998 | 0.00605999 |
| U266 | 6 | 1305 | 1448 | 0.18492976 |
| U937 | 294 | 9821 | 26254 | 0.00909269 |
| WSUDLCL2 | 321 | 8645 | 16694 | 0.00601576 |

### Supplementary Table 2

| Geneset_name (page 1/3) |
|---|
| ACEVEDO_LIVER_CANCER |
| ACEVEDO_NORMAL_TISSUE_ADJACENT_TO_LIVER_TUMOR |
| AFFAR_YY1_TARGETS |
| AZARE_NEOPLASTIC_TRANSFORMATION_BY_STAT3 |
| BASAKI_YBX1_TARGETS |
| BERENJENO_TRANSFORMED_BY_RHOA |
| BIDUS_METASTASIS |
| BLALOCK_ALZHEIMERS_DISEASE |
| BLALOCK_ALZHEIMERS_DISEASE_INCIPIENT |
| BLUM_RESPONSE_TO_SALIRASIB |
| BOQUEST_STEM_CELL |
| BOQUEST_STEM_CELL_CULTURED_VS_FRESH |
| BORCZUK_MALIGNANT_MESOTHELIOMA |
| BROWNE_HCMV_INFECTION_14HR |
| BROWNE_HCMV_INFECTION_18HR |
| BROWNE_HCMV_INFECTION_30MIN |
| BROWNE_HCMV_INFECTION_48HR |
| BROWNE_HCMV_INFECTION_4HR |
| BUYTAERT_PHOTODYNAMIC_THERAPY_STRESS |
| CAIRO_HEPATOBLASTOMA |
| CAIRO_HEPATOBLASTOMA_CLASSES |
| CAIRO_LIVER_DEVELOPMENT |
| CASORELLI_ACUTE_PROMYELOCYTIC_LEUKEMIA |
| CHANDRAN_METASTASIS |
| CHARAFE_BREAST_CANCER_LUMINAL_VS_BASAL |
| CHARAFE_BREAST_CANCER_LUMINAL_VS_MESENCHYMAL |
| CHEMNITZ_RESPONSE_TO_PROSTAGLANDIN_E2 |
| CHIANG_LIVER_CANCER_SUBCLASS_PROLIFERATION |
| CONCANNON_APOPTOSIS_BY_EPOXOMICIN |
| CUI_TCF21_TARGETS_2 |
| DACOSTA_UV_RESPONSE_VIA_ERCC3 |
| DANG_REGULATED_BY_MYC |
| DAVICIONI_MOLECULAR_ARMS_VS_ERMS |
| DAVICIONI_TARGETS_OF_PAX_FOXO1_FUSIONS |
| DAZARD_RESPONSE_TO_UV_NHEK |
| DAZARD_RESPONSE_TO_UV_SCC |
| DELYS_THYROID_CANCER |
| DEURIG_T_CELL_PROLYMPHOCYTIC_LEUKEMIA |
| DURCHDEWALD_SKIN_CARCINOGENESIS |
| DUTERTRE_ESTRADIOL_RESPONSE_24HR |
| DUTERTRE_ESTRADIOL_RESPONSE_6HR |
| ELVIDGE_HYPOXIA |
| ENK_UV_RESPONSE_EPIDERMIS |
| ENK_UV_RESPONSE_KERATINOCYTE |
| FEVR_CTNNB1_TARGETS |
| FLECHNER_BIOPSY_KIDNEY_TRANSPLANT_REJECTED_VS_OK |
| FLORIO_NEOCORTEX_BASAL_RADIAL_GLIA |
| FORTSCHEGGER_PHF8_TARGETS |
| FOSTER_TOLERANT_MACROPHAGE |
| FUJII_YBX1_TARGETS |
| FULCHER_INFLAMMATORY_RESPONSE_LECTIN_VS_LPS |
| GAL_LEUKEMIC_STEM_CELL |
| GARGALOVIC_RESPONSE_TO_OXIDIZED_PHOSPHOLIPIDS_TURQUOISE |
| GARY_CD5_TARGETS |
| GENTILE_UV_HIGH_DOSE |
| GOBERT_OLIGODENDROCYTE_DIFFERENTIATION |
| GOTZMANN_EPITHELIAL_TO_MESENCHYMAL_TRANSITION |
| GOZGIT_ESR1_TARGETS |
| GRADE_COLON_AND_RECTAL_CANCER |
| GRAESSMANN_APOPTOSIS_BY_DOXORUBICIN |
| GRAESSMANN_APOPTOSIS_BY_SERUM_DEPRIVATION |
| GRAESSMANN_RESPONSE_TO_MC_AND_DOXORUBICIN |

### Supplementary Table 2 (cont'd)

| Geneset_name (page 2/3) |
|---|
| GRAHAM_NORMAL_QUIESCENT_VS_NORMAL_DIVIDING |
| GROSS_HYPOXIA_VIA_ELK3 |
| GRUETZMANN_PANCREATIC_CANCER |
| HAN_SATB1_TARGETS |
| HATADA_METHYLATED_IN_LUNG_CANCER |
| HELLER_HDAC_TARGETS |
| HELLER_HDAC_TARGETS_SILENCED_BY_METHYLATION |
| HERNANDEZ_MITOTIC_ARREST_BY_DOCETAXEL_1 |
| HOLLMANN_APOPTOSIS_VIA_CD40 |
| HORIUCHI_WTAP_TARGETS |
| HUANG_GATA2_TARGETS |
| IWANAGA_CARCINOGENESIS_BY_KRAS_PTEN |
| JAATINEN_HEMATOPOIETIC_STEM_CELL |
| JACKSON_DNMT1_TARGETS |
| JOHNSTONE_PARVB_TARGETS_2 |
| JOHNSTONE_PARVB_TARGETS_3 |
| KANG_IMMORTALIZED_BY_TERT |
| KAYO_AGING_MUSCLE |
| KIM_ALL_DISORDERS_CALB1_CORR |
| KIM_WT1_TARGETS |
| KINSEY_TARGETS_OF_EWSR1_FLII_FUSION |
| KOBAYASHI_EGFR_SIGNALING_24HR |
| KOKKINAKIS_METHIONINE_DEPRIVATION_48HR |
| KOKKINAKIS_METHIONINE_DEPRIVATION_96HR |
| KRIGE_RESPONSE_TO_TOSEDOSTAT_24HR |
| KRIGE_RESPONSE_TO_TOSEDOSTAT_6HR |
| KYNG_DNA_DAMAGE |
| KYNG_WERNER_SYNDROM_AND_NORMAL_AGING |
| LANDIS_ERBB2_BREAST_TUMORS_324 |
| LASTOWSKA_NEUROBLASTOMA_COPY_NUMBER |
| LEE_BMP2_TARGETS |
| LE_EGR2_TARGETS |
| LIM_MAMMARY_STEM_CELL |
| LINDGREN_BLADDER_CANCER_CLUSTER_1 |
| LINDGREN_BLADDER_CANCER_CLUSTER_3 |
| LIU_SOX4_TARGETS |
| LI_WILMS_TUMOR_VS_FETAL_KIDNEY_1 |
| LU_AGING_BRAIN |
| LU_EZH2_TARGETS |
| MARKEY_RB1_ACUTE_LOF |
| MARTENS_TRETINOIN_RESPONSE |
| MARTINEZ_RB1_AND_TP53_TARGETS |
| MARTINEZ_RB1_TARGETS |
| MARTINEZ_RESPONSE_TO_TRABECTEDIN |
| MARTINEZ_TP53_TARGETS |
| MARTORIATI_MDM4_TARGETS_FETAL_LIVER |
| MASSARWEH_TAMOXIFEN_RESISTANCE |
| MCBRYAN_PUBERTAL_BREAST_6_7WK |
| MILI_PSEUDOPODIA_CHEMOTAXIS |
| MILI_PSEUDOPODIA_HAPTOTAXIS |
| MISSIAGLIA_REGULATED_BY_METHYLATION |
| MITSIADES_RESPONSE_TO_APLIDIN |
| MIYAGAWA_TARGETS_OF_EWSR1_ETS_FUSIONS |
| MOHANKUMAR_HOXA1_TARGETS |
| MONNIER_POSTRADIATION_TUMOR_ESCAPE |
| MOREAUX_MULTIPLE_MYELOMA_BY_TACI |
| MULLIGHAN_MLL_SIGNATURE_2 |
| MULLIGHAN_NPM1_MUTATED_SIGNATURE_1 |
| MULLIGHAN_NPM1_SIGNATURE_3 |
| NAKAMURA_TUMOR_ZONE_PERIPHERAL_VS_CENTRAL |
| NAKAYAMA_SOFT_TISSUE_TUMORS_PCA2 |
| NEMETH_INFLAMMATORY_RESPONSE_LPS |
| NGUYEN_NOTCH1_TARGETS |
| NUYTTEN_EZH2_TARGETS |
| NUYTTEN_NIPP1_TARGETS |

### Supplementary Table 2 (cont'd)

| Geneset_name (page 3/3) |
|---|
| ODONNELL_TFRC_TARGETS |
| OISHI_CHOLANGIOMA_STEM_CELL_LIKE |
| ONDER_CDH1_TARGETS_2 |
| ONKEN_UVEAL_MELANOMA |
| OSWALD_HEMATOPOIETIC_STEM_CELL_IN_COLLAGEN_GEL |
| PARENT_MTOR_SIGNALING |
| PEART_HDAC_PROLIFERATION_CLUSTER |
| PLASARI_TGFB1_TARGETS_10HR |
| POOLA_INVASIVE_BREAST_CANCER |
| PROVENZANI_METASTASIS |
| PURBEY_TARGETS_OF_CTBP1_NOT_SATB1 |
| QI_PLASMACYTOMA |
| RHEIN_ALL_GLUCOCORTICOID_THERAPY |
| RICKMAN_METASTASIS |
| RICKMAN_TUMOR_DIFFERENTIATED_WELL_VS_POORLY |
| RIGGI_EWING_SARCOMA_PROGENITOR |
| RIZKI_TUMOR_INVASIVENESS_3D |
| RODRIGUES_THYROID_CARCINOMA_ANAPLASTIC |
| RODRIGUES_THYROID_CARCINOMA_POORLY_DIFFERENTIATED |
| RODWELL_AGING_KIDNEY |
| RUIZ_TNC_TARGETS |
| RUTELLA_RESPONSE_TO_CSF2RB_AND_IL4 |
| RUTELLA_RESPONSE_TO_HGF |
| RUTELLA_RESPONSE_TO_HGF_VS_CSF2RB_AND_IL4 |
| SANSOM_APC_TARGETS |
| SARRIO_EPITHELIAL_MESENCHYMAL_TRANSITION |
| SCHAEFFER_PROSTATE_DEVELOPMENT_48HR |
| SCHAEFFER_PROSTATE_DEVELOPMENT_6HR |
| SCHLOSSER_SERUM_RESPONSE |
| SCHUETZ_BREAST_CANCER_DUCTAL_INVASIVE |
| SENESE_HDAC1_AND_HDAC2_TARGETS |
| SENESE_HDAC1_TARGETS |
| SENESE_HDAC3_TARGETS |
| SENGUPTA_NASOPHARYNGEAL_CARCINOMA |
| SENGUPTA_NASOPHARYNGEAL_CARCINOMA_WITH_LMP1 |
| SERVITJA_LIVER_HNF1A_TARGETS |
| SHEN_SMARCA2_TARGETS |
| SHEPARD_BMYB_MORPHOLINO |
| SHEPARD_CRUSH_AND_BURN_MUTANT |
| SMID_BREAST_CANCER_BASAL |
| SMID_BREAST_CANCER_LUMINAL_B |
| SMID_BREAST_CANCER_NORMAL_LIKE |
| SMID_BREAST_CANCER_RELAPSE_IN_BONE |
| SOTIRIOU_BREAST_CANCER_GRADE_1_VS_3 |
| SPIELMAN_LYMPHOBLAST_EUROPEAN_VS_ASIAN |
| STARK_PREFRONTAL_CORTEX_22Q11_DELETION |
| SWEET_LUNG_CANCER_KRAS |
| TAKEDA_TARGETS_OF_NUP98_HOXA9_FUSION_8D |
| TARTE_PLASMA_CELL_VS_PLASMABLAST |
| THEILGAARD_NEUTROPHIL_AT_SKIN_WOUND |
| THUM_SYSTOLIC_HEART_FAILURE |
| TIEN_INTESTINE_PROBIOTICS_24HR |
| VART_KSHV_INFECTION_ANGIOGENIC_MARKERS |
| VECCHI_GASTRIC_CANCER_EARLY |
| WANG_CISPLATIN_RESPONSE_AND_XPC |
| WANG_RESPONSE_TO_GSK3_INHIBITOR_SB216763 |
| WANG_SMARCE1_TARGETS |
| WELCSH_BRCA1_TARGETS |
| WEST_ADRENOCORTICAL_TUMOR |
| YANG_BCL3_TARGETS |
| YOSHIMURA_MAPK8_TARGETS |
| ZHANG_BREAST_CANCER_PROGENITORS |
| ZHOU_INFLAMMATORY_RESPONSE_FIMA |
| ZHOU_INFLAMMATORY_RESPONSE_LIVE |
| ZHU_CMV_ALL |

#### Supplementary Table 3

The number of benchmark signatures (out of 192) in which the first method has significantly higher (two-sided Mann-Whitney U p <= 0.05) chemical diversity compared to the second method.

| | PACOS (enhance) | PACOS (repress) | PACOS + Tool (enhance) | PACOS + Tool (repress) |
|---|---|---|---|---|
| QUIZ-C vs MODZ\|CD | 143 | 144 | 146 | 151 |
| QUIZ-C vs L1000CDS2 | 125 | 127 | 103 | 108 |
| L1000CDS2 vs QUIZ-C | 45 | 40 | 55 | 64 |

#### Supplementary Table 4

| Perturbation | Drug description/MoA | Cell type | PACOS + Tool combined score (Repress) | AUROC | AUROC emp. pval | VGD stage | References |
|---|---|---|---|---|---|---|---|
| betamethasone | Corticosteroid | SKB | 0.655199891 | 0.8125 | 0.0025 | Early | 7, 8 |
| tamoxifen | | CORL23 | 0.643488296 | 0.679338843 | 0.0875 | Early | |
| estriol | | ASC | 0.607079937 | 0.640532757 | 0.1325 | Early | |
| thalidomide | | A673 | 0.601328083 | 0.618666667 | 0.175 | Early | |
| 7-nitroindazole | | RMGI | 0.590180388 | 0.549677419 | 0.36 | Early | |
| splitomycin | SIRT1 inhibitor | NEU | 0.573809314 | 0.697330791 | 0.015 | Early | 20 |
| AG-490 | EGFR tyrosine kinase inhibitor | SKB | 0.491281184 | 0.8125 | 0.0025 | Early | 12-15 |
| GW-7647 | | ASC | 0.485314527 | 0.640532757 | 0.1325 | Early | |
| formoterol | Long-acting beta agonist | NEU | 0.474404473 | 0.697330791 | 0.015 | Early | |
| rosiglitazone | PPAR-$\gamma$ agonist | NCIH2073 | 0.472365851 | 0.698780488 | 0.05 | Early | 1 |
| loperamide | Opioid-receptor agonist | HEC108 | 0.471989585 | 0.763841808 | 0.0125 | Early | |
| GW-7647 | PPAR-$\alpha$ agonist | SKB | 0.47185258 | 0.8125 | 0.0025 | Early | 3, 4 |
| nitrendipine | Calcium channel blocker | HEC108 | 0.466840953 | 0.763841808 | 0.0125 | Early | 10, 11 |
| GW-843682X | | RMGI | 0.466694863 | 0.549677419 | 0.36 | Early | |
| capsaicin | | NCIH596 | 0.463770687 | 0.588996764 | 0.2725 | Early | |
| cyclosporin-a | | SW620 | 0.462206053 | 0.662395543 | 0.1075 | Early | |
| DG-041 | | HCC15 | 0.461541243 | 0.624605678 | 0.1775 | Early | |
| calcitriol | Active vitamin D metabolite | SNUC4 | 0.452233834 | 0.746107784 | 0.025 | Early | 21 |
| dephostatin | | ASC | 0.449709245 | 0.640532757 | 0.1325 | Early | |
| sildenafil | | SKMEL1 | 0.448945213 | 0.576219512 | 0.2625 | Early | |
| SCH-58261 | Adenosine receptor A2A antagonist | SKB | 0.431723728 | 0.8125 | 0.0025 | Early | 22-24 |

| Perturbation | Drug description/MoA | Cell type | PACOS + Tool combined score (Repress) | AUROC | AUROC emp. pval | VGD stage | References |
|---|---|---|---|---|---|---|---|
| estriol | Steroid (Estrogen) | ASC | 0.58611692 | 0.73981002 | 0.0325 | Late | 5, 6 |
| splitomycin | | NEU | 0.5148704 | 0.62552381 | 0.1625 | Late | |
| dephostatin | Protein tyrosine phosphatase 1B inhibitor | ASC | 0.47709501 | 0.73981002 | 0.0325 | Late | 17, 18 |
| GW-7647 | PPAR-$\alpha$ agonist | ASC | 0.47022795 | 0.73981002 | 0.0325 | Late | 3,4 |
| RS-100329 | $\alpha$1-AR antagonist | ASC | 0.43838991 | 0.73981002 | 0.0325 | Late | 2 |
| sunitinib | Multitarget RTK inhibitor | ASC | 0.42459856 | 0.73981002 | 0.0325 | Late | 9 |
| formoterol | | NEU | 0.41836029 | 0.62552381 | 0.1625 | Late | |
| sulpiride | Dopamine/serotonin receptor antagonist | ASC | 0.41523462 | 0.73981002 | 0.0325 | Late | |
| paroxetine | Selective serotonin reuptake inhibitor | ASC | 0.40442384 | 0.73981002 | 0.0325 | Late | |
| gefitinib | EGFR tyrosine kinase inhibitor | ASC | 0.40350811 | 0.73981002 | 0.0325 | Late | 12-15 |
| pravastatin | HMG CoA reductase inhibitor | ASC | 0.40282203 | 0.73981002 | 0.0325 | Late | 16 |
| gefitinib | | NEU | 0.39579329 | 0.62552381 | 0.1625 | Late | |
| tozasertib | | NEU | 0.39447828 | 0.62552381 | 0.1625 | Late | |
| SB-203580 | p38 kinase inhibitor | ASC | 0.39028988 | 0.73981002 | 0.0325 | Late | 19 |
